# snPATHO-seq: unlocking the pathology archives

**DOI:** 10.1101/2023.12.07.570700

**Authors:** Taopeng Wang, Kate Harvey, Javier Escudero Morlanes, Beata Kiedik, Ghamdan Al-Eryani, Alissa Greenwald, Nikolaos Kalavros, Felipe Segato Dezem, Yuling Ma, Yered H. Pita-Juarez, Kellie Wise, Cyril Degletagne, Anna Elz, Azi Hadadianpour, Jack Johanneson, Fiona Pakiam, Heeju Ryu, Evan W. Newell, Laurie Tonon, Andrew Kohlway, Tingsheng Drennon, Jawad Abousoud, Ryan Stott, Paul Lund, Jens Durruthy, Andres F Vallejo, Dominik Kaczorowski, Joanna Warren, Lisa M. Butler, Sandra O’Toole, Jasmine Plummer, Ioannis S Vlachos, Joakim Lundeberg, Alexander Swarbrick, Luciano Martelotto

## Abstract

Formalin-fixed paraffin-embedded (FFPE) samples are valuable but underutilized in single-cell omics research due to their low DNA and RNA quality. In this study, leveraging recent single-cell genomic technology advances, we introduce a versatile method to derive high-quality single-nucleus transcriptomic data from FFPE samples.

## Main text

Clinical samples are routinely preserved as formalin-fixed paraffin-embedded (FFPE) tissue of proven utility in histology and bulk DNA or RNA studies^1,2^. However, exploration of single-cell genomics methods enabling profiling FFPE samples is nascent^2–7^.

Single-cell/nucleus RNA sequencing (scRNA/snRNA-seq) requires extensive amplification of the captured RNA for next-generation sequencing. FFPE samples, with their low RNA integrity and pronounced fragmentation, are not suitable for standard sc/snRNA-seq assays requiring intact transcripts. To address this challenge, we adopted a novel strategy (i.e. Flex by 10x Genomics) using RNA-targeting probes that latch onto smaller sections of the RNA molecules, thus reducing the requirements for RNA integrity^4^. Initial testing of this probe-based technology on human peripheral blood mononuclear cell (PBMC) samples confirmed its comparable performance to 10X Genomics’ conventional 3’ gene expression assays in terms of unique molecular identifiers (UMIs), gene detection per cell barcode, cell type representation, and marker gene expression (Extended Data Fig.1a-e). Moreover, the detected cell type signatures mirrored published PBMC data^8^ (Extended Data Fig.1f,g).

We then developed a novel snRNA-seq workflow, snPATHO-seq, tailored for FFPE tissue samples^3^. In brief, it involves an optimized versatile procedure for nuclei extraction from FFPE sections (or punches). The nuclei are then hybridized with Flex probes, followed by a PCR cycle-optimized library preparation and sequencing (Fig.1a). We applied this workflow to a wide range of clinical cancer and normal tissue samples including brain, breast, intestine, lung, liver, kidney and female reproductive organs (Fig.1a). We detected a median of 533 to 12496 UMIs and 388 to 4000 genes per nucleus across different tissue types (Extended Data Fig.2a,b). Furthermore, we successfully identified tissue-specific cell types in line with the tissue histology (Extended Data Fig.3).

To assess the performance of snPATHO-seq on FFPE against other snRNA-seq workflows, specifically 3’ and Flex gene expression assays on snap-frozen samples, we selected three breast cancer specimens with both FFPE and snap-frozen material available (Fig.1a, Supplementary Table S1). During the preparation of our manuscript, 10X Genomics introduced the scFFPE protocol using the same Flex chemistry^4^ (Fig.1a). While its performance matched snPATHO-seq across multiple tissues (Extended Data Fig.2, 3a-g, i), scFFPE was ineffective for our selected three breast cancer samples (Extended Data Fig.2a-c). Our initial attempts to extract whole cells from these breast cancer samples with Miltenyi’s FFPE Tissue Dissociation kit yielded mixed-quality suspensions (Extended Data Fig.4). This inconsistency led us to favor nuclear preparations for snPATHO-seq, which proved to be more reliable. However, given 10x’s previous success with scFFPE on breast FFPE tissue^4^, the exact reason for this discrepancy remains unclear. Due to sample constraints, we were unable to retest on the same FFPE blocks and therefore opted to exclude the scFFPE data from further breast cancer sample analysis in this study.

When applied to the three breast cancer samples, all three workflows, snPATHO-seq (FFPE), 3’ (frozen), and Flex (frozen), successfully identified cell types associated with both the cancerous region and its adjacent normal microenvironment (Fig.1c, Extended Data Fig.5b, Extended Data Fig.6b). For primary breast cancer sample 4066 and liver metastasis 4411, snPATHO-seq on FFPE and the two snap-frozen workflows produced comparable data (Fig.1d-f, Extended Data Fig.5c,d). Additionally, the cell type labels could be effectively transferred to Visium spatial transcriptomics data showing high consistency with pathology annotation (Extended Data Fig.7) and between techniques (Extended Data Fig.5e, Extended Data Fig.8).

For the liver metastasis sample 4399, fewer liver tissue-resident cell types were identified in snap-frozen workflows (Extended Data Fig.6c,d). Since these cell types were detectable by both 3’ and Flex assays in liver metastasis 4411, the inconsistency likely stems from sampling bias rather than workflow performance differences. We were able to confirm the presence of adjacent normal liver tissue in the 4399 FFPE sample through hematoxylin and eosin (H&E) staining (Extended Data Fig.6a). However, the snap-frozen sample from the same patient lacked Optimal Cutting Temperature (OCT) preservation, precluding histological assessment. This bias can influence downstream data interpretation. For example, when inferring cellular composition from FFPE Visium data using sample-matched snRNA-seq signatures, missing cell type signatures like hepatocytes and liver sinusoidal endothelial cells (LSECs) in the 3’ or Flex references might overemphasize the predicted weight of other cell types (Fig.1g, Extended Data Fig.6e). In comparison, snPATHO-seq offers snRNA-seq analysis from tissue scrolls informed by an adjacent H&E image, ensuring accurate tissue feature sampling.

Besides cell type annotation, snPATHO-seq also allows the detection of cancer heterogeneity. We discerned gene co-expression modules via non-negative matrix factorization (NMF) analysis^9^ using cancer nuclei and Visium spots with a dominant cancer epithelial signature from snRNA-seq and Visium data respectively. We defined 67 robust NMF programs grouped into 12 clusters (Fig.2a,b, Supplementary Table S2-4). These clusters span two categories: 1) recurrent cell states from diverse samples and workflows (5 programs); 2) tumour-specific patterns yet consistently seen across snRNA-seq workflows underscoring consistency across workflows (7 programs) (Fig.2a). Cross-sample clusters displayed gene enrichment resembling published pan-cancer metaprograms^9^ and utilization by both malignant and non-malignant cell types echoing previous literature^9,10^ (Fig.2c, Extended Data Fig.9). Conversely, sample-specific NMF clusters showcased the heterogeneity between cancer. For example, we identified a calcium signalling related NMF cluster (cluster 6) in sample 4066 that includes *ITPR2*, *CADPS2*, *CAPN13*, *NEATC4* and *ADRGV1* (Supplementary Table S4), all pivotal to calcium regulation and signalling. The cancer cell population defined by this gene module exhibited upregulation of the Reactome calcium pathway (Extended Data Fig.10). Mapping this calcium related module to sample 4066 Visium data revealed a spatial distribution pattern specific to ductal carcinoma *in situ* (DCIS) (Fig.2d), a condition often tied to calcification^11^. Further examination with a larger clinical cohort exhibiting similar tissue features is warranted to validate the significance of this calcium gene module.

**Fig 1:**
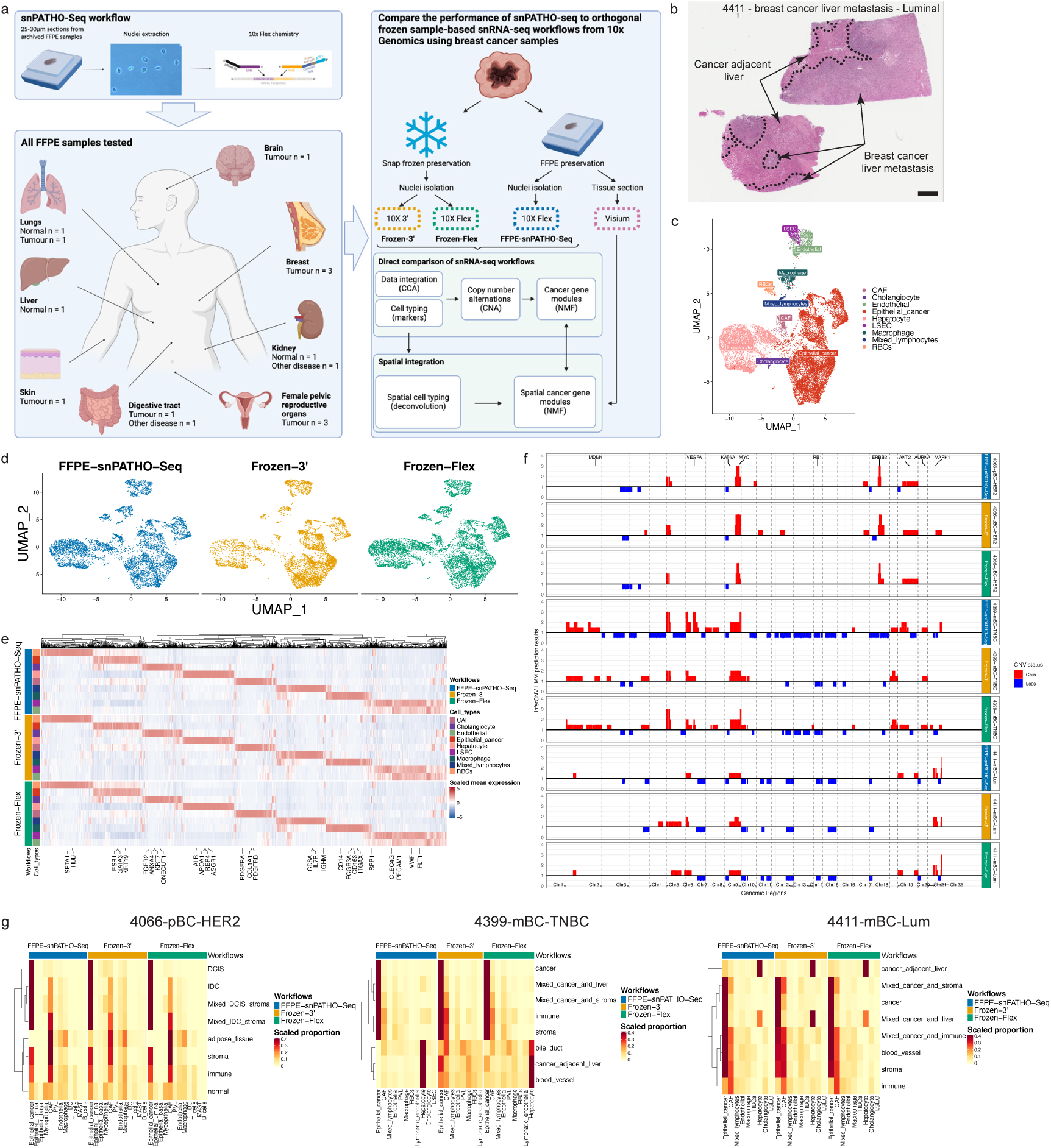
snPATHO-seq workflow and performance against snRNA-seq workflows on snap-frozen samples. (**a**) snPATHO-seq workflow. (**b**) H&E image of breast cancer liver metastasis (sample 4411) highlighting the tumor-liver interface. Scale bar = 1 mm. (**c**) Cell type annotation of down sampled (25000 reads per nucleus for all datasets) and Seurat CCA integrated snRNA-seq data from sample 4411. (**d**) UMAP visualization of data according to snRNA-seq workflows. (**e**) Differential gene expression heatmap with selected markers. (**f**) CNV profiles of cancer nuclei across datasets. (**g**) Heatmaps showing predicted snRNA-seq cell fractions in Visium data from corresponding patient samples.**snPATHO-seq workflow and performance against snRNA-seq workflows on snap-frozen samples.** (**a**) snPATHO-seq workflow. (**b**) H&E image of breast cancer liver metastasis (sample 4411) highlighting the tumor-liver interface. Scale bar = 1 mm. (**c**) Cell type annotation of down sampled (25000 reads per nucleus for all datasets) and Seurat CCA integrated snRNA-seq data from sample 4411. (**d**) UMAP visualization of data according to snRNA-seq workflows. (**e**) Differential gene expression heatmap with selected markers. (**f**) CNV profiles of cancer nuclei across datasets. (**g**) Heatmaps showing predicted snRNA-seq cell fractions in Visium data from corresponding patient samples.

**Fig 2:**
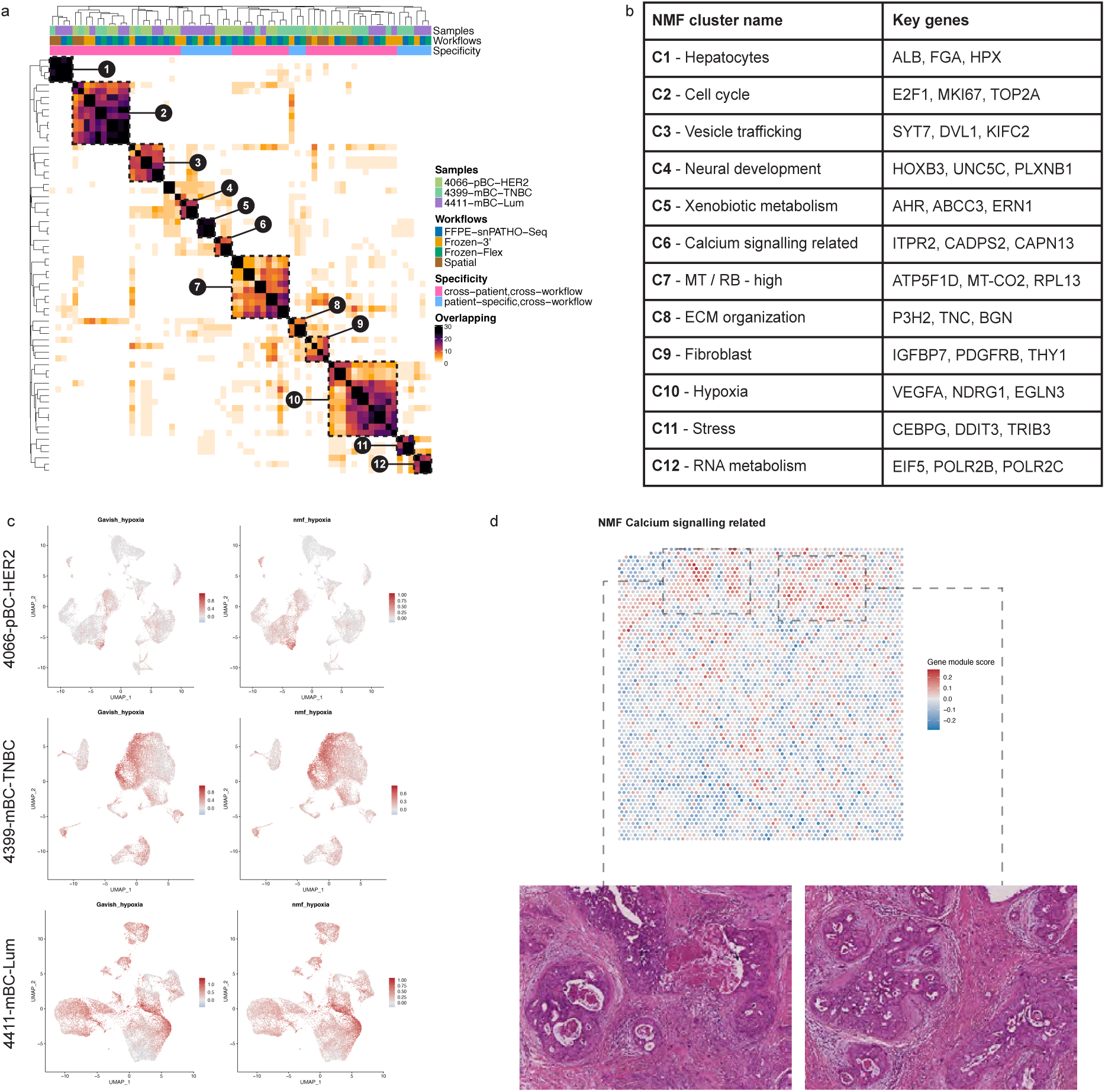
Characterization of cancer heterogeneity using snRNA-seq and Visium spatial transcriptomic. (**a**) Heatmap of overlapping genes in NMF programs from cancer epithelial nuclei in breast cancer datasets. (**b**) NMF cluster annotation. (**c**) UMAP comparison of hypoxia signature scores from this study and a pan-cancer metaprogram. (**d**) Spatial layout of calcium signalling NMF signature alongside tissue morphology.

In conclusion, we developed a robust snRNA-seq workflow tailored for FFPE tissue samples and demonstrated its versatility across diverse clinical specimens. snPATHO-seq enables high-quality and sensitive profiling of preserved tissue samples, unlocking the vast archives of FFPE tissues and facilitating comprehensive retrospective clinical research. Through the integration of H&E and spatial transcriptomics with snPATHO-seq, we bridged single-nucleus transcriptomics to their spatial morphology.

## Methods

### Patient material, ethics and consent for publication

For breast cancer samples, primary breast cancer sample 4066 was collected with written informed consent under the SVH 17/173 protocol with approval from St Vincent’s Hospital Ethics Committee. Metastatic breast cancer samples 4399 and 4411 were collected as autopsy samples under protocol X19-0496. Consent included the use of all de-identified patient data for publication. Collected tumour samples were macroscopically dissected. For FFPE tissue preservation, samples were fixed in 10% neutral buffered formalin for 24 hours at room temperature before being processed for paraffin embedding. For snap-frozen sample preservation, samples were diced into small chunks before snap freezing in liquid nitrogen and stored at -80℃.

For samples processed in the Cancer Research Centre of Lyon, the collection was provided by the 3D Onco platform (S. Ballesta) and Infirmerie Protestante de Lyon and was consented under the following ethical protocols (endometrium: 2022-05, colon: 2021-33, ovary: I-3422-01_MTA_IP_CLB) approved by the ethical review board of Centre Léon Bérard. The study was compliant with GDPR requirements and the CNIL (French National Commission for Computing and Liberties) law.

All other samples were obtained from commercial vendors as documented (Supplementary Table S1).

### Single-cell RNA profiling for frozen PBMC samples

Two frozen PBMC aliquots from the same healthy donor were thawed and divided equally for processing using the standard 10X 3’ and Flex assays, strictly following the respective recommended protocols. For the 3’ assay, the Chromium NextGEM Single-cell 3’ Reagent kit (v3.1, 10x Genomics) was utilized as per the Chromium Single Cell 3’ Reagent Kits User Guide (v3.1 Chemistry) (CG000204 - Rev D), targeting approximately 5000 cells. Meanwhile, for the Flex assay, PBMC cells underwent a brief fixation as directed by the Fixation of Cells & Nuclei for Chromium Fixed RNA Profiling guide (CG000478 - Rev A, 10x Genomics). Subsequently, the cells were profiled using the Chromium Fixed RNA Kit, Human Transcriptome (1000474, 10x Genomics), targeting a similar cell count as the 3’ assay.

### Single-nucleus RNA profiling for snap-frozen tissue samples

For snap-frozen samples, nuclei were isolated as previously described^12^ with modifications. Frozen samples were first thawed and finely minced. The fragments were subsequently homogenized using 1x Nuclei Ez lysis buffer (NUC-101, Sigma-Aldrich) enriched with 1U/μL RNAse inhibitor (RiboLock RNAse Inhibitor, EO0382, Thermo Fisher Scientific) and chilled for 5 minutes on ice. The separated nuclei were filtered through a 70μm mesh (pluriStrainer 70μm, 43-50070-51, pluriSelect) and rinsed twice with 1x PBS augmented with 1% BSA (MACS® BSA Stock Solution, 130-091-376, Miltenyi) and once with a 0.5x PBS + 0.02% BSA mix. They were subsequently resuspended in the 0.5x PBS + 0.02% BSA solution and re-filtered using a 40μm mesh (pluriStrainer Mini 40 um, 43-10040-50, pluriSelect). The final nucleus count was determined using the LUNA-FX7 cell counter (AO/PI viability kit, F23011, Logos).

For the 3’ experiments, gene expression libraries were constructed using the Chromium NextGEM Single-cell 3’ Reagent kit (v3.1, 10x Genomics), adhering to the Chromium Single Cell 3’ Reagent Kits User Guide (v3.1 Chemistry) (CG000204 - Rev D), with a target of 10,000 nuclei per reaction.

For the Flex experiments, nuclei were first fixed as per the Fixation of Cells & Nuclei for Chromium Fixed RNA Profiling guide (CG000478 - Rev A, 10x Genomics). Gene expression libraries were then constructed using the Chromium Fixed RNA Kit, Human Transcriptome (1000474, 10x Genomics), based on the Chromium Fixed RNA Profiling user guide (CG000477 - RevB), targeting 10,000 nuclei for each reaction.

### Single-nucleus RNA profiling for FFPE tissue samples (snPATHO-seq)

Two tissue sections, each between 25-30 µm thick, were first washed thrice with 1mL of Xylene for 10 minutes to remove paraffin. Subsequently, they were rehydrated via a sequence of 1mL ethanol baths, each lasting 1 minute: two rounds in 100% ethanol, then in 70%, 50%, and finally 30% ethanol. Specifically for breast tissues, the initial xylene wash was conducted at 55°C. The sections were then briefly rinsed with 1 mL of RPMI1640 (Gibco).

Tissue disruption began physically using a pestle in 100μL of a digestion mix composed of 1mg/mL Liberase TM (5401119001, Roche), 1mg/mL Collagenase D (11088858001, Roche), and 1U/uL of RNAse inhibitor (RiboLock RNAse Inhibitor, EO0382, Thermo Fisher Scientific) in RPMI1640. This mix was then filled up to a 1 mL volume and subjected to digestion at 37°C for 45-60 minutes.

For nuclear extraction, the pre-digested tissue was treated with 1x Nuclei Ez lysis buffer (NUC-101, Sigma-Aldrich) that included 2% BSA. Disintegration further proceeded via pipetting using a P1000 pipette. The separated nuclei were filtered through a 70 μm mesh, double-rinsed with 1x PBS supplemented with 1% BSA, and once with a 0.5x PBS + 0.02% BSA blend. They were then resuspended in this blend and re-filtered via a 40 μm mesh (pluriStrainer Mini 40μm, 43-10040-50, pluriSelect). The concluding nuclear count was ascertained using the LUNA-FX7 cell counter (AO/PI viability kit, F23011, Logos).

Gene expression libraries were prepared using the Chromium Fixed RNA Kit, Human Transcriptome (1000474 or 1000475, 10x Genomics), in accordance with the user guide (Chromium Fixed RNA Profiling, CG000477 - RevB). We implemented additional optimizations in the pre-amplification and Indexing cycles tailored for nuclei derived from FFPE samples. Over 200,000 nuclei underwent a 20-hour hybridization with the BC01 probe set. This was followed by three washes using the Post-Hyb Buffer as recommended in the guide, with an additional wash step. Post-hybridization, nuclei were resuspended in the Post-Hyb Resuspension Buffer, counted and loaded onto Chip Q/Chromium X for capture, adhering to the guide’s procedures. A 9-cycle pre-amplification was conducted. Indexing PCR cycles followed the guide’s recommendations for PBMC and nuclei, but with two added cycles. For sample 4399, the capture targeted 5,000 nuclei, while for other samples, the target was set at 10,000 nuclei.

### Single-cell RNA profiling for FFPE tissue samples (scFFPE)

scFFPE workflow was conducted on the FFPE samples by the 10x Genomics and Fred Hutch Innovation Lab as outlined in Supplementary Table S1 following the demonstrated protocol (CG000632, RevB).

### Spatial transcriptomics on breast cancer samples

A 5μm-thick section was prepared from the FFPE blocks and processed using the Visium Spatial Gene Expression for FFPE Kit v2 (10x Genomics) according to the manufacturer’s instructions. Briefly, sections were H&E stained and imaged followed by probe hybridization and ligation.

### Illumina sequencing

The indexed libraries were sequenced using the Illumina NextSeq550 or NovaSeq 6000 systems. For sc-/sn-RNAseq libraries, the read format was configured as 28, 10, 10, and 90 for Read 1, i7, i5 and Read 2 sequences respectively. For Visium spatial transcriptomics libraries, the read format was configured as 28, 10, 10 and 50 for Read 1, i7, i5 and Read 2 sequences respectively. Libraries were sequenced to a depth of more than 10000 read pairs per cell/nuclei for sc-/sn-RNAseq experiments or more than 25,000 read pairs per spot for Visium experiments.

### FASTQ files processing

For data generated by the 10X Genomics, raw reads were de-multiplexed and aligned using Cellranger (v2023.0415.0). GRCh38 (build 2020-A, 10X Genomics) and Chromium Human Transcriptome Probe Set v1.0.1 were used as references.

For other single-cell/nucleus data, sequencing reads were de-multiplexed and aligned using Cellranger v7.0.1. GRCh38 (build 2020-A, 10X Genomics) was used as the reference for read mapping. For Flex related workflows, Chromium Human Transcriptome Probe Set v1.0 reference was used for probe read mapping.

For Visium data, reads were processed using Spaceranger v2.0.0 with GRCh38 (build 2020-A, 10X Genomics) and Visium Human Transcriptome Probe Set v2.0 as references.

For direct comparison of different single-cell/nucleus workflows, the PBMC data was downsampled to ∼30,000 reads per cell and the breast cancer snRNA-seq datasets were downsampled to ∼25,000 reads per nucleus. For 3’ data, FASTQ files were downsampled using Seqtk (v1.3)^13^. For Flex assay data, FASTQ files were downsampled during Cellranger analysis by specifying the targeted numbers of reads per cell/nuclei in the configuration file supplied to the “cellranger multi” function.

### Ambient background filtering for snRNA-seq datasets

Additional ambient reads filtering was conducted using Cellbender v0.2.0^14^. In total, 40000 droplets were used to estimate the levels of background. The cell numbers estimated using the EmptyDrops method^15^ implemented in the Cellranger were supplied as the expected numbers of cells for Cellbender processing.

### Low quality data filtering for single-cell/nucleus RNA sequencing datasets

The count matrices from snRNA-seq datasets were then processed using Seurat (v4.3.0.9002)^8^ in R (v4.1.1) unless otherwise specified. Low quality cells/nuclei were defined as cells/nuclei with less than 200 UMIs, over 8000 UMIs or over 10% mitochondrial gene products. Gene expression data was normalized using the “NormalizeData” function in the Seurat package. Doublets were identified using the DoubletFinder package (v2.0.3)^16^ and excluded from the downstream analysis.

### Dimensionality reduction, clustering and cell type annotation of sc/snRNA-seq data

For samples processed using more than one sc/sn-RNA workflow, the filtered counts were integrated at a per-sample level using the canonical correlation analysis (CCA) method as implemented in the Seurat package. The top 2000 highly variable genes in each dataset were used for the principal component analysis (PCA) and the top 2000 highly variable genes across datasets to be integrated were used to identify integration anchors.

For other datasets, PCA analysis was conducted using the top 2000 highly variable features.

For all datasets, Uniform Manifold Approximation and Projection (UMAP) processing was conducted using the top 30 principal components from the CCA or PCA analysis. Gene expression data was clustered using the top 30 principal components with the Louvain method as implemented in the Seurat package. Clusters were manually annotated based on the expression of canonical cell type markers. Re-integration or subclustering was also performed to separate cell lineages unable to be resolved by initial clustering. The most differentially expressed genes were derived using Student’s t-test with the “FindAllMarkers” function in the Seurat package and visualised using the ComplexHeatmap package (v2.10.0)^17^. For all datasets, the top 200 differentially expressed genes ranked by fold change for each cell type cluster were plotted. For breast cancer datasets, canonical cell type markers were annotated. For all other datasets, the top 5 differentially expressed genes from each cluster were annotated.

### Automatic cell type annotation using singleCellNet

The published PBMC 3’ dataset from Hao et al., 2021 was annotated using singleCellNet (v0.1.0)^18^ and PBMC cell type signatures from this study. A classifier was trained using the top 50 genes from each cell type for 3’ or Flex PBMC data respectively. All other parameters were kept the same. The cell type annotation results were then visualised using the ggplot2 package (v3.4.2) included in tidyverse (v2.0.0) in R.

### Copy number variation inference

The copy number alteration of cancer cells was inferred using inferCNV (v1.10.1)^19^ on a per sample basis. All stromal and immune cell types within each dataset were selected as the reference cell types. Red blood cells (RBCs) and clusters with mixed cell type signatures were excluded from the dataset. For breast cancer 4399 datasets, as liver resident cell types were mainly identified in the snPATHO-seq dataset, they were also excluded from the reference. This includes cholangiocytes, hepatocytes and LSECs. The analysis was then conducted on all normal and cancer epithelial cells with HMM prediction. Default parameters were kept except for the cut-off which was set to 0.1 according to the authors’ recommendation for 10x Chromium data.

For breast cancer datasets, the HMMi6 results for each gene were summarised across all cancer nuclei. The results were rounded up to the nearest multiple of 0.5. Any gene either not detected in the dataset or showed no CNV change was assigned with a value of 1. The results were summarised per dataset and plotted together using the ggplot2 package (v3.4.2) included in tidyverse (v2.0.0) in R.

### Spatial transcriptomics data processing

For Visium spatial transcriptomics data, tissue morphology was annotated at per spot level using the Loupe browser (v6.2.0, 10X Genomics). Spots underneath regions affected by processing artefacts were annotated as “exclude”. The count metrics were converted into STutility objects (v1.1.1)^20^ for downstream processing. Low quality spots annotated as “exclude” were removed from the datasets. The metrics were then normalised using the “NormalizeData” function from the Seurat package implemented by the STutility package.

### Inference of spatial transcriptomics data cellular composition using deconvolution

Deconvolution of spatial cellular composition was performed using the RCTD method implemented in the spacexr package (v2.2.0)^21^. Datasets generated using 3’, Flex and snPATHO-seq were used as single-cell type references respectively for the deconvolution of matching Visium data. Default parameters were used for the analysis except that the minimal number of cells required for each cell type was reduced to 3 to accommodate relatively low cell number in the current study.

Deconvolution results were then visualised as heatmaps using the ComplexHeatmap package or as spatial feature plots using the ggplot2 package implemented in tidyverse.

### Robust NMF program identification and downstream analysis

Robust NMF programs were derived using a published method^9^ with the NMF package (v0.26) installed in R (v4.2.3). NMF programs were first derived from cancer epithelial nuclei in snRNA-seq data or from Visium spots with over 50% cancer epithelial signatures through deconvolution. The NMF algorithm was run using ranks from 4 to 9 with 10 iterations at each rank. The derived NMF programs were then collapsed by similarity in gene composition to select robust NMF programs which were further clustered based on similarity. Robust NMF clusters with at least 3 programs were annotated according to their interception with published pan-cancer meta programs, selected gene markers and sample origin.

To evaluate the usage of robust NMF programs in the snRNA-seq and Visium datasets. Robust NMF programs belonging to the same cluster (modules) as defined above were selected. Any gene present at least twice across all NMF programs in each module was defined as the core gene. Gene module scores were then calculated using the core gene lists using the “AddModuleScore” function defined in a previous publication^22^ and implemented in the Seurat package.

For nucleus annotation using gene module scores, module scores of relevant gene modules were first calculated in the cancer epithelial nuclei population in the integrated snRNA-seq data for each sample. Any cancer nucleus with no gene module score passing 0.5 was not annotated. The rest of the nuclei were annotated based on the gene module with the maximum score.

To evaluate the top biological pathways upregulated in nuclei annotated using gene modules, differential gene expression analysis between the annotated nuclei and all other cancer epithelial nuclei was conducted using the Wilcoxon rank sum test in the “FindAllMarkers” function in the Seurat package. All genes were tested and ranked from highest to lowest based on the fold change. Pathway analysis was then conducted using the fgsea package (v1.20.0)^23^ with Reactome gene set reference downloaded from the MsigDB. Any gene set with less than 15 or over 200 genes was excluded from the analysis. The analysis was conducted with a permutation of 10,000. Results of top enriched pathways were visualised using the ggplot2 package in tidyverse.

## Supporting information

biorender Publication License Oct-03-2023

scripts

## Data availability

Raw and processed sequencing data is being uploaded to Gene Expression Omnibus (GEO). Accession code will be provided once the upload is complete.

## Abbreviations

FFPE: formalin-fixed paraffin-embedded
scRNA: seq single-cell RNA-sequencing
snRNA: seq single-nucleus RNA-sequencing
PBMC: peripheral blood mononuclear cell
UMI: unique molecular identifier
H&E: hematoxylin and eosin
OCT: Optimal Cutting Temperature
LSEC: liver sinusoidal endothelial cell
NMF: non-negative matrix factorization

## Acknowledgements

This work is supported by the National Health and Medical Research Council (NHMRC) Ideas Grant (APP2004774), the Commonwealth Standard Grant Agreement (4-F26M8TZ) and Institut PLAsCAN and “La ligue contre le cancer – Comité de l’Ain” for samples processed in the Cancer Research Center of Lyon. The funders had no role in study design, data collection and analysis, decision to publish or preparation of the manuscript.

We thank Jens Durruthy Durruthy, Daniel “Telstra” Dlugolensky, Andrew Kohlway from 10x Genomics and the team at Millennium Sciences for the technical support and for providing access to the 10x Genomics Flex kit.

We thank Naiara Bediaga and Monika Mohenska from the University of Adelaide for facilitating data organisation.

We thank Garvan Histology for fast turnover tissue processing.

---

**Extended Data Fig.1:**
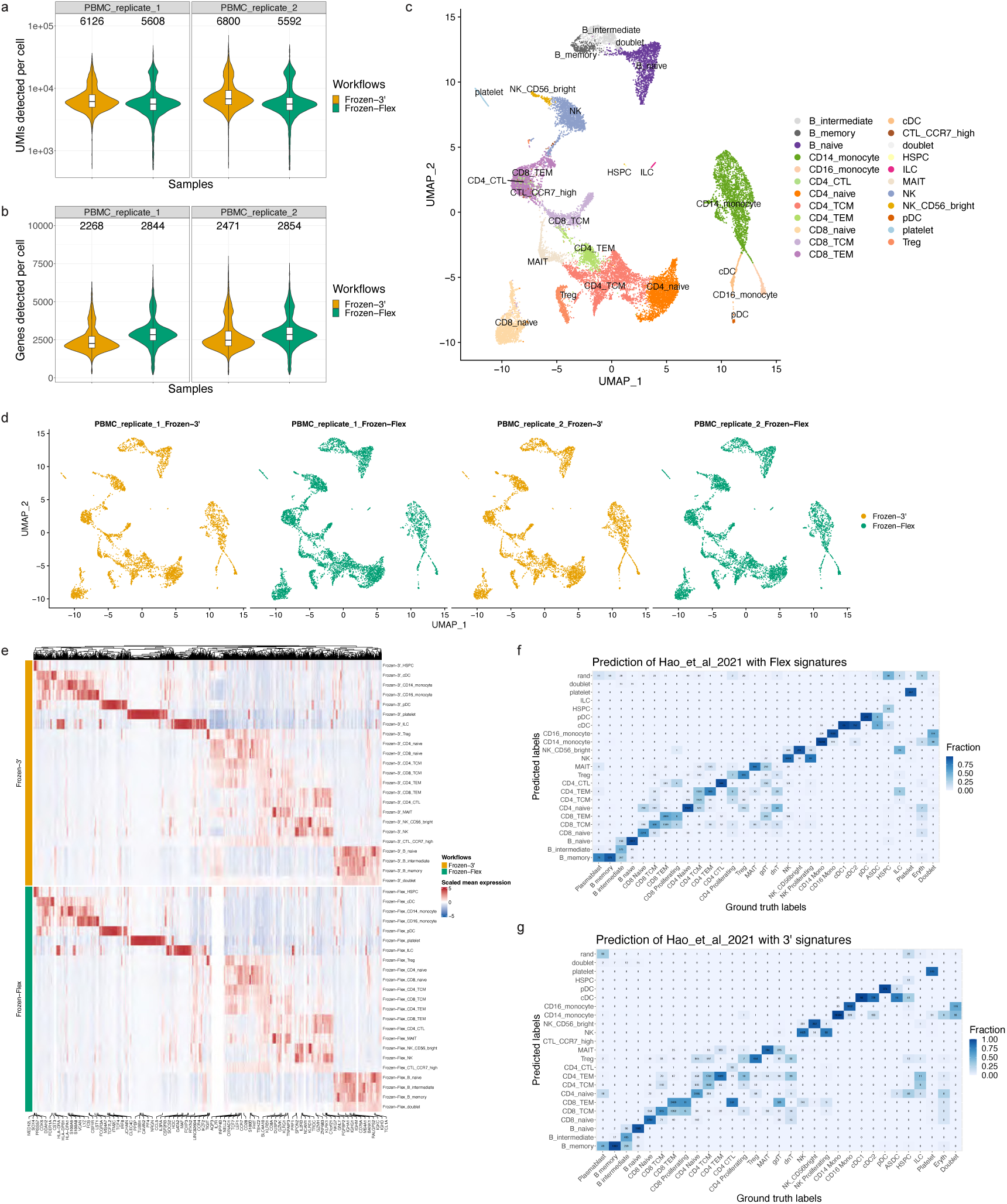
Comparison of the performance of the Flex chemistry to 3’ chemistry using PBMC samples. (**a & b**) The numbers of UMI and genes detected per cell by the Flex and 3’ assays across two PBMC replicates from the same donor. The numbers correspond to medians of UMIs and genes detected per cell in each dataset. (**c**) Cell type annotation of integrated Flex and 3’ data. (**d**) UMAP representation of the integrated data split by sample and processing method. (**e**) Heatmap of top differentially expressed genes between cell types. Top 5 differentially expressed genes by fold change for each cell type were plotted. (**f & g**) Annotation of the 3’ dataset published in Hao et al., 2021 using cell type signatures obtained from the Flex or 3’ assays in this study.

**Extended Data Fig.2:**
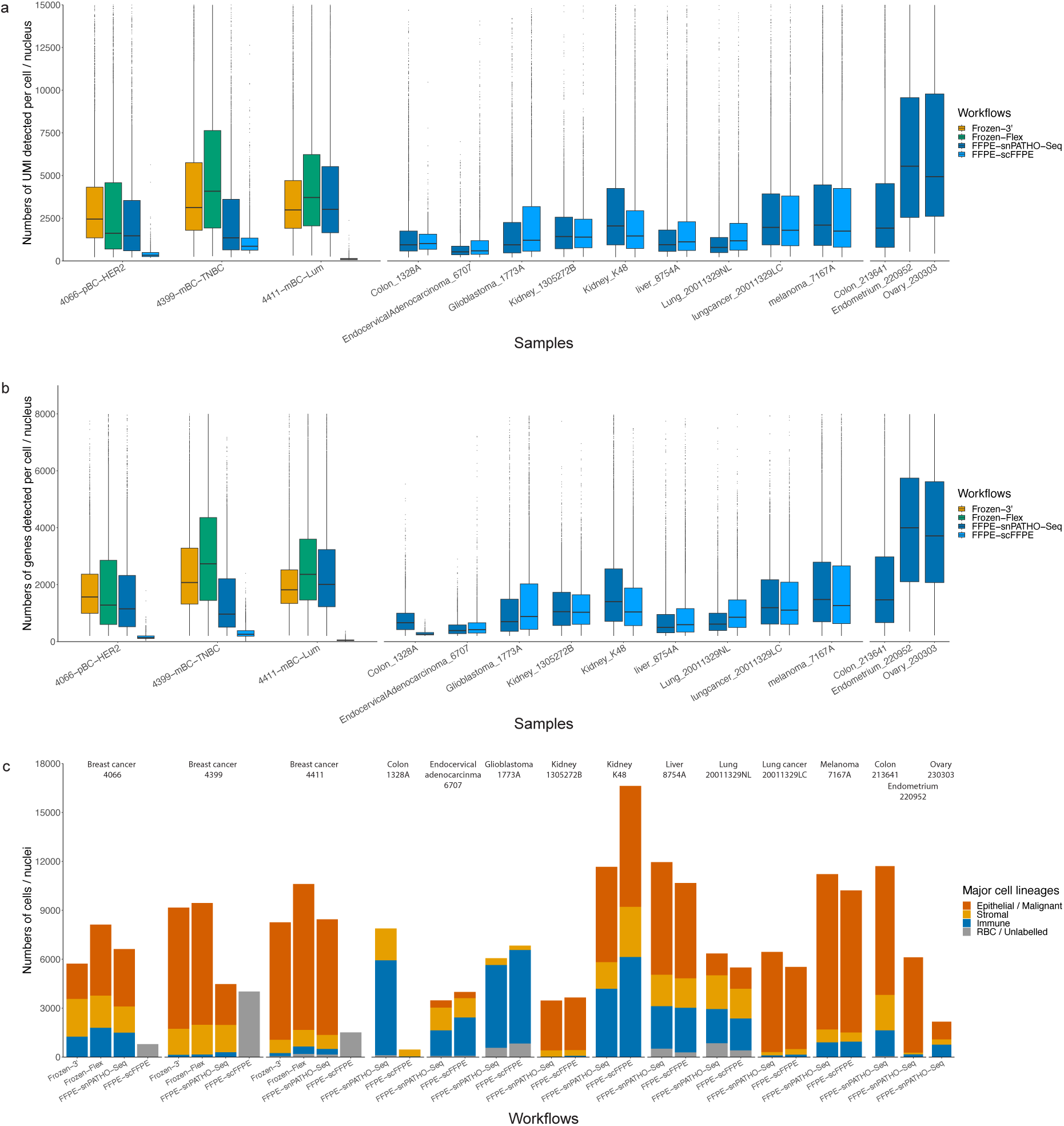
Quality control metrics and cellular composition of all snRNA-seq datasets generated. (**a & b**) The numbers of UMI and genes detected per nucleus/cell using different snRNA-seq/scFFPE workflows. (**c**) Numbers of nuclei or cells detected in each dataset.

**Extended Data Fig.3:**
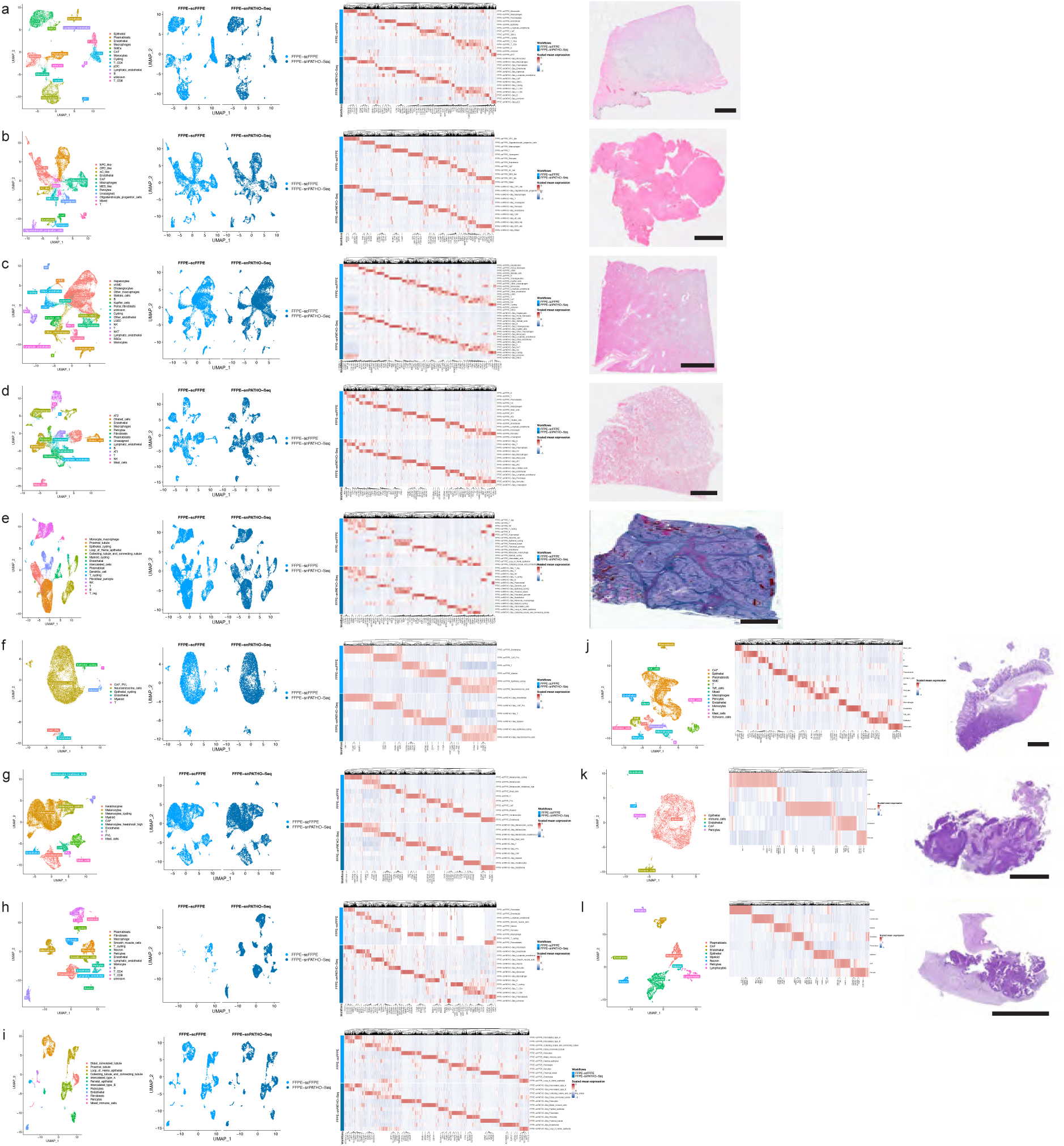
Overview of snPATHO-seq and scFFPE data collected from FFPE tissue samples other than breast cancer. (**a-e**) (left to right) Cell type annotation, data integration, top differentially expressed markers and H&E image of Endocervical_Adenocarcinoma_6707 (**a**), Glioblastoma_1773A (**b**), liver_8754A (**c**) and lung_20011329NL (**d**), Kidney_K48 (**e**). (**f-i**) (left to right) Cell type annotation, data integration and top differentially expressed markers of lung_cancer_20011329LC (**f**), melanoma_7167A (**g**), Colon_1328A (**h**) and Kidney_1305272B (**i**). (**j-l**) (left to right) Cell type annotation, top differentially expressed markers and H&E image of Colon_213641(**j**), Endometrium_220952 (**k**) and Ovary_230303 (**l**). Scale bars = 2mm.

**Extended Data Fig.4:**
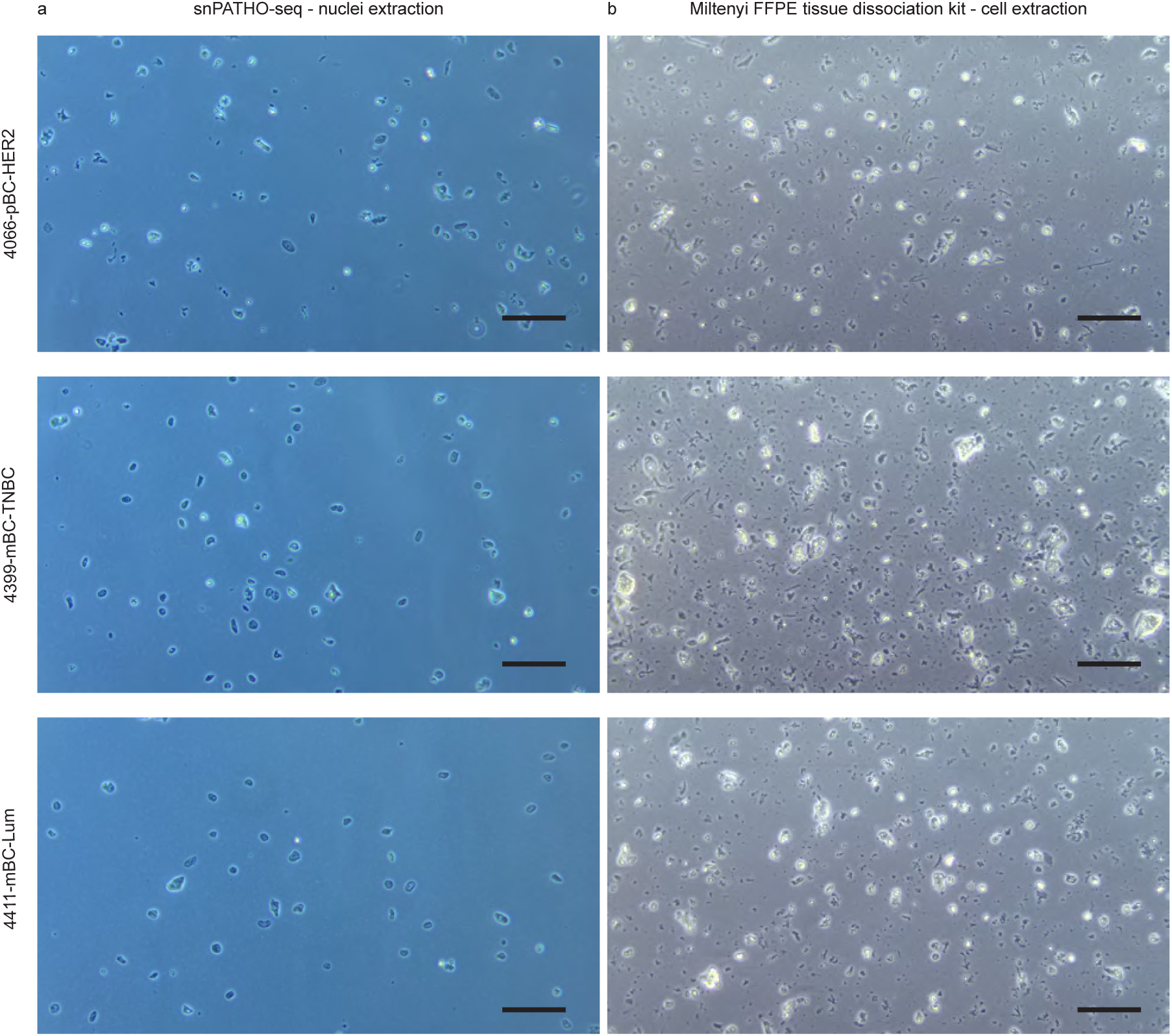
Microscopic images of isolated samples using Miltenyi’s FFPE tissue dissociation kit or snPATHO-seq workflow. **(a)** Images of extracted nuclei prepared using snPATHO-seq workflow. **(b)** Images of extracted breast cancer cells prepared using Miltenyi’s FFPE tissue dissociation kit. Scale bars = 100p.m.

**Extended Data Fig.5:**
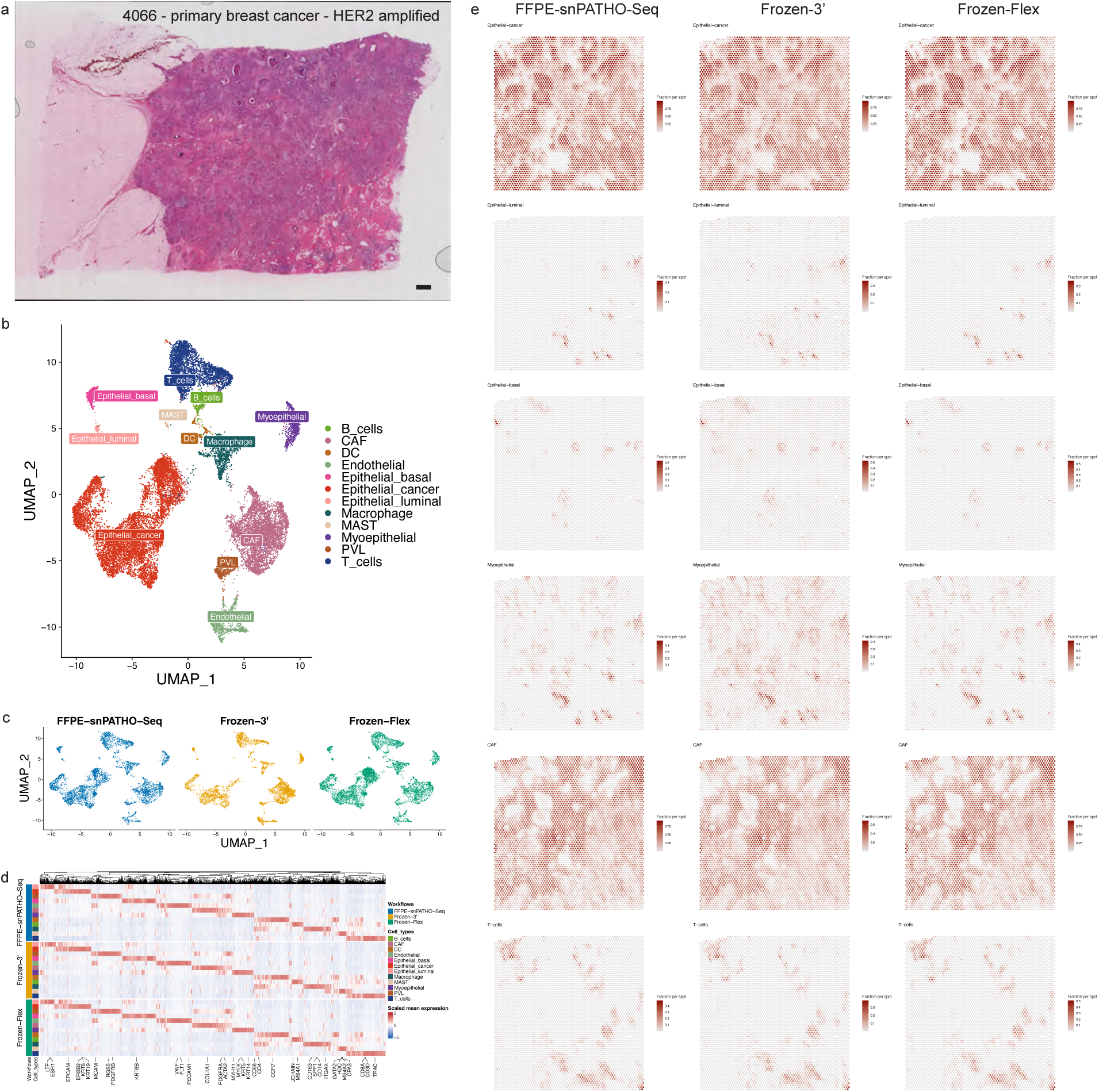
Compare the performance of all snRNA-seq workflows using 4066. (**a**) H&E image of 4066. Scale bar = 1mm. (**b**) Cell type annotation of integrated dataset generated using different snRNA-seq workflows using 4066. (**c**) UMAP representation of the integrated data split by snRNA-seq workflows. (**d**) Heatmap of top differentially expressed genes. (**e**) Spatial distribution of deconvolution results of selected cell type in 4066 Visium data.

**Extended Data Fig.6:**
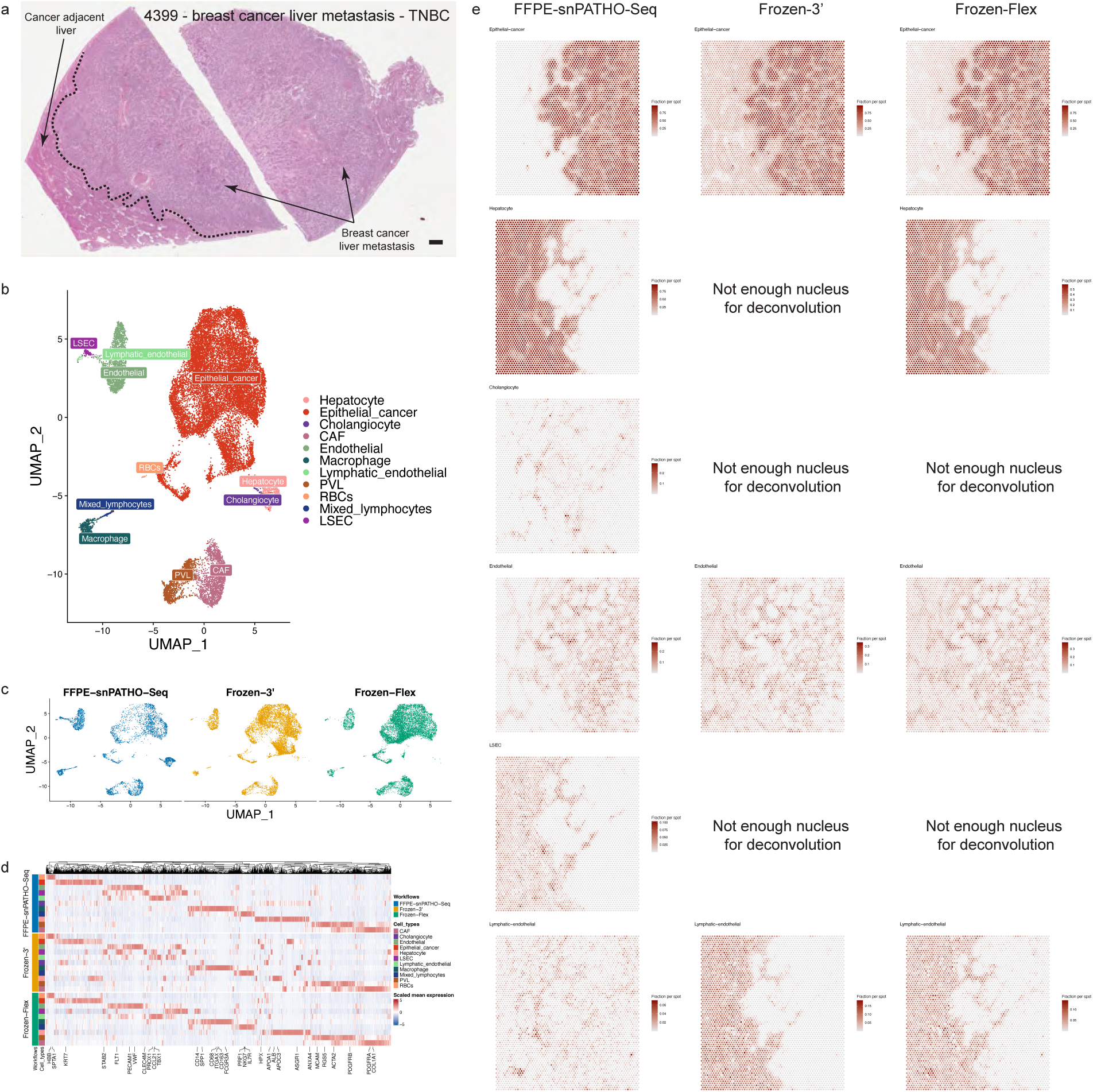
Compare the performance of all snRNA-seq workflows using 4399. (**a**) H&E image of 4399. Scale bar = 1mm. (**b**) Cell type annotation of integrated dataset generated using different snRNA-seq workflows using 4399. (**c**) UMAP representation of the integrated data split by snRNA-seq workflows. (**d**) Heatmap of top differentially expressed genes. (**e**) Spatial distribution of deconvolution results of selected cell type in 4399 Visium data.

**Extended Data Fig.7:**
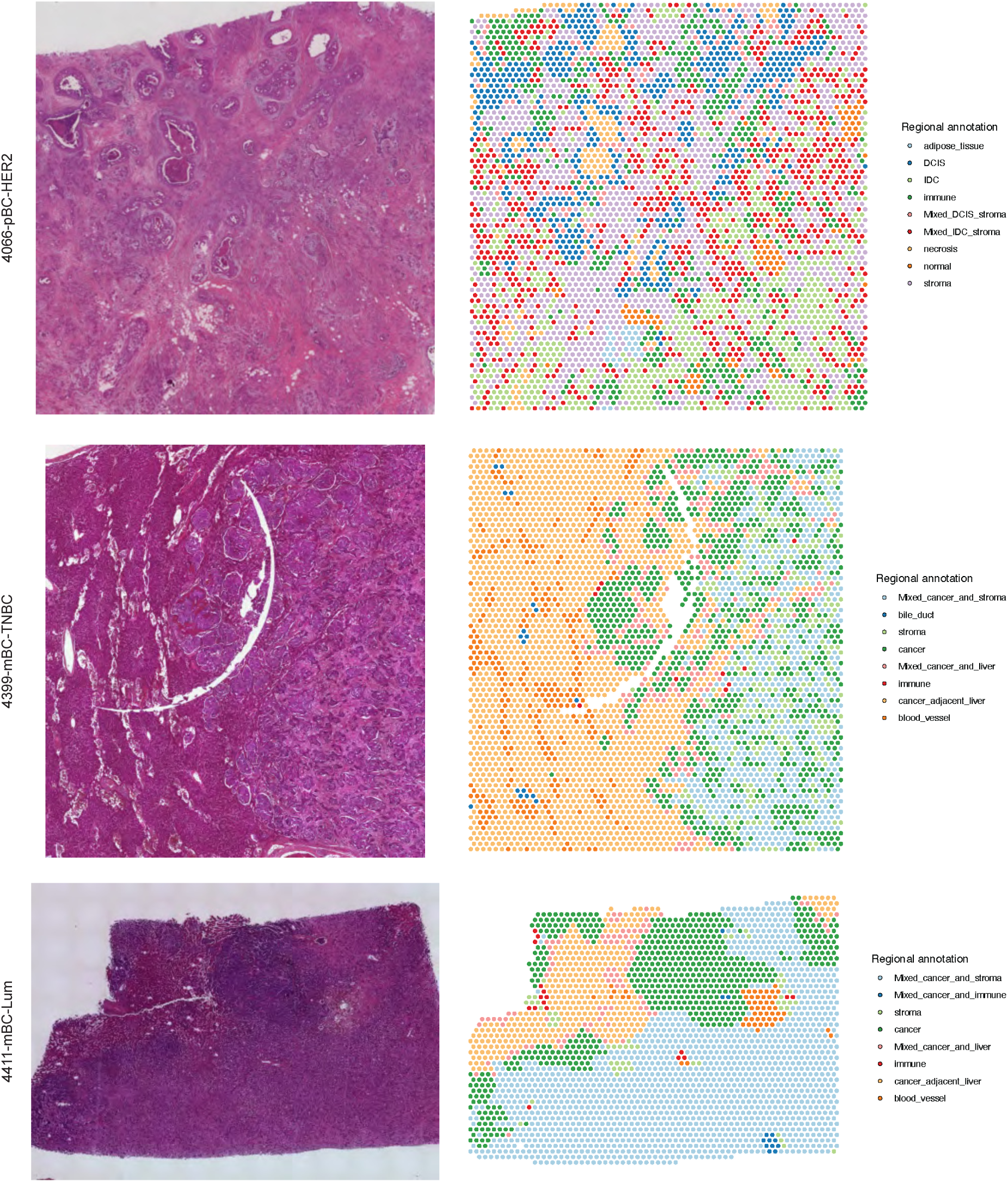
Pathology annotation of the tissue regions targeted by Visium experiments. H&E images of the tissue region targeted by Visium assay and per spot pathology annotation.

**Extended Data Fig.8:**
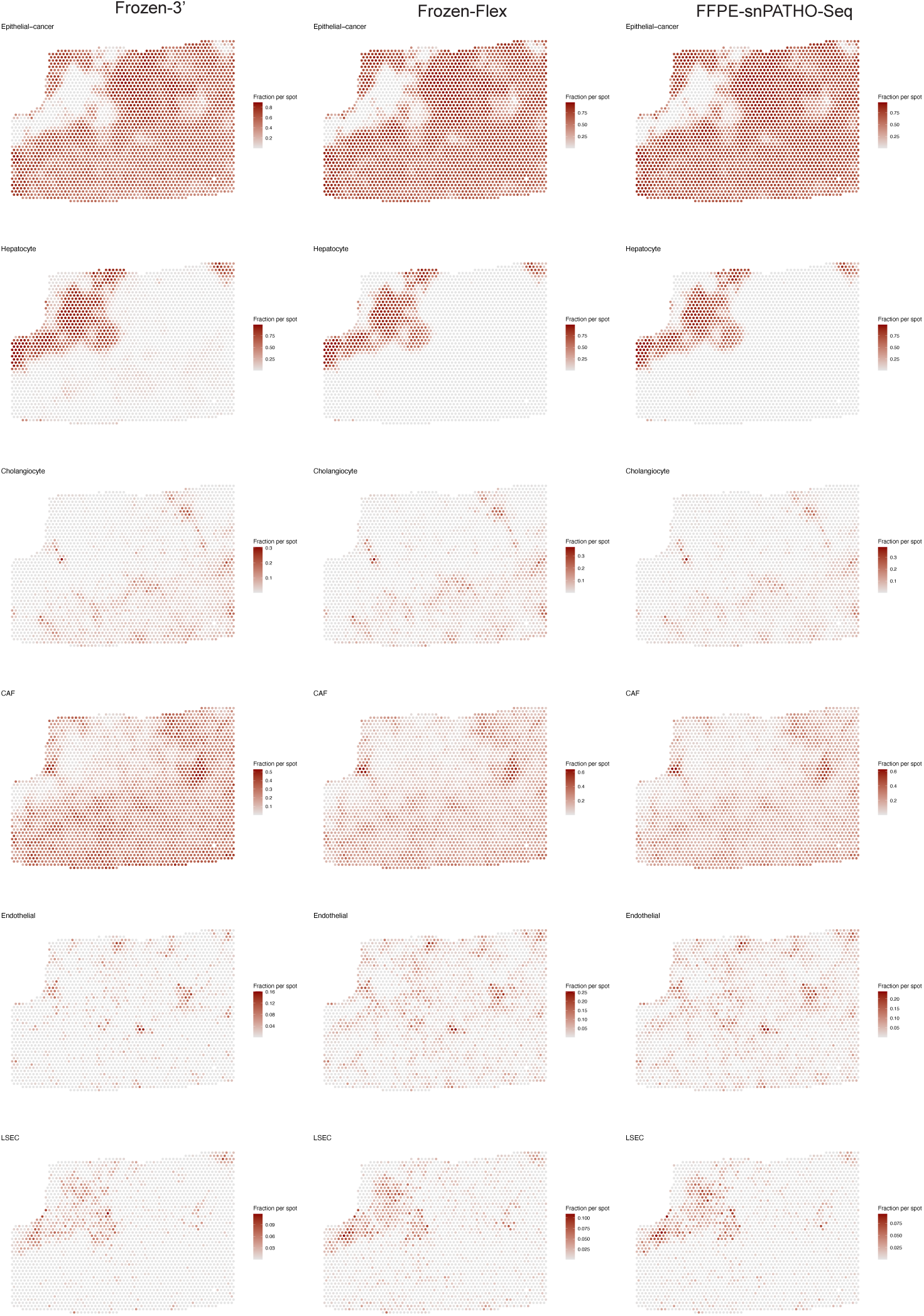
Spatial distribution of predicted cell proportions of selected cell type in sample 4411 Visium spatial transcriptomic data. Deconvolution was conducted on the FFPE Visium data using cell type signatures obtained from 3’, Flex and snPATHO-Seq workflows respectively from breast cancer liver metastasis sample 4411.

**Extended Data Fig.9:**
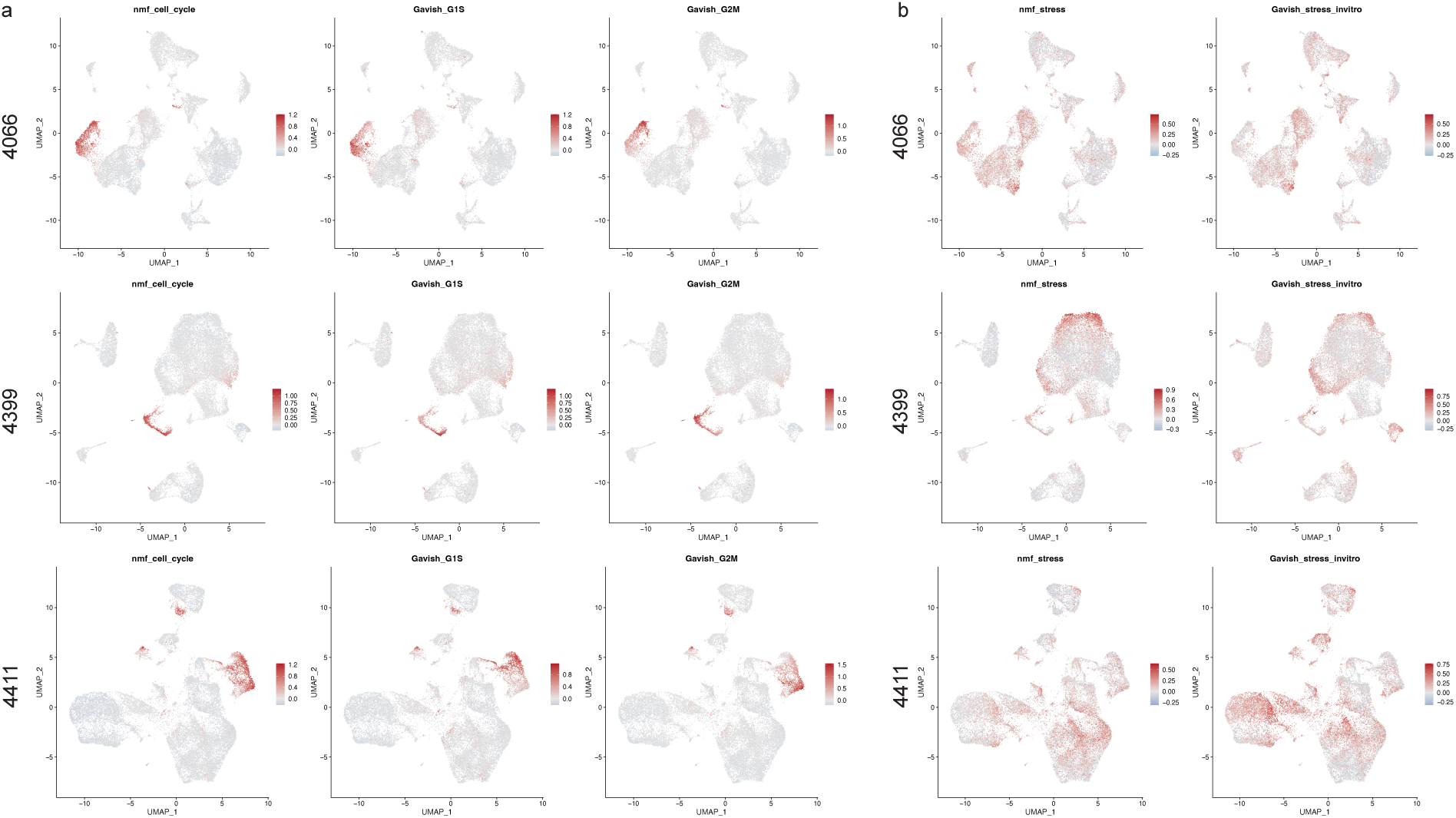
Gene module usage in different cancer populations. UMAP of gene module scores calculated using cell cycle and stress signatures derived from the current study and published pan-cancer metaprograms. Genes present in at least two robust NMF programs within the same NMF clusters (cell cycle or stress) were defined as the core genes and used for gene module score calculation.

**Extended Data Fig.10:**
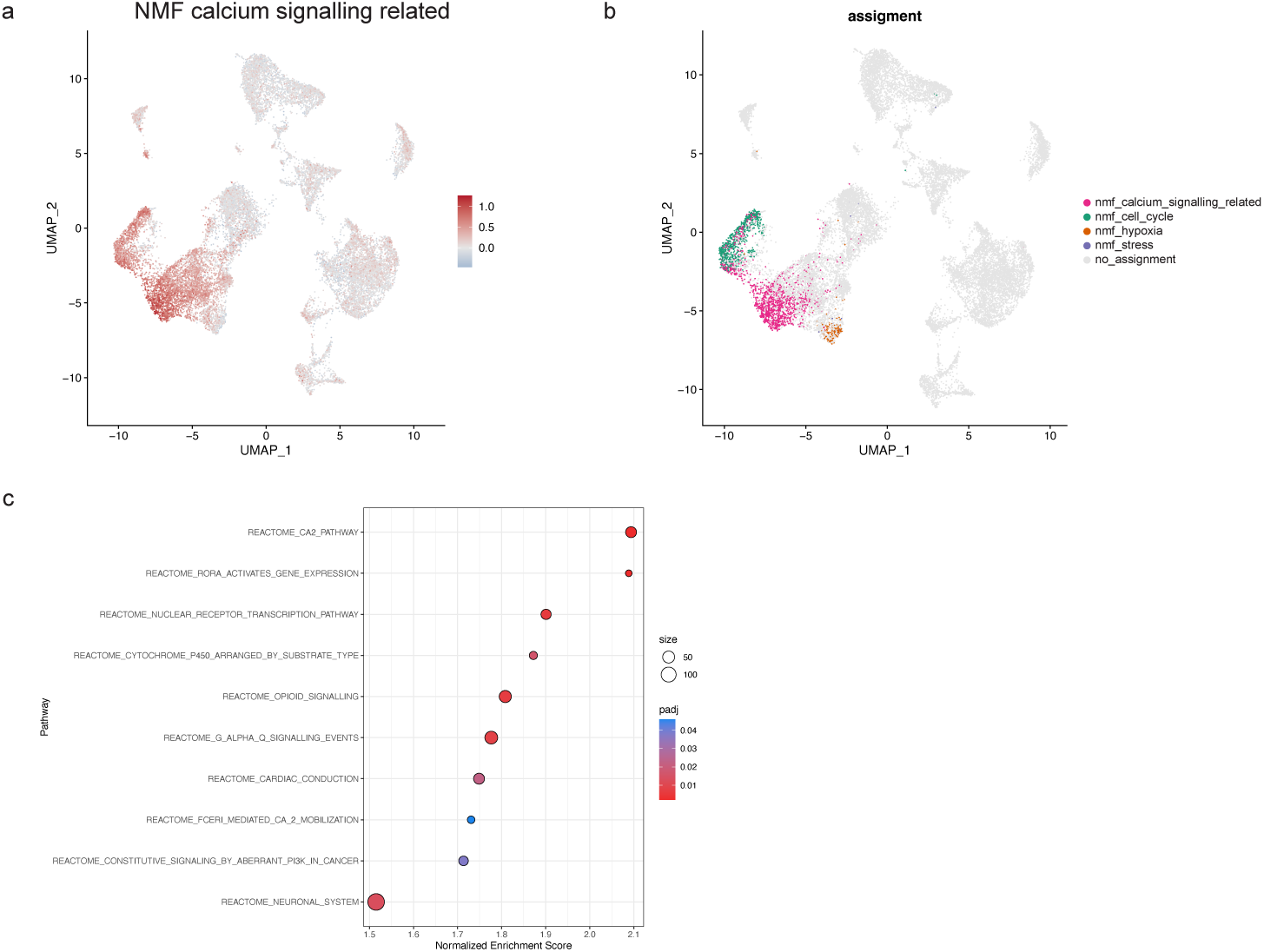
Identification of a DCIS related calcium signalling module in 4066. (**a**) UMAP of calcium signalling NMF cluster gene module score across cancer and non-cancer nuclei in sample 4066. (**b**) Annotation of snRNA-seq data using calculated gene module score. Gene module scores for calcium signalling, cell cycle, hypoxia and stress NMF signature characterised in this study were calculated for the cancer nuclei in sample 4066. Cancer nuclei were filtered so that at least one signature has a gene module score above 0.5 and annotated according to the signature with the highest score. The other cancer nuclei and tumour microenvironment nuclei were kept as unlabelled. (**c**) Top differentially regulated pathways in the cancer population annotated with calcium signalling NMF signature in sample 4066.

