## Supplementary material for "snPATHO-seq: unlocking the pathology archives": scripts: 01_initial_processing.R

```
# aim ----
    # take cellbender outputs and turn them into organised seurat objects 
        # QC plots
        # basic cell filtering
        # doubletfinder filtering
        # dimension reduction
        # clustering optimisation


# define environment ----
.libPaths(
    c(
        '/share/ScratchGeneral/tonwan/projects/snPATHO-Seq/renv/library/R-4.1/x86_64-conda-linux-gnu',
        .libPaths()
    )
)
library(Seurat)
library(tidyverse)
library(RColorBrewer)


# arguments ----
data_dir <- '/share/ScratchGeneral/tonwan/projects/snPATHO-Seq/data/external_data/10X_data_cellbender_filtered'

datasets <- list.dirs(data_dir, full.names = FALSE, recursive = FALSE)

out_data_dir <- '/share/ScratchGeneral/tonwan/projects/snPATHO-Seq/data/external_data/10X_data_processed'
dir.create(out_data_dir, recursive = T)

out_dir <- '/share/ScratchGeneral/tonwan/projects/snPATHO-Seq/results/snRNA_external_data/10X_data'
dir.create(out_dir, recursive = T)


########################################################################################################################
########################################################################################################################
########################################################################################################################
# loop through each dataset
for (ds in datasets) {
    if (file.exists(paste0(out_data_dir, '/', ds, '_clustered.rds'))) {
        next
    }


    # make a directory to store results
    working_dir <- paste0(out_dir, '/', ds)
    dir.create(working_dir, recursive = T)

    # load data and build an object
    cellbender_mtx <- paste0(data_dir, '/', ds, '/', 'cellbender_filtered_matrix_filtered.h5')
    counts <- Read10X_h5(cellbender_mtx, use.names = TRUE, unique.features = TRUE)
    obj <- CreateSeuratObject(counts, project = "10X_data", assay = "RNA")
    obj[["percent.mt"]] <- PercentageFeatureSet(obj, pattern = "^MT-")

    # populate metadata
    obj$dataset_id <- ds
    obj$processing_method <- unlist(lapply(str_split(obj$dataset_id, '_'), '[[', 3))
    sample_id_1 <- unlist(lapply(str_split(obj$dataset_id, '_'), '[[', 1))
    sample_id_2 <- unlist(lapply(str_split(obj$dataset_id, '_'), '[[', 2))
    obj$sample_id <- paste0(sample_id_1, '_', sample_id_2)

    obj <- subset(obj, subset = nFeature_RNA > 0)
    
    # simple plots to check QC
    pdf(file = paste0(working_dir, '/', 'QC.pdf'), width = 15, height = 5)
        print(
            VlnPlot(obj, features = c('nFeature_RNA', 'nCount_RNA', 'percent.mt'), group.by = 'sample_id',
                pt.size = 0)
        )
    dev.off()

    # minimal filtering
    filtered_obj <- obj %>% 
        subset(subset = nFeature_RNA > 200) %>% 
        subset(subset = nFeature_RNA < 8000) %>% 
        subset(subset = percent.mt < 10)


    # ----
    # use doubletfinder to exclude doublets
    db_removal_dir <- paste0(working_dir, '/', 'doublet_removal')
    dir.create(db_removal_dir, recursive = T)

    source('/share/ScratchGeneral/tonwan/projects/snPATHO-Seq/scripts/function_scripts/05_doublet_finder.R')
    
    filtered_obj <- DB_finder_removal(filtered_obj,
                        expected_doublet_rate = 0.08,
                        out_dir = db_removal_dir,
                        output_name = 'Doublet_finder',
                        output_col_name = 'doublet_status')
    
    # exclude doublets
    filtered_obj <- subset(filtered_obj, subset = doublet_status == 'Singlet')
    
    # check QC again after filtering
    pdf(file = paste0(working_dir, '/', 'QC_post_filtering.pdf'), width = 15, height = 5)
        print(
            VlnPlot(filtered_obj, features = c('nFeature_RNA', 'nCount_RNA', 'percent.mt'), group.by = 'sample_id',
                pt.size = 0)
        )
    dev.off()


    # simple dimension reduction of the datasets
    DR_obj <- filtered_obj %>% 
        NormalizeData() %>% 
        FindVariableFeatures(selection.method = "vst", nfeatures = 2000) %>% 
        ScaleData(verbose = FALSE) %>% 
        RunPCA(verbose = FALSE) %>% 
        RunUMAP(reduction = "pca", dims = 1:30) %>% 
        FindNeighbors(reduction = "pca", dims = 1:30)
    
    # check dimension reduction results
    pdf(file = paste0(working_dir, '/', 'Dimension_reduction.pdf'), width = 8, height = 7)
        print(
            FeaturePlot(DR_obj, features = 'nCount_RNA')
        )
        print(
            FeaturePlot(DR_obj, features = 'nFeature_RNA')
        )
        print(
            FeaturePlot(DR_obj, features = 'percent.mt')
        )
    dev.off()


    # ----
    # plot the expression of some markers
    marker_dir <- paste0(working_dir, '/', 'markers')
    dir.create(marker_dir, recursive = T)

    markers <- c(
        'MKI67', 'TOP2A', 'KRT5', 'KRT8', 'KRT18', 'KRT6B',
        'PDGFRA', 'PDGFRB', 'PDPN', 'CXCL12', 'IGF1R', 'ACTA2', 'COL1A1',
        'RGS5', 'MYH11', 'MYL9', 'MCAM',
        'PECAM1', 'VWF', 'FLT1', 'FLT4', 'CLEC4G', 'CLEC4M',
        'PTPRC', 'CD3D', 'CD4', 'CD8A', 'TRAC', 'TRBC1', 'CCR7', 'CD2',
        'ITGAX', 'CD14', 'CD68', 'AIF1',
        'NCAN1', 'NKG7', 'KLRD1',
        'HDC',
        'MS4A1', 'CD79A', 'CD19', 'IGHM', 'IGHG1',
        'MLANA', 'PMEL'
    )

    DefaultAssay(DR_obj) <- 'RNA'
    to_plot <- markers[markers %in% rownames(DR_obj)]
    for (g in to_plot) {
        pdf(paste0(marker_dir, '/', g, '.pdf'))
            print(FeaturePlot(DR_obj, features = g, reduction = 'umap', order = T))
        dev.off()
    }
 


    
    # ----
    # optimise clustring resolution
    clust_dir <- paste0(working_dir, '/', 'cluster_optimisation')
    dir.create(clust_dir, recursive = T)

    # load function
    source('/share/ScratchGeneral/tonwan/projects/snPATHO-Seq/scripts/function_scripts/02_Seurat_clustering_optimisation.R')

    # cluster and find optimal clustering resolution
    clustered_obj <- clustering_resolution_optimisation(DR_obj,
                                        resolution_range = seq(0.2, 2, by = 0.1),
                                        plot_dir = clust_dir,
                                        batch_factor = "processing_method",
                                        run_silhouette = FALSE,
                                        dist.matrix = NULL)

    # save processed data for further evaluation
    saveRDS(file = paste0(out_data_dir, '/', ds, '_clustered.rds'), clustered_obj)
}
```
