## Supplementary material for "snPATHO-seq: unlocking the pathology archives": scripts: 02_integrate_each_sample.R

```
# aim ----
    # take pre-processed data
    # integrate all data
    # derive joint cell type annotations


# define environment ----
.libPaths(
    c(
        '/share/ScratchGeneral/tonwan/projects/snPATHO-Seq/renv/library/R-4.1/x86_64-conda-linux-gnu',
        .libPaths()
    )
)
library(Seurat)
library(tidyverse)
library(RColorBrewer)


# arguments ----
data_dir <- '/share/ScratchGeneral/tonwan/projects/snPATHO-Seq/data/external_data/10X_data_processed'

out_dir <- '/share/ScratchGeneral/tonwan/projects/snPATHO-Seq/results/snRNA_external_data/10X_data_Tony/integration'

out_data_dir <- '/share/ScratchGeneral/tonwan/projects/snPATHO-Seq/data/external_data/10X_data_integrated'


# load predefined colors
source('/share/ScratchGeneral/tonwan/projects/snPATHO-Seq/scripts/colors.R')


# get names of the samples to be integrated
datasets <- list.files(data_dir)
sample_ids <- paste0(unlist(lapply(str_split(datasets, '_'), '[[', 1)), '_', unlist(lapply(str_split(datasets, '_'), '[[', 2)))


# load script for data integration
source('/share/ScratchGeneral/tonwan/projects/snPATHO-Seq/scripts/function_scripts/01_Seurat_integration.R')

# integrate each sample
for (s in unique(sample_ids)) {
    if (file.exists(paste0(out_data_dir, '/', s, '_integrated.rds'))) {
        next
    }

    working_datasets <- grep(s, datasets, value = T)
    working_dir <- paste0(out_dir, '/', s)
    dir.create(working_dir, recursive = T)

    obj_list <- list()
    obj_list[[1]] <- readRDS(file = paste0(data_dir, '/', working_datasets[[1]]))
    obj_list[[2]] <- readRDS(file = paste0(data_dir, '/', working_datasets[[2]]))

    integrated_obj <- seurat_integration(obj_list,
                                    normalisation_method = 'RNA',
                                    nfeatue_each_dataset = 2000,
                                    nfeatures_common = 2000,
                                    vars_to_regress = NULL,
                                    integration_method = 'cca',
                                    nthread = NULL)
    
    # only keep the selected workflow colors
    working_colors <- workflow_colors[names(workflow_colors) %in% unique(integrated_obj$processing_method)]

    # check integration results
    pdf(paste0(working_dir, '/', 'umap_batch_factors.pdf'))
        print(
            DimPlot(integrated_obj, reduction = "umap", group.by = 'processing_method')
        )
        print(
            FeaturePlot(integrated_obj, features = "nCount_RNA", reduction = 'umap') +
                scale_color_continuous(trans='log2', type = 'viridis')
        )
        print(
            FeaturePlot(integrated_obj, features = "nFeature_RNA", reduction = 'umap') +
                scale_color_continuous(trans='log2', type = 'viridis')
        )
        print(
            FeaturePlot(integrated_obj, features = "percent.mt", reduction = 'umap')
        )
    dev.off()

    # split UMAP by processing methods
    pdf(file = paste0(working_dir, '/', 'umap_split_by_sample_paper_figure.pdf'),
        width = 10,
        height = 5)
        print(
            DimPlot(integrated_obj, reduction = "umap", group.by = 'processing_method', 
                split.by = 'processing_method', ncol = 2, cols = working_colors) +
                labs(title = NULL)
        )
    dev.off()


    # cluster at different resolution to pick an optimal resolution
    clust_dir <- paste0(working_dir, '/', 'cluster_optimisation')
    dir.create(clust_dir, recursive = T)

    # load function
    source('/share/ScratchGeneral/tonwan/projects/snPATHO-Seq/scripts/function_scripts/02_Seurat_clustering_optimisation.R')

    # make sure the default assay is the integrated assay
    DefaultAssay(integrated_obj) <- 'integrated'
    # cleanup metadta
    integrated_obj@meta.data <- integrated_obj@meta.data[, c('orig.ident', 'nCount_RNA', 'nFeature_RNA', 'percent.mt',
                                                            'dataset_id', 'processing_method', 'sample_id')]
    # cluster and find optimal clustering resolution
    integrated_obj <- clustering_resolution_optimisation(integrated_obj,
                                        resolution_range = seq(0.2, 2, by = 0.1),
                                        plot_dir = clust_dir,
                                        batch_factor = "processing_method",
                                        run_silhouette = FALSE,
                                        dist.matrix = NULL)

    # save processed data
    saveRDS(file = paste0(out_data_dir, '/', s, '_integrated.rds'), integrated_obj)
}
```
