## Supplementary material for "snPATHO-seq: unlocking the pathology archives": scripts: GBM_annotation.R

```
# aim ----
    # annotate major cell types


# define environment ----
.libPaths(
    c(
        '/share/ScratchGeneral/tonwan/projects/snPATHO-Seq/renv/library/R-4.1/x86_64-conda-linux-gnu',
        .libPaths()
    )
)
library(Seurat)
library(tidyverse)
library(RColorBrewer)

library(ComplexHeatmap)
library(circlize)


# arguments ----
data_file <- '/share/ScratchGeneral/tonwan/projects/snPATHO-Seq/data/external_data/10X_data_integrated/Glioblastoma_1773A_integrated.rds'

out_dir <- '/share/ScratchGeneral/tonwan/projects/snPATHO-Seq/results/snRNA_external_data/10X_data_Tony/integration/Glioblastoma_1773A'
dir.create(out_dir, recursive = T)

sample_id <- 'Glioblastoma_1773A'

out_data_dir <- '/share/ScratchGeneral/tonwan/projects/snPATHO-Seq/data/external_data/10X_data_annotated'
dir.create(out_dir, recursive = T)


# load data ----
obj <- readRDS(file = data_file)


# check the expression of selected cell type markers 
marker_dir <- paste0(out_dir, '/', 'markers')
dir.create(marker_dir, recursive = T)

markers <- c(
    'KRT8', 'KRT18', 'EGFR', 'MKI67', 'ESR1', 'TOP2A', 'EPCAM',
    'PDGFRA', 'PDGFRB', 'COL1A1', 'FAP', 'FAK',
    'ACTA2', 'MYH11', 'MYL9', 'MCAM', 'RGS5', 
    'PECAM1', 'VWF', 'FLT1', 'PROX1',
    'PTPRC', 'CD3D', 'CD2', 'TRAC', 'TRBC1', 'CD4', 'CD8A', 'FOXP3',
    'MS4A1', 'CD79A', 'CD19',
    'ITGAX', 'CD68', 'CD163', 'TNF', 'CD80', 'CD86', 'NOS2', 'S100A8', 'S100A9',
    'CD14', 'CCR2', 'FCGR3A',
    'CLEC10A', 'CLEC9A', 'XCR1',
    'ITGAM',
    'NCAM1', 'NKG7',
    'SPTA1', 'HBB', 'HBA',
    'NRXN1', 'CADM2', 'NRXN3',
    'DNAH12', 'DNAAF1', 'DNAH7', 'CD27', 'IRF4', 'BCL6', 'IGHG1', 'JCHAIN',
    'ITM2C', 'PLD4', 'SERPINF1', 'LILRA4', 'IL3RA', 'TPM2', 'MZB1', 'SPIB', 'IRF4', 'SMPD3',
    'TUBB3', 'MAP2',
    'SPARCL1', 'GFAP', 'CLU', 'CRYAB', 'TTYH1', 'SLC1A3', 'CST3', 'ID3', 'AGT',
    'STMN2', 'DCX', 'STMN4', 'TAGLN3', 'ELAVL4', 'MLLT11', 'NREP', 'MIAT',
    'BTG2', 'SOX6', 'BAZ2B', 'SOX4', 'DSEL', 'MXD4', 'PNRC1', 'SOX5',
    'EMP1', 'GAP43', 'TNFRSF12A', 'CD44', 'TNC', 'LYST', 'PLAT', 'WWTR1',
    'APOD', 'MBP', 'PLA2G16', 'PLP1', 'PPP1R14A', 'SEPP1', 'TF', 'GSN', 'PTGDS', 'MAG', 'MOG', 'CLDN11')


DefaultAssay(obj) <- 'RNA'
to_plot <- markers[markers %in% rownames(obj)]
for (g in to_plot) {
    pdf(paste0(marker_dir, '/', g, '.pdf'))
        print(FeaturePlot(obj, features = g, reduction = 'umap', order = T))
    dev.off()
}


# score cells using published gene modules to better determine cell types
GM_dir <- paste0(out_dir, '/', 'gene_module_scores')
dir.create(GM_dir, recursive = T)

NPC_GBM <- c('STMN2','DCX','STMN4','TAGLN3','ELAVL4','MLLT11','NREP','MIAT','TUBB2A','CD24','HMP19','RND3','SOX11','SOX4','TUBB3','MAP1B','NSG1','SNAP25','FNBP1L','HN1','IGFBPL1','PKIA','NNAT','RBP1','SCG3','DLX6-AS1','KLHDC8A','MEG3','RBFOX2','SCG2','STMN1','TMEM161B-AS1','ARL4D','BEX2','DDAH2','DLX5','GAP43','CRMP1','ENO2','TCEAL7','UCHL1','BASP1','CXADR','GNG3','GPC2','KIF5C','PAK3','RTN1','TERF2IP','SYT1')
oligo_prog <- c('SIRT2','GPR17','OLIG1','APOD','DLL3','PLP1','NEU4','BCAN','SOX8','TNR','OMG','TMEM206','LPPR1','NKAIN4','FIBIN','SGK1','SLC1A1','CA10','LMF1','PTGDS','SERINC5','ASCL1','DLL1','OLIG2','PSAT1','RAB33A','SCRG1','CNP','LIMA1','THY1','TUBB4A','BCAS1','CNTN1','HES6','LHFPL3','PDGFRA','RGR','SCD5','ANGPTL2','DHCR7','FGF12','MPZL1','MSMO1','P2RX7','RTKN','TNFRSF21','SMOC1','ACAT2','FDFT1','PGRMC1')
oligo_normal <- c('APOD','MBP','PLA2G16','PLP1','PPP1R14A','SEPP1','TF','GSN','PTGDS','APLP1','CLDN11','CLDND1','CNP','RNASE1','SCD','44443','ANLN','CRYAB','ELOVL1','HSPA2','PLLP','SIRT2','SLC44A1','BIN1','CMTM5','ERMN','FAM107B','LARP6','LHPP','MAG','MOBP','PLEKHB1','QDPR','S100A1','SLAIN1','SLC48A1','TUBB4A','TMEM144','ABCA2','CA2','KLK6','MAL','SPOCK3','HAPLN2','UGT8','STMN4','ATP1B1','44447','PACS2','ANKS1B')
NPC_OPC <- c('BTG2','SOX6','BAZ2B','SOX4','DSEL','MXD4','PNRC1','SOX5','CHD7','GRIA2','MAP3K1','OLIG1','SESN3','UBE2H','ABAT','DCX','DOCK10','GLCCI1','KCNQ1OT1','KDM5B','KLHL24','MAP2','OLIG2','PEAK1','RAB3IP','RFTN2','RFX4','TANC2','TRIO','VCAN','ZNF91','FAM181B','STMN4','DLL3','BTG1','TXNIP','LDLRAD3','SOX11','HIP1','FGFBP3','MTSS1','SPRY4','REV3L','MMP16','KLF12','TFDP2','CADM2','ARHGEF7','GOLGB1','ZMYM2')
MES_GBM <- c('EMP1','GAP43','TNFRSF12A','CD44','TNC','LYST','PLAT','WWTR1','ANXA2','GADD45A','LMNA','VMP1','IGFBP3','SCG2','HIVEP3','METTL7B','RCAN1','ZYX','CAMK2D','IL1RAP','JAG1','NAMPT','PGM2L1','VIM','ACTN1','AKAP12','IGF2BP2','MIR4435-2HG','NRP2','PDLIM4','ANXA1','BCAT1','CLIC4','CREB5','DIRAS3','EGR1','ELL2','F3','FRMD5','IGFBP7','LINC00152','S100A16','VCL','SAMD4A','ADAMTS9-AS2','SLC4A7','GFAP','CADPS','GNG12','KLHL4')
Astrocyte <- c('SPARCL1','GFAP','CLU','CRYAB','TTYH1','SLC1A3','CST3','ID3','AGT','APOE','EDNRB','ALDOC','AQP4','GATM','NDRG2','SPARC','MT2A','MT3','BCAN','HEPN1','NTRK2','PMP2','PON2','ATP1A2','HEY1','MLC1','PLTP','ATP1B2','ITM2C','MT1X','VIM','S100B','AHCYL1','RAMP1','SCRG1','C1ORF61','IGFBP7','RASSF4','ID1','IFITM3','CPE','CD9','FABP7','GRAMD3','ZFP36','F3','PLP1','ID4','PSAT1','GJA1')

gm_list <- list(NPC_GBM, oligo_prog, oligo_normal, NPC_OPC, MES_GBM, Astrocyte)
names(gm_list) <- c('NPC_GBM', 'oligo_prog', 'oligo_normal', 'NPC_OPC', 'MES_GBM', 'Astrocyte')

for (n in names(gm_list)) {
    annotated_obj <- AddModuleScore(
        object = obj,
        features = gm_list[n],
        assay = 'RNA',
        name = n)
    annotated_obj@meta.data[[n]] <- annotated_obj@meta.data[[paste0(n, '1')]] 
    pdf(file = paste0(GM_dir, '/', n, '.pdf'))
        print(
            FeaturePlot(annotated_obj, features = n, order = TRUE) +
                scale_color_gradient2(low = '#1878b2', mid = '#e5e5e5', high = '#B2182B')
        )
    dev.off()
}


########################################################################################################################
########################################################################################################################
########################################################################################################################
optimal_clustering_dir <- paste0(out_dir, '/', 'optimal_cluster')
dir.create(optimal_clustering_dir, recursive = T)


opt_res <- 0.8
cluster_name <- grep(paste0('res.', opt_res), colnames(obj@meta.data), value = T)[1]


# make a plot with label at the optimal cluster resolution ----
pdf(file = paste0(optimal_clustering_dir, '/', 'umap_', opt_res, '.pdf'))
    print(
        DimPlot(obj, reduction = "umap", group.by = cluster_name, label = T, repel = T)
    )
dev.off()


# check markers for selected clusters
Idents(obj) <- cluster_name

for (cl in c(8)) {
    de_markrers <- FindMarkers(obj, ident.1 = cl, test.use = 't')
    write.csv(file = paste0(optimal_clustering_dir, '/', 'Cluster_', cl, '_markers.csv'), de_markrers)
}


####################################################################################
####################################################################################
####################################################################################
# annotate clusters ----
Idents(obj) <- cluster_name

new_annotation_df <- data.frame(cluster_id = unique(as.character(Idents(obj))),
                                initial_annotation = '')
new_annotation_df[new_annotation_df$cluster_id == '0', 'initial_annotation'] <- 'OPC_like'
new_annotation_df[new_annotation_df$cluster_id == '1', 'initial_annotation'] <- 'Macrophages'
new_annotation_df[new_annotation_df$cluster_id == '2', 'initial_annotation'] <- 'NPC_like'
new_annotation_df[new_annotation_df$cluster_id == '3', 'initial_annotation'] <- 'OPC_like'
new_annotation_df[new_annotation_df$cluster_id == '4', 'initial_annotation'] <- 'NPC_like'
new_annotation_df[new_annotation_df$cluster_id == '5', 'initial_annotation'] <- 'Mixed'
new_annotation_df[new_annotation_df$cluster_id == '6', 'initial_annotation'] <- 'OPC_like'
new_annotation_df[new_annotation_df$cluster_id == '7', 'initial_annotation'] <- 'Macrophages'
new_annotation_df[new_annotation_df$cluster_id == '8', 'initial_annotation'] <- 'Unassigned'
new_annotation_df[new_annotation_df$cluster_id == '9', 'initial_annotation'] <- 'NPC_like'
new_annotation_df[new_annotation_df$cluster_id == '10', 'initial_annotation'] <- 'MES_like'
new_annotation_df[new_annotation_df$cluster_id == '11', 'initial_annotation'] <- 'Oligodendrocyte_progenitor_cells'
new_annotation_df[new_annotation_df$cluster_id == '12', 'initial_annotation'] <- 'AC_like'
new_annotation_df[new_annotation_df$cluster_id == '13', 'initial_annotation'] <- 'Macrophages'
new_annotation_df[new_annotation_df$cluster_id == '14', 'initial_annotation'] <- 'T'
new_annotation_df[new_annotation_df$cluster_id == '15', 'initial_annotation'] <- 'CAF'
new_annotation_df[new_annotation_df$cluster_id == '16', 'initial_annotation'] <- 'NPC_like'
new_annotation_df[new_annotation_df$cluster_id == '17', 'initial_annotation'] <- 'Macrophages'
new_annotation_df[new_annotation_df$cluster_id == '18', 'initial_annotation'] <- 'NPC_like'
new_annotation_df[new_annotation_df$cluster_id == '19', 'initial_annotation'] <- 'Endothelial'
new_annotation_df[new_annotation_df$cluster_id == '20', 'initial_annotation'] <- 'Pericytes'
new_annotation_df[new_annotation_df$cluster_id == '21', 'initial_annotation'] <- 'MES_like'
new_annotation_df[new_annotation_df$cluster_id == '22', 'initial_annotation'] <- 'AC_like'


# annotate the object
new_annotations <- new_annotation_df$initial_annotation
names(new_annotations) <- new_annotation_df$cluster_id

obj <- RenameIdents(obj, new_annotations)
obj$initial_annotation <- Idents(obj)


# export cell type annotations as csv file for easier modification and assessemnt
write.csv(file = paste0(optimal_clustering_dir, '/', 'initial_annotation.csv'), obj$initial_annotation)


# plot a umap with new annotations
Idents(obj) <- 'initial_annotation'
pdf(file = paste0(optimal_clustering_dir, '/', 'umap_initial_annotation.pdf'), width = 10, height = 7)
    print(
        DimPlot(obj, reduction = "umap", group.by = 'initial_annotation', label = T, repel = T)
    )
dev.off()


####################################################################################
####################################################################################
####################################################################################
# save processed object 
saveRDS(file = paste0(out_data_dir, '/', sample_id, '_annotated.rds'), obj)


####################################################################################
####################################################################################
####################################################################################
# plot for the paper
paper_fig_dir <- paste0(out_dir, '/', 'paper_figures')
dir.create(paper_fig_dir, recursive = T)

# define colors
# generate colors for cell types
cell_colors <- scales::hue_pal()(length(unique(obj$initial_annotation)))
names(cell_colors) <- unique(obj$initial_annotation)

# load predefined colors for workflows
source('/share/ScratchGeneral/tonwan/projects/snPATHO-Seq/scripts/colors.R')
workflow_colors <- workflow_colors[names(workflow_colors) %in% unique(obj$processing_method)]


# plot umap with annotation
plot <- DimPlot(obj, group.by = 'initial_annotation', cols = cell_colors) +
            labs(title = NULL)
plot <- LabelClusters(plot = plot, id = "initial_annotation", repel = TRUE, box = TRUE, force = 1, color = 'white')

pdf(file = paste0(paper_fig_dir, '/', 'umap_with_annotation.pdf'), width = 8)
    print(
        plot
    )     
dev.off()


# split umap by processing methods
pdf(file = paste0(paper_fig_dir, '/', 'umap_split_by_workflows.pdf'),
    width = 8,
    height = 5)
    print(
        DimPlot(obj, reduction = "umap", group.by = 'processing_method', 
            split.by = 'processing_method', ncol = 3, cols = workflow_colors) +
            labs(title = NULL)
    )
dev.off()


########################################################################################################################
########################################################################################################################
########################################################################################################################
# run DE between annotated cell types
Idents(obj) <- 'initial_annotation'
de_res <- FindAllMarkers(obj, assay = 'RNA', test.use = 't', only.pos = TRUE)
write.csv(file = paste0(paper_fig_dir, '/', 'de_between_major_cell_types.csv'), de_res)


probe_ref_file <- '/share/ScratchGeneral/tonwan/projects/snPATHO-Seq/resources/Chromium_Human_Transcriptome_Probe_Set_v1.0_GRCh38-2020-A.csv'
probe_ref <- read.csv(file = probe_ref_file, skip = 5)
probe_ref <- probe_ref[probe_ref$included == TRUE, ]
gene_names <- unique(unlist(lapply(str_split(probe_ref$probe_id, '\\|'), '[[', 2)))

de_res <- de_res[de_res$gene %in% gene_names, ]


# plot expression of top DE genes between clusters - annotate canonical cell type markers
# get top 200 DE genes from each cluster to plot
top_markers <- de_res %>% 
    group_by(cluster) %>%
    filter(p_val_adj < 0.05) %>%
    slice_max(avg_log2FC, n = 200) %>% 
    pull(gene) %>% 
    unique()


# get mean expresion of the selected genes for each processing method 
plot_mtx <- as.data.frame(AverageExpression(obj, assays = 'RNA', slot = 'data', features = top_markers, group.by = c('processing_method', 'initial_annotation'))[[1]])
plot_mtx$gene_names <- rownames(plot_mtx)
plot_mtx_long <- gather(plot_mtx, key = 'annotation', value = 'mean_expression', -gene_names)

# annotate the expression dataframe
plot_mtx_long$processing_method <- unlist(lapply(str_split(plot_mtx_long$annotation, '_'), '[[', 1))
plot_mtx_long$cell_type <- unlist(lapply(str_split(plot_mtx_long$annotation, '\\_'), function(x) paste(x[-1], collapse = '_')))


# scale the gene expression within each processing method
plot_mtx_long <- plot_mtx_long %>% 
        group_by(gene_names, processing_method) %>%
        mutate(scale(mean_expression)) %>% 
        as.data.frame() 
  
# plot_order <- plot_mtx_long %>% 
#     arrange(as.factor(cell_type)) %>% 
#     pull(annotation) %>% 
#     unique()

plot_order <- plot_mtx_long %>% 
    arrange(factor(cell_type, 
        levels = c(
            'OPC_like', 'Oligodendrocyte_progenitor_cells', 'Macrophages', 'T', 'Unassigned', 'Pericytes',
            'Endothelial', 'CAF', 'AC_like', 'MES_like', 'NPC_like', 'Mixed'))) %>% 
    pull(annotation) %>% 
    unique()

scaled_mtx <- spread(plot_mtx_long[, c('gene_names', 'annotation', 'scale(mean_expression)')],
        value = 'scale(mean_expression)', key = 'annotation')
scaled_mtx <- column_to_rownames(scaled_mtx, var = 'gene_names')
# cap the scaled expression at -5 to 5
scaled_mtx[scaled_mtx > 5] <- 5
scaled_mtx[scaled_mtx < -5] <- -5
scaled_mtx[is.na(scaled_mtx)] <- 0
scaled_mtx <- scaled_mtx[, plot_order]

# make heatmap annotation
annot_df <- plot_mtx_long[, c('annotation', 'processing_method', 'cell_type')]
annot_df <- distinct(annot_df)
annot_df <- annot_df[match(colnames(scaled_mtx), annot_df$annotation), ]


# set colors
col_fun <- colorRamp2(c(-5, 0, 5), 
                      c("#2166AC", "white", "#B2182B"))


# make annotation
left_annot <- HeatmapAnnotation(
        Workflows = annot_df$processing_method,
        which = 'row',
        col = list(Workflows = workflow_colors), 
        annotation_name_gp = grid::gpar(fontsize = 12),
        annotation_legend_param = list(
            Workflows = list(title_gp = gpar(fontsize = 12, fontface = "bold"), 
                                    labels_gp = gpar(fontsize = 12))))


# left_annot <- HeatmapAnnotation(
#         Workflows = annot_df$processing_method,
#         Cell_types = annot_df$cell_type,
#         which = 'row',
#         col = list(Workflows = workflow_colors,
#                 Cell_types = cell_colors), 
#         annotation_name_gp = grid::gpar(fontsize = 12),
#         annotation_legend_param = list(
#             Workflows = list(title_gp = gpar(fontsize = 12, fontface = "bold"), 
#                                     labels_gp = gpar(fontsize = 12)),
#             Cell_types = list(title_gp = gpar(fontsize = 12, fontface = "bold"), 
#                                     labels_gp = gpar(fontsize = 12))))


# annotate selected genes 
genes_to_plot <- de_res %>% 
    group_by(cluster) %>%
    filter(p_val_adj < 0.05) %>%
    slice_max(avg_log2FC, n = 5) %>% 
    pull(gene)

# make sure the selected markers are in the matrix
all_markers <- genes_to_plot[genes_to_plot %in% rownames(scaled_mtx)]

# get location of the genes to be annotated
mtx_index <- seq(1:nrow(scaled_mtx))
names(mtx_index) <- rownames(scaled_mtx)
position_in_matrix <- mtx_index[names(mtx_index) %in% all_markers]

bottom_annotation <- HeatmapAnnotation(
    markers = anno_mark(
        at = position_in_matrix, 
        labels = names(position_in_matrix),
        which = "column", side = "bottom",
        labels_gp = gpar(fontsize = 8)))


# plot heatmap ----
plot <- Heatmap(t(scaled_mtx),
                name = 'Scaled mean expression',
                heatmap_legend_param = list(title_gp = gpar(fontsize = 12, fontface = "bold")),
                col = col_fun,
                left_annotation = left_annot,
                bottom_annotation = bottom_annotation,
                show_column_names= FALSE,
                show_row_names = TRUE,
                cluster_rows = FALSE,
                cluster_columns = TRUE,
                split = annot_df$processing_method,
                column_names_gp = gpar(fontsize = 6),
                row_names_gp = gpar(fontsize = 8))


pdf(file = paste0(paper_fig_dir, '/', 'top_DE_gene_heatmap', '.pdf'),
    width = 14, height = 7)
    draw(plot)
dev.off()
```
