## Supplementary material for "snPATHO-seq: unlocking the pathology archives": scripts: Kidney_annotation.R

```
# aim ----
    # annotate major cell types


# define environment ----
.libPaths(
    c(
        '/share/ScratchGeneral/tonwan/projects/snPATHO-Seq/renv/library/R-4.1/x86_64-conda-linux-gnu',
        .libPaths()
    )
)
library(Seurat)
library(tidyverse)
library(RColorBrewer)

library(ComplexHeatmap)
library(circlize)


# arguments ----
data_file <- '/share/ScratchGeneral/tonwan/projects/snPATHO-Seq/data/external_data/10X_data_integrated/Kidney_1305272B_integrated.rds'

out_dir <- '/share/ScratchGeneral/tonwan/projects/snPATHO-Seq/results/snRNA_external_data/10X_data_Tony/integration/Kidney_1305272B'
dir.create(out_dir, recursive = T)

sample_id <- 'Kidney_1305272B'

out_data_dir <- '/share/ScratchGeneral/tonwan/projects/snPATHO-Seq/data/external_data/10X_data_annotated'
dir.create(out_dir, recursive = T)


# load data ----
obj <- readRDS(file = data_file)


# check the expression of selected cell type markers 
marker_dir <- paste0(out_dir, '/', 'markers')
dir.create(marker_dir, recursive = T)

markers <- c(
    'KRT8', 'KRT18', 'EGFR', 'MKI67', 'ESR1', 'TOP2A', 'EPCAM',
    'PDGFRA', 'PDGFRB', 'COL1A1', 'FAP', 'FAK',
    'ACTA2', 'MYH11', 'MYL9', 'MCAM', 'RGS5', 
    'PECAM1', 'VWF', 'FLT1', 'PROX1',
    'PTPRC', 'CD3D', 'CD2', 'TRAC', 'TRBC1', 'CD4', 'CD8A', 'FOXP3',
    'MS4A1', 'CD79A', 'CD19',
    'ITGAX', 'CD68', 'CD163', 'TNF', 'CD80', 'CD86', 'NOS2', 'S100A8', 'S100A9',
    'CD14', 'CCR2', 'FCGR3A',
    'CLEC10A', 'CLEC9A', 'XCR1',
    'ITGAM',
    'NCAM1', 'NKG7',
    'SPTA1', 'HBB', 'HBA',
    'NRXN1', 'CADM2', 'NRXN3', 'TUBB3', 'MAP2',
    'NPHS2','ADAMTS19','WT1','FMN2','NTNG1','PTPRQ','NPHS1','PTPRO','ZNF804A','ST6GALNAC3',
    'SH3GL3','PLCB1','LGR4','BICDL1','PAX8','MAL','CACNA2D3','CDR2','PTPN14','PKHD1',
    'LINC01099','PDE4D','NR3C2','SCN2A','CADPS2','COBLL1','TOX3','DCDC2','SLC14A2','ABTB2',
    'MYO9A','LINC01320','FOXP2','PAX8','BICC1','OXR1','PKHD1',
    'SLC12A3','CACNA1D','LINC01055','ARHGAP6','FGF13','TRPM6','TRPM7','ESRRG','ATP1A1','LINC01762',
    'LINC01187','SLC4A9','ATP6V1C2','GALNT17','KIT','IL18','CLNK','ATP6V0D2','RHCG','ITGA6',
    'KRT13','EEF1A1','S100A6','EHF','AMBRA1','LCN2','SLPI','TACSTD2','RPL41','RPLP1',
    'KCNT2','ALDH1A2','CFH','PAWR','FRMD4A','SLC4A11','LINC01435','PDE1A','RHEX','SYNE1',
    'KSR2','ABTB2','GRIP1','EYA4','GATA3','ANK3','PRKAG2','C11orf80','MECOM','PAPPA',
    'LINC01020','PLG','AGXT2','ACMSD','SLC36A2','SLC13A1','CDH9','LINC02027','CUBN','RAB11FIP3',
    'CDH19','NRXN1','NCAM1','PRIMA1','CADM2','ZNF536','ADGRB3','EPB41L2','STARD13','SCN7A',
    'GP2','SLC12A1','CASR','CLCN5','PLCB1','CGNL1','ESRRB','UMOD','PKP4','CACNA2D3',
    'RAMP3','TLL1','PLVAP','CEACAM1','TEK','PLAT','ADGRL4','LIMS2','LINC02147','GPM6A',
    'PALMD','ADGRL4','ADAMTS6','EFNB2','CDH13','SYN3','PTPRB','PLPP1','RUNX1T1','TIMP3',
    'ABCC8','REN','COL13A1','CACNB2','ADCY3','PIP5K1B','GRID2','WFDC1','PRKG1','SLIT3',
    'PWRN1','GATA3','AQP2','PDE4D','KSR2','PIK3C2G','GRIP1','ABTB2','MECOM','LINC01482',
    'GALNT17','RHCG','ATP6V1C2','ATP6V0D2','CLNK','KIT','PDE1C','PACRG','PTGER3','SLC26A7',
    'LINC01924','ANK3','CAPN8','ABTB2','SAMD12','SLC14A2','GRIP1','NAALADL2','LRBA','PCDH7',
    'SLC4A9','SLC26A4','LINC01187','SLC35F3','IL18','DGKI','PPM1E','PDZD2','PDE1C','ATP6V0D2',
    'TBX1','CCL21','MMRN1','PKHD1L1','SEMA3A','FLRT2','RELN','GRAPL','ELK3','KALRN',
    'NOS1','TMPRSS4','PPFIA2','BBOX1','PAPPA2','ERBB4','ITGB6','CALCR','PKP4','DEPTOR',
    'GRIP1','ABTB2','BMPR1B','ANK3','PRKAG2','MECOM','TRPM3','PIK3C2G','PAPPA','CADPS2',
    'AFM','SLC36A2','PLG','PAH','SLC13A1','AGXT2','SLC34A1','ACMSD','SLC17A1','SLC28A1',
    'SMIM2-AS1','SLC13A3','ACSM2B','ACSM2A','CLDN10','SLC2A9','CRYL1','SUGCT','SLC5A12','ERRFI1',
    'AGXT2','CDH9','LINC00671','ACMSD','SLC28A1','SMIM2-AS1','SLC13A1','TINAG','SLC6A13','DPYS')


DefaultAssay(obj) <- 'RNA'
to_plot <- markers[markers %in% rownames(obj)]
for (g in to_plot) {
    if (file.exists(paste0(marker_dir, '/', g, '.pdf'))) {
        next
    }
    pdf(paste0(marker_dir, '/', g, '.pdf'))
        print(FeaturePlot(obj, features = g, reduction = 'umap', order = T))
    dev.off()
}


########################################################################################################################
########################################################################################################################
########################################################################################################################
optimal_clustering_dir <- paste0(out_dir, '/', 'optimal_cluster')
dir.create(optimal_clustering_dir, recursive = T)


opt_res <- 1.1
cluster_name <- grep(paste0('res.', opt_res), colnames(obj@meta.data), value = T)[1]


# make a plot with label at the optimal cluster resolution ----
pdf(file = paste0(optimal_clustering_dir, '/', 'umap_', opt_res, '.pdf'))
    print(
        DimPlot(obj, reduction = "umap", group.by = cluster_name, label = T, repel = T)
    )
dev.off()


# check markers for selected clusters
Idents(obj) <- cluster_name

for (cl in c(5, 9)) {
    de_markrers <- FindMarkers(obj, ident.1 = cl, test.use = 't')
    write.csv(file = paste0(optimal_clustering_dir, '/', 'Cluster_', cl, '_markers.csv'), de_markrers)
}


####################################################################################
####################################################################################
####################################################################################
# annotate clusters ----
Idents(obj) <- cluster_name

new_annotation_df <- data.frame(cluster_id = unique(as.character(Idents(obj))),
                                initial_annotation = '')
new_annotation_df[new_annotation_df$cluster_id == '0', 'initial_annotation'] <- 'Distal_convoluted_tubule'
new_annotation_df[new_annotation_df$cluster_id == '1', 'initial_annotation'] <- 'Loop_of_Henle_epithelial'
new_annotation_df[new_annotation_df$cluster_id == '2', 'initial_annotation'] <- 'Collecting_tubule_and_connecting_tubule'
new_annotation_df[new_annotation_df$cluster_id == '3', 'initial_annotation'] <- 'Proximal_tubule'
new_annotation_df[new_annotation_df$cluster_id == '4', 'initial_annotation'] <- 'Proximal_tubule'
new_annotation_df[new_annotation_df$cluster_id == '5', 'initial_annotation'] <- 'Collecting_tubule_and_connecting_tubule'
new_annotation_df[new_annotation_df$cluster_id == '6', 'initial_annotation'] <- 'Intercalated_type_A'
new_annotation_df[new_annotation_df$cluster_id == '7', 'initial_annotation'] <- 'Loop_of_Henle_epithelial'
new_annotation_df[new_annotation_df$cluster_id == '8', 'initial_annotation'] <- 'Endothelial'
new_annotation_df[new_annotation_df$cluster_id == '9', 'initial_annotation'] <- 'Proximal_tubule'
new_annotation_df[new_annotation_df$cluster_id == '10', 'initial_annotation'] <- 'Loop_of_Henle_epithelial'
new_annotation_df[new_annotation_df$cluster_id == '11', 'initial_annotation'] <- 'Proximal_tubule'
new_annotation_df[new_annotation_df$cluster_id == '12', 'initial_annotation'] <- 'Proximal_tubule'
new_annotation_df[new_annotation_df$cluster_id == '13', 'initial_annotation'] <- 'Collecting_tubule_and_connecting_tubule'
new_annotation_df[new_annotation_df$cluster_id == '14', 'initial_annotation'] <- 'Parietal_epithelial'
new_annotation_df[new_annotation_df$cluster_id == '15', 'initial_annotation'] <- 'Collecting_tubule_and_connecting_tubule'
new_annotation_df[new_annotation_df$cluster_id == '16', 'initial_annotation'] <- 'Podocytes'
new_annotation_df[new_annotation_df$cluster_id == '17', 'initial_annotation'] <- 'Fibroblasts'
new_annotation_df[new_annotation_df$cluster_id == '18', 'initial_annotation'] <- 'Intercalated_type_A'
new_annotation_df[new_annotation_df$cluster_id == '19', 'initial_annotation'] <- 'Pericytes'
new_annotation_df[new_annotation_df$cluster_id == '20', 'initial_annotation'] <- 'Mixed_immune_cells'
new_annotation_df[new_annotation_df$cluster_id == '21', 'initial_annotation'] <- 'Loop_of_Henle_epithelial'
new_annotation_df[new_annotation_df$cluster_id == '22', 'initial_annotation'] <- 'Endothelial'
new_annotation_df[new_annotation_df$cluster_id == '23', 'initial_annotation'] <- 'Endothelial'
new_annotation_df[new_annotation_df$cluster_id == '24', 'initial_annotation'] <- 'Intercalated_type_B'


# annotate the object
new_annotations <- new_annotation_df$initial_annotation
names(new_annotations) <- new_annotation_df$cluster_id

obj <- RenameIdents(obj, new_annotations)
obj$initial_annotation <- Idents(obj)


# export cell type annotations as csv file for easier modification and assessemnt
write.csv(file = paste0(optimal_clustering_dir, '/', 'initial_annotation.csv'), obj$initial_annotation)


# plot a umap with new annotations
Idents(obj) <- 'initial_annotation'
pdf(file = paste0(optimal_clustering_dir, '/', 'umap_initial_annotation.pdf'), width = 10, height = 7)
    print(
        DimPlot(obj, reduction = "umap", group.by = 'initial_annotation', label = T, repel = T)
    )
dev.off()


####################################################################################
####################################################################################
####################################################################################
# save processed object 
saveRDS(file = paste0(out_data_dir, '/', sample_id, '_annotated.rds'), obj)


####################################################################################
####################################################################################
####################################################################################
# plot for the paper
paper_fig_dir <- paste0(out_dir, '/', 'paper_figures')
dir.create(paper_fig_dir, recursive = T)

# define colors
# generate colors for cell types
cell_colors <- scales::hue_pal()(length(unique(obj$initial_annotation)))
names(cell_colors) <- unique(obj$initial_annotation)

# load predefined colors for workflows
source('/share/ScratchGeneral/tonwan/projects/snPATHO-Seq/scripts/colors.R')
workflow_colors <- workflow_colors[names(workflow_colors) %in% unique(obj$processing_method)]


# plot umap with annotation
plot <- DimPlot(obj, group.by = 'initial_annotation', cols = cell_colors) +
            labs(title = NULL)
# plot <- LabelClusters(plot = plot, id = "initial_annotation", repel = TRUE, box = TRUE, force = 1, color = 'white')

pdf(file = paste0(paper_fig_dir, '/', 'umap_with_annotation.pdf'), width = 8)
    print(
        plot
    )     
dev.off()


# split umap by processing methods
pdf(file = paste0(paper_fig_dir, '/', 'umap_split_by_workflows.pdf'),
    width = 8,
    height = 5)
    print(
        DimPlot(obj, reduction = "umap", group.by = 'processing_method', 
            split.by = 'processing_method', ncol = 3, cols = workflow_colors) +
            labs(title = NULL)
    )
dev.off()


########################################################################################################################
########################################################################################################################
########################################################################################################################
# run DE between annotated cell types
Idents(obj) <- 'initial_annotation'
de_res <- FindAllMarkers(obj, assay = 'RNA', test.use = 't', only.pos = TRUE)
write.csv(file = paste0(paper_fig_dir, '/', 'de_between_major_cell_types.csv'), de_res)


probe_ref_file <- '/share/ScratchGeneral/tonwan/projects/snPATHO-Seq/resources/Chromium_Human_Transcriptome_Probe_Set_v1.0_GRCh38-2020-A.csv'
probe_ref <- read.csv(file = probe_ref_file, skip = 5)
probe_ref <- probe_ref[probe_ref$included == TRUE, ]
gene_names <- unique(unlist(lapply(str_split(probe_ref$probe_id, '\\|'), '[[', 2)))

de_res <- de_res[de_res$gene %in% gene_names, ]


# plot expression of top DE genes between clusters - annotate canonical cell type markers
# get top 200 DE genes from each cluster to plot
top_markers <- de_res %>% 
    group_by(cluster) %>%
    filter(p_val_adj < 0.05) %>%
    slice_max(avg_log2FC, n = 200) %>% 
    pull(gene) %>% 
    unique()


# get mean expresion of the selected genes for each processing method 
plot_mtx <- as.data.frame(AverageExpression(obj, assays = 'RNA', slot = 'data', features = top_markers, group.by = c('processing_method', 'initial_annotation'))[[1]])
plot_mtx$gene_names <- rownames(plot_mtx)
plot_mtx_long <- gather(plot_mtx, key = 'annotation', value = 'mean_expression', -gene_names)

# annotate the expression dataframe
plot_mtx_long$processing_method <- unlist(lapply(str_split(plot_mtx_long$annotation, '_'), '[[', 1))
plot_mtx_long$cell_type <- unlist(lapply(str_split(plot_mtx_long$annotation, '\\_'), function(x) paste(x[-1], collapse = '_')))


# scale the gene expression within each processing method
plot_mtx_long <- plot_mtx_long %>% 
        group_by(gene_names, processing_method) %>%
        mutate(scale(mean_expression)) %>% 
        as.data.frame() 
  
# plot_order <- plot_mtx_long %>% 
#     arrange(as.factor(cell_type)) %>% 
#     pull(annotation) %>% 
#     unique()

plot_order <- plot_mtx_long %>% 
    arrange(factor(cell_type, 
        levels = c(
            'Intercalated_type_A', 'Intercalated_type_B', 'Collecting_tubule_and_connecting_tubule',
            'Distal_convoluted_tubule', 'Podocytes', 'Mixed_immune_cells', 'Parietal_epithelial',
            'Fibroblasts', 'Pericytes', 'Proximal_tubule', 'Endothelial', 'Loop_of_Henle_epithelial'))) %>% 
    pull(annotation) %>% 
    unique()


scaled_mtx <- spread(plot_mtx_long[, c('gene_names', 'annotation', 'scale(mean_expression)')],
        value = 'scale(mean_expression)', key = 'annotation')
scaled_mtx <- column_to_rownames(scaled_mtx, var = 'gene_names')
# cap the scaled expression at -5 to 5
scaled_mtx[scaled_mtx > 5] <- 5
scaled_mtx[scaled_mtx < -5] <- -5
scaled_mtx[is.na(scaled_mtx)] <- 0
scaled_mtx <- scaled_mtx[, plot_order]

# make heatmap annotation
annot_df <- plot_mtx_long[, c('annotation', 'processing_method', 'cell_type')]
annot_df <- distinct(annot_df)
annot_df <- annot_df[match(colnames(scaled_mtx), annot_df$annotation), ]


# set colors
col_fun <- colorRamp2(c(-5, 0, 5), 
                      c("#2166AC", "white", "#B2182B"))


# make annotation
left_annot <- HeatmapAnnotation(
        Workflows = annot_df$processing_method,
        which = 'row',
        col = list(Workflows = workflow_colors), 
        annotation_name_gp = grid::gpar(fontsize = 12),
        annotation_legend_param = list(
            Workflows = list(title_gp = gpar(fontsize = 12, fontface = "bold"), 
                                    labels_gp = gpar(fontsize = 12))))


# left_annot <- HeatmapAnnotation(
#         Workflows = annot_df$processing_method,
#         Cell_types = annot_df$cell_type,
#         which = 'row',
#         col = list(Workflows = workflow_colors,
#                 Cell_types = cell_colors), 
#         annotation_name_gp = grid::gpar(fontsize = 12),
#         annotation_legend_param = list(
#             Workflows = list(title_gp = gpar(fontsize = 12, fontface = "bold"), 
#                                     labels_gp = gpar(fontsize = 12)),
#             Cell_types = list(title_gp = gpar(fontsize = 12, fontface = "bold"), 
#                                     labels_gp = gpar(fontsize = 12))))


# annotate selected genes 
genes_to_plot <- de_res %>% 
    group_by(cluster) %>%
    filter(p_val_adj < 0.05) %>%
    slice_max(avg_log2FC, n = 5) %>% 
    pull(gene)

# make sure the selected markers are in the matrix
all_markers <- genes_to_plot[genes_to_plot %in% rownames(scaled_mtx)]

# get location of the genes to be annotated
mtx_index <- seq(1:nrow(scaled_mtx))
names(mtx_index) <- rownames(scaled_mtx)
position_in_matrix <- mtx_index[names(mtx_index) %in% all_markers]

bottom_annotation <- HeatmapAnnotation(
    markers = anno_mark(
        at = position_in_matrix, 
        labels = names(position_in_matrix),
        which = "column", side = "bottom",
        labels_gp = gpar(fontsize = 8)))


# plot heatmap ----
plot <- Heatmap(t(scaled_mtx),
                name = 'Scaled mean expression',
                heatmap_legend_param = list(title_gp = gpar(fontsize = 12, fontface = "bold")),
                col = col_fun,
                left_annotation = left_annot,
                bottom_annotation = bottom_annotation,
                show_column_names= FALSE,
                show_row_names = TRUE,
                cluster_rows = FALSE,
                cluster_columns = TRUE,
                split = annot_df$processing_method,
                column_names_gp = gpar(fontsize = 6),
                row_names_gp = gpar(fontsize = 8))


pdf(file = paste0(paper_fig_dir, '/', 'top_DE_gene_heatmap', '.pdf'),
    width = 16, height = 7)
    draw(plot)
dev.off()
```
