## Supplementary material for "snPATHO-seq: unlocking the pathology archives": scripts: 04_inferCNV.R

```
# aim ---- 
    # run inferCNV to identify malignant cells


# define environment ----
.libPaths(
    c(
        '/share/ScratchGeneral/tonwan/projects/snPATHO-Seq/renv/library/R-4.1/x86_64-conda-linux-gnu',
        .libPaths()
    )
)
library(Seurat)
library(tidyverse)
library(infercnv)


# arguments ----
obj_file_path <- '/share/ScratchGeneral/tonwan/projects/snPATHO-Seq/data/external_data/10X_data_annotated/melanoma_7167A_annotated.rds'

batch_col_name <- 'processing_method'
batch <- 'snPATHO'
# snPATHO scFFPE

initial_out_dir <- '/share/ScratchGeneral/tonwan/projects/snPATHO-Seq/results/snRNA_external_data/10X_data_Tony/CNV/melanoma_7167A/melanocyte_only'
dir.create(initial_out_dir, recursive = T)

out_dir <- paste0(initial_out_dir, '/', batch)
dir.create(out_dir, recursive = T)


cell_type_col_name <- 'initial_annotation'
cell_types_to_exclude <- c('unknown', 'RBCs', 'Cycling', 'Unassigned', 'Keratinocytes')

gene_order_file <- '/share/ScratchGeneral/tonwan/projects/snPATHO-Seq/resources/Javier_gene_ordering_hg38_gencode_v27.txt'

ref_cell_types <- c('CAF', 'Macrophage', 'PVL', 'T_cycling', 'Neuron', 'RGS5+_PVL', 'Endothelial', 
    'Lymphatic_endothelial', 'Monocyte', 'B', 'T_CD4', 'T_CD8', 'Plasmablasts', 'Macrophages', 'SMCs',
    'CAF', 'Monocytes', 'pDC', 'Pericytes', 'Fibroblasts', 'Mixed_immune_cells', 'vSMC', 'Other_macrophages',
    'Stellate_cells', 'Kupffer_cells', 'Portal_fibroblasts', 'Other_endothelial', 'LSEC', 'NK', 'T', 'NKT',
    'Monocytes', 'Mast_cells', 'Myeloid', 'CAF_PVL')

# sd amplifer for denoising
sd_amplifer <- 1.5


# load the data ----
obj <- readRDS(file = paste0(obj_file_path))

# only keep the batch of interest ----
subset_data <- subset(obj, subset = !!sym(batch_col_name) == batch)

# remove unwanted cell types ----
subset_data <- subset(subset_data, subset = !!sym(cell_type_col_name) %in% cell_types_to_exclude, invert = TRUE)


# get count matrix ----
mtx <- GetAssayData(subset_data, assay = 'RNA', slot = 'counts')

# make sure the reference cell types are present in the dataset
ref_cell_types <- ref_cell_types[ref_cell_types %in% unique(subset_data@meta.data[[cell_type_col_name]])]

# make an annotation file
annotation_df <- data.frame(barcodes = rownames(subset_data@meta.data),
  annotation = subset_data@meta.data[cell_type_col_name])
write.table(annotation_df, file = paste0(out_dir, '/', 'annotation_input.tsv'),
  quote = FALSE, row.names = FALSE, col.names = FALSE, sep = "\t")
  
# make an infercnv object
infercnv_obj = CreateInfercnvObject(
    raw_counts_matrix=mtx,
    gene_order_file=gene_order_file,
    annotations_file=paste0(out_dir, '/', 'annotation_input.tsv'),
    ref_group_names=ref_cell_types,
    delim="\t")


# run infercnv 
infercnv_obj_run <- infercnv::run(
    infercnv_obj = infercnv_obj,
    cutoff = 0.1, #this is the threshol recommeded for Chromium. Even though Visium data is more sparse than sc data, this cutoff was used for the work on siCNVs in prostate cancer
    sd_amplifier = sd_amplifer,
    out_dir = out_dir,
    cluster_by_groups = FALSE,
    cluster_references = TRUE,
    HMM = TRUE,
    denoise=TRUE,
    plot_steps=FALSE,
    png_res = 300) 


# save the results as an RDS file
saveRDS(infercnv_obj_run, file = paste0(out_dir, '/', 'infercnv_obj.rds')) 
```
