## Supplementary material for "snPATHO-seq: unlocking the pathology archives": scripts: 01_integration.R

```
# aim ----
    # integrated cellbender filtered data from 4066


# define environment ----
library(here)
setwd(here())

# make sure renv is loaded
source(paste0(here(), '/', '.Rprofile'))
print(.libPaths())

# load other packages
library(Seurat)
library(tidyverse)
library(RColorBrewer)


# arguments ----
data_dir <- '/share/ScratchGeneral/tonwan/projects/snPATHO-Seq/data/snPATHO_seq/25k_downsampled_processed_objs/cellbender_filtered/seurat_objs'

sample_id <- '4066'

out_dir <- '/share/ScratchGeneral/tonwan/projects/snPATHO-Seq/results/integration/25k_downsampled/cellbender'

out_data_dir <- '/share/ScratchGeneral/tonwan/projects/snPATHO-Seq/data/snPATHO_seq/25k_downsampled_processed_objs/integrated'


########################################################################################################################
########################################################################################################################
########################################################################################################################
working_dir <- paste0(out_dir, '/', sample_id)
dir.create(working_dir, recursive = T)


# load data ----
ffpe_obj <- readRDS(file = paste0(data_dir, '/', sample_id, 'FFPE', '.rds'))
sf_fix_obj <- readRDS(file = paste0(data_dir, '/', sample_id, 'SNAPFix', '.rds'))
gex_obj <- readRDS(file = paste0(data_dir, '/', sample_id, 'GEX', '.rds'))


# save data into a list for easier processing ----
obj_list <- list(ffpe_obj, sf_fix_obj, gex_obj)

names(obj_list) <- c('FFPE_snPATHO', 'SNAP_snPATHO', 'SNAP_3p')


# basic filtering of low quality cells ----
filtered_obj_list <- lapply(obj_list, function(x){
    x <- subset(x, subset = nFeature_RNA > 200)
    x <- subset(x, subset = nFeature_RNA < 8000)
    x[["percent.mt"]] <- PercentageFeatureSet(x, pattern = "^MT-")
    x <- subset(x, subset = percent.mt < 10)
})


########################################################################################################################
########################################################################################################################
########################################################################################################################
# identify and remove doublet from the dataset ----
db_removal_dir <- paste0(working_dir, '/', 'doublet_removal')
dir.create(db_removal_dir, recursive = T)

source('/share/ScratchGeneral/tonwan/projects/snPATHO-Seq/scripts/function_scripts/05_doublet_finder.R')

# identify doublet
doublet_identified_obj_list <- lapply(filtered_obj_list, function(x){
    output_label <- unique(x$processing_method)
    x <- DB_finder_removal(x,
                        expected_doublet_rate = 0.08,
                        out_dir = db_removal_dir,
                        output_name = output_label,
                        output_col_name = 'doublet_status')
    return(x)
})


# remove the cells
doublet_filtered_obj_list <- lapply(doublet_identified_obj_list, function(x){
    Idents(x) <- 'doublet_status'
    x <- subset(x, idents = 'Singlet')
})


####################################################################################################
####################################################################################################
####################################################################################################
# integrate data
# load function
source('/share/ScratchGeneral/tonwan/projects/snPATHO-Seq/scripts/function_scripts/01_Seurat_integration.R')


# integrate data 
integrated_obj <- seurat_integration(doublet_filtered_obj_list,
                                    normalisation_method = 'RNA',
                                    nfeatue_each_dataset = 2000,
                                    nfeatures_common = 2000,
                                    vars_to_regress = NULL,
                                    integration_method = 'cca',
                                    nthread = NULL)


####################################################################################################
####################################################################################################
####################################################################################################
# check integration quality and annotate clusters
integrated_obj$cells_with_low_gene_count <- ifelse(integrated_obj$nFeature_RNA < 500, 'yes', 'no')
integrated_obj$cells_with_high_gene_count <- ifelse(integrated_obj$nFeature_RNA > 5000, 'yes', 'no')

# plot check batch factors and QC values
pdf(paste0(working_dir, '/', 'umap_batch_factors.pdf'))
    print(
        DimPlot(integrated_obj, reduction = "umap", group.by = 'processing_method')
    )
    print(
        FeaturePlot(integrated_obj, features = "nCount_RNA", reduction = 'umap') +
            scale_color_continuous(trans='log2', type = 'viridis')
    )
    print(
        FeaturePlot(integrated_obj, features = "nFeature_RNA", reduction = 'umap') +
            scale_color_continuous(trans='log2', type = 'viridis')
    )
    print(
        FeaturePlot(integrated_obj, features = "percent.mt", reduction = 'umap')
    )
    print(
        DimPlot(integrated_obj, reduction = "umap", group.by = 'cells_with_low_gene_count')
    )
    print(
        DimPlot(integrated_obj, reduction = "umap", group.by = 'cells_with_high_gene_count')
    )
dev.off()


# split umap by sample
pdf(file = paste0(working_dir, '/', 'umap_split_by_processing_method.pdf'),
    width = 15,
    height = ceiling(length(unique(integrated_obj$sample_id)) / 3) * 5)
    print(
        DimPlot(integrated_obj, reduction = "umap", group.by = 'processing_method', 
            split.by = 'processing_method', ncol = 3)
    )
dev.off()


# plot markers ----
marker_dir <- paste0(working_dir, '/', 'markers')
dir.create(marker_dir, recursive = T)

markers <- c(
    'EPCAM', 'ESR1', 'ERBB2', 'KRT8', 'KRT18', 'KRT5', 'KRT14', 'KIT', 'GATA3', 'KRT19', 'MYH11', 'MYLK', 'MKI67',
    'PTPRC', 'CD3D', 'CD3E', 'CD4', 'CD8A', 'TRAC', 'FOXP3', 'TRBC1', 'TRDC', 'TRGC1', 'TRGC2',
    'MS4A1', 'CD79A', 'CD19', 'IGHM', 'IGHG1', 'IGHA',
    'CD14', 'CD68', 'ITGAM', 'CCR7', 'S100A8', 'S100A9', 'EGR1', 'FCN1', 'FCGR3A', 'FCER1A', 'CST3',
    'COL1A1', 'PDGFRB', 'PDGFRA', 'PDPN', 'ICAM1', 'CXCL12',
    'CLEC9A', 'ITGAE', 'THBD', 'XCR1', 'ITGAX', 
    'CD1C', 'CD207', 'ITGAM', 'NOTCH2', 'SIRPA',
    'MCAM', 'ACTA2', 'RGS5',
    'PECAM1', 'FLT1', 'VWF', 'CD34', 'CLEC4G', 'PROX1', 'LYVE1',
    'DEFA4', 'DEFA3', 'MMP8', 'DEFA1', 'CEACAM6', 'CEACAM8', 'LTF', 'MPO', 'ARG1', 'MSHA3', 'DAAM2',
    'NKG7', 'NCAM1',
    'TUBB3',
    'PPBP', 'TUBB1',
    'HDC', 'CLC', 'IL3RA', 'FUT4', 'FCGR3A', 'CD33', 'PI3', 'CHI3L1', 'CXCL8', 'ANXA3', 'TGM3',
    'ADAMTS3', 'CPA3', 'CMA1', 'CTSG', 'ARHGAP15', 'CPM', 'FCN1', 'FTL', 'HSPA6', 'ITGA9', 'RNASE3',
    'S100A4', 'SIGLEC8', 'SLC6A4', 'PTGS2', 'EGR3', 'PILRA',
    'ZEB1', 'ZEB2', 'TGFB1', 'CXCL12', 'VIM', 'FN1', 
    'IFITM1', 'IFITM2', 'IFITM3', 'IRF1', 'B2M', 'HLA-A', 'HLA-B',
    'PKD1', 'PKD2', 'PKHD1', 'FGF19', 'FGF21', 'FGF23', 'FGFR1', 'FGFR2', 'FGFR3', 'FGFR4')


DefaultAssay(integrated_obj) <- 'RNA'
to_plot <- markers[markers %in% rownames(integrated_obj)]
for (g in to_plot) {
    pdf(paste0(marker_dir, '/', g, '.pdf'))
        print(FeaturePlot(integrated_obj, features = g, reduction = 'umap', order = T))
    dev.off()
}
 


####################################################################################################
####################################################################################################
####################################################################################################
# find optimal clustering resolution ----
clust_dir <- paste0(working_dir, '/', 'cluster_optimisation')
dir.create(clust_dir, recursive = T)

# load function
source('/share/ScratchGeneral/tonwan/projects/snPATHO-Seq/scripts/function_scripts/02_Seurat_clustering_optimisation.R')

# make sure the default assay is the integrated assay
DefaultAssay(integrated_obj) <- 'integrated'

# cluster and find optimal clustering resolution
integrated_obj <- clustering_resolution_optimisation(integrated_obj,
                                        resolution_range = seq(0.2, 2, by = 0.1),
                                        plot_dir = clust_dir,
                                        batch_factor = "processing_method",
                                        run_silhouette = FALSE,
                                        dist.matrix = NULL)


####################################################################################################
####################################################################################################
####################################################################################################
# save processed data
saveRDS(file = paste0(out_data_dir, '/', sample_id, '_cellbender_filtered_integration.rds'), integrated_obj)
```
