## Supplementary material for "snPATHO-seq: unlocking the pathology archives": scripts: 02_cell_type_annotation_0.4.R

```
# aim ----
    # annotate cell types in 4066


# define environment ----
library(Seurat)
library(tidyverse)
library(RColorBrewer)


# arguments ----
data_dir <- '/share/ScratchGeneral/tonwan/projects/snPATHO-Seq/data/snPATHO_seq/25k_downsampled_processed_objs/integrated'
obj_file <- '4066_cellbender_filtered_integration.rds'

sample_id <- '4066'

out_dir <- '/share/ScratchGeneral/tonwan/projects/snPATHO-Seq/results/integration/25k_downsampled/cellbender'


# load data ----
integrated_obj <- readRDS(file = paste0(data_dir, '/', obj_file))


# check the expression of selected cell type markers 
marker_dir <- paste0(out_dir, '/', sample_id, '/', 'markers')
dir.create(marker_dir, recursive = T)

markers <- c('APOE', 'TREM2', 'SPP1',
    'IL3RA', 'TLR7', 'TLR9',
    'CD274', 'PDCD1LG2', 'CD5', 'CD163',
    'CLEC10A', 'EREG')


DefaultAssay(integrated_obj) <- 'RNA'
to_plot <- markers[markers %in% rownames(integrated_obj)]
for (g in to_plot) {
    pdf(paste0(marker_dir, '/', g, '.pdf'))
        print(FeaturePlot(integrated_obj, features = g, reduction = 'umap', order = T))
    dev.off()
}


Navin_lum_sec <- c('KRT17', 'LTF', 'KRT23', 'SLPI', 'KRT15')
pdf(paste0(marker_dir, '/', deparse(substitute(Navin_lum_sec)), '.pdf'))
    for (g in Navin_lum_sec) {
        if (g %in% c(rownames(integrated_obj))) {
            print(FeaturePlot(integrated_obj, features = g, reduction = 'umap', order = T))
        }
    }
dev.off()


Navin_lum_HR <- c('AZGP1', 'AGR2', 'ANKRD30A', 'AR', 'ESR1', 'PGR')
pdf(paste0(marker_dir, '/', deparse(substitute(Navin_lum_HR)), '.pdf'))
    for (g in Navin_lum_HR) {
        if (g %in% c(rownames(integrated_obj))) {
            print(FeaturePlot(integrated_obj, features = g, reduction = 'umap', order = T))
        }
    }
dev.off()


Navin_sp_brca_atlas <- c('ADH1B', 'CD36', 'PLIN1', 'PLIN4', 'ADIPOQ', 'FABP4', 'LEP', 'LPL')

pdf(paste0(marker_dir, '/', deparse(substitute(Navin_sp_brca_atlas)), '.pdf'))
    for (g in Navin_sp_brca_atlas) {
        if (g %in% c(rownames(integrated_obj))) {
            print(FeaturePlot(integrated_obj, features = g, reduction = 'umap', order = T))
        }
    }
dev.off()


Bindea_Mast <- c('CMA1', 'MS4A2', 'TPSAB1', 'ABCC4', 'ADCYAP1', 'CALB2', 'CEACAM8', 'CPA3', 'CTSG', 'ELA2',
    'GATA2', 'HPGD', 'HPGDS', 'KIT', 'LINC01140', 'MAOB', 'MLPH', 'MPO', 'NR0B1', 'PPM1H', 'PRG2', 'PTGS1',
    'HDC', 'SCG2', 'SIGLEC6', 'SLC18A2', 'SLC24A3', 'TAL1', 'TPSB2', 'VWA5A')

pdf(paste0(marker_dir, '/', deparse(substitute(Bindea_Mast)), '.pdf'))
    for (g in Bindea_Mast) {
        if (g %in% c(rownames(integrated_obj))) {
            print(FeaturePlot(integrated_obj, features = g, reduction = 'umap', order = T))
        }
    }
dev.off()

Hao_pDC <- c('TCF4', 'PTGDS', 'CCDC50', 'JCHAIN', 'UGCG', 'PPP1R14B', 'IRF8', 'ITM2C', 'GZMB', 'IRF7', 'PLD4', 
    'SERPINF1', 'C12orf75', 'BCL11A', 'ALOX5AP', 'LILRA4', 'IL3RA', 'APP', 'MZB1', 'TPM2', 'TXN', 'SEC61B', 
    'HERPUD1', 'CLIC3', 'PLAC8', 'SPIB', 'SMPD3', 'LINC00996', 'IRF4', 'TCL1A')

pdf(paste0(marker_dir, '/', deparse(substitute(Hao_pDC)), '.pdf'))
    for (g in Hao_pDC) {
        if (g %in% c(rownames(integrated_obj))) {
            print(FeaturePlot(integrated_obj, features = g, reduction = 'umap', order = T))
        }
    }
dev.off()


Dutertre_cDC1 <- c('CLEC9A', 'C1orf54', 'CADM1', 'TMEM14A', 'XCR1', 'HLA-DOB', 'RAB7B', 
    'SLAMF7', 'SHTN1', 'CLNK', 'HLA-DQB2', 'HLA-DQA2', 'CPNE3', 'IDO1', 'WDFY4', 'TPD52', 
    'WFDC21P', 'HLA-DPA1', 'CD74', 'HLA-DRB1', 'HLA-DRB6', 'HLA-DPB1', 'HLA-DQA1', 
    'HLA-DRB5', 'HLA-DQB1', 'CAMK2D', 'HLA-DRA', 'FLT3', 'CPVL', 'RGS10')

pdf(paste0(marker_dir, '/', deparse(substitute(Dutertre_cDC1)), '.pdf'))
    for (g in Dutertre_cDC1) {
        if (g %in% c(rownames(integrated_obj))) {
            print(FeaturePlot(integrated_obj, features = g, reduction = 'umap', order = T))
        }
    }
dev.off()


Dutertre_cDC2 <- c('PKIB','ARL4C','HLA-DQB2','CD1C','HLA-DPB1','HLA-DRA','HLA-DRB1',
    'HLA-DRB5','HLA-DQB1','HLA-DQA1','CD74','HLA-DMA','HLA-DPA1','GPR183','HLA-DMB','SNHG15',
    'FCER1A','C7orf50','GHRL','ZC3HAV1','AREG','CIITA','CLEC10A','FCGR2C','CCR6','PLSCR1',
    'CD1B','LSP1','PSMD8','CDKN1A')

pdf(paste0(marker_dir, '/', deparse(substitute(Dutertre_cDC2)), '.pdf'))
    for (g in Dutertre_cDC2) {
        if (g %in% c(rownames(integrated_obj))) {
            print(FeaturePlot(integrated_obj, features = g, reduction = 'umap', order = T))
        }
    }
dev.off()


Hao_cDC1 <- c('HLA-DPB1', 'HLA-DPA1', 'CPVL', 'CLEC9A', 'IRF8', 'C1orf54', 'SNX3', 
    'CPNE3', 'WDFY4', 'BASP1', 'SHTN1', 'IDO1', 'CLNK', 'BATF3', 'CADM1', 'HLA-DQA1', 
    'HLA-DQB1', 'DNASE1L3', 'HLA-DRB1', 'RGS10', 'HLA-DRB5', 'CST3', 'CD74', 'HLA-DMA', 
    'HLA-DRA', 'GSTP1', 'ID2', 'ACTB', 'LGALS2', 'S100B')

pdf(paste0(marker_dir, '/', deparse(substitute(Hao_cDC1)), '.pdf'))
    for (g in Hao_cDC1) {
        if (g %in% c(rownames(integrated_obj))) {
            print(FeaturePlot(integrated_obj, features = g, reduction = 'umap', order = T))
        }
    }
dev.off()


Hao_cDC2_1 <- c('CLEC10A', 'FCER1A', 'HLA-DRA', 'HLA-DQA1', 'HLA-DRB1', 'HLA-DPB1', 
    'HLA-DPA1', 'CST3', 'HLA-DQB1', 'HLA-DRB5', 'CD74', 'HLA-DMA', 'CPVL', 'LYZ', 'LMNA', 
    'HLA-DMB', 'LGALS2', 'CD1C', 'CCDC88A', 'ANXA2', 'CTSZ', 'S100A10', 'JAML', 'ALDH2', 
    'MS4A6A', 'LGALS1', 'GADD45B', 'GPR183', 'AC020656.1', 'EREG')

pdf(paste0(marker_dir, '/', deparse(substitute(Hao_cDC2_1)), '.pdf'))
    for (g in Hao_cDC2_1) {
        if (g %in% c(rownames(integrated_obj))) {
            print(FeaturePlot(integrated_obj, features = g, reduction = 'umap', order = T))
        }
    }
dev.off()


Hao_cDC2_2 <- c('FCER1A', 'HLA-DQA1', 'HLA-DPB1', 'HLA-DPA1', 'HLA-DRA', 'HLA-DRB1', 
    'HLA-DQB1', 'CST3', 'CLEC10A', 'HLA-DRB5', 'CD74', 'CD1C', 'HLA-DMA', 'ENHO', 'HLA-DMB', 
    'AREG', 'CPVL', 'BASP1', 'CCDC88A', 'TUBA1B', 'CTSZ', 'LGALS2', 'HLA-DQA2', 'SAMHD1', 
    'JAML', 'GSTP1', 'LGALS1', 'CIITA', 'ANXA2', 'ACTB')

pdf(paste0(marker_dir, '/', deparse(substitute(Hao_cDC2_2)), '.pdf'))
    for (g in Hao_cDC2_2) {
        if (g %in% c(rownames(integrated_obj))) {
            print(FeaturePlot(integrated_obj, features = g, reduction = 'umap', order = T))
        }
    }
dev.off()


Charoentong_immature_dendritic_cells <- c('ACADM', 'AHCYL1', 'ALDH1A2', 'ALDH3A2', 'ALDH9A1', 
    'ALOX15', 'AMT', 'ARL1', 'ATIC', 'ATP5A1', 'CAPZA1', 'LILRA5', 'RDX', 'RRAGD', 'TACSTD2', 
    'INPP5F', 'RAB38', 'PLAU', 'CSF3R', 'SLC18A2', 'AMPD2', 'CLTB', 'C1orf162')

pdf(paste0(marker_dir, '/', deparse(substitute(Charoentong_immature_dendritic_cells)), '.pdf'))
    for (g in Charoentong_immature_dendritic_cells) {
        if (g %in% c(rownames(integrated_obj))) {
            print(FeaturePlot(integrated_obj, features = g, reduction = 'umap', order = T))
        }
    }
dev.off()


neuron_markers <- c('TUBB3', 'IL12A', 'SOX2', 'NES', 'PCNA', 'DCX', 'CR1', 'DPYSL3', 'CB1')

pdf(paste0(marker_dir, '/', deparse(substitute(neuron_markers)), '.pdf'))
    for (g in neuron_markers) {
        if (g %in% c(rownames(integrated_obj))) {
            print(FeaturePlot(integrated_obj, features = g, reduction = 'umap', order = T))
        }
    }
dev.off()


Wu_LAMP3_DC <- c('BIRC3', 'LAMP3', 'CCR7', 'IL7R')

pdf(paste0(marker_dir, '/', deparse(substitute(Wu_LAMP3_DC)), '.pdf'))
    for (g in Wu_LAMP3_DC) {
        if (g %in% c(rownames(integrated_obj))) {
            print(FeaturePlot(integrated_obj, features = g, reduction = 'umap', order = T))
        }
    }
dev.off()


selected_LAMP3_DC_markers <- c('BASP1', 'SHTN1', 'IDO1', 'BATF3', 'GSTP1', 'HLA−DRA', 'HLA−DMA', 'ID2')

pdf(paste0(marker_dir, '/', deparse(substitute(selected_LAMP3_DC_markers)), '.pdf'))
    for (g in selected_LAMP3_DC_markers) {
        if (g %in% c(rownames(integrated_obj))) {
            print(FeaturePlot(integrated_obj, features = g, reduction = 'umap', order = T))
        }
    }
dev.off()


########################################################################################################################
########################################################################################################################
########################################################################################################################
optimal_clustering_dir <- paste0(out_dir, '/', sample_id, '/', 'optimal_cluster')
dir.create(optimal_clustering_dir, recursive = T)


opt_res <- 0.4
cluster_name <- grep(paste0('res.', opt_res), colnames(integrated_obj@meta.data), value = T)[1]


# make a plot with label at the optimal cluster resolution ----
pdf(file = paste0(optimal_clustering_dir, '/', 'umap_', opt_res, '.pdf'))
    print(
        DimPlot(integrated_obj, reduction = "umap", group.by = cluster_name, label = T, repel = T)
    )
dev.off()


# check markers for selected clusters
Idents(integrated_obj) <- cluster_name

for (cl in unique(Idents(integrated_obj))) {
    de_markrers <- FindMarkers(integrated_obj, ident.1 = cl, test.use = 't')
    write.csv(file = paste0(optimal_clustering_dir, '/', 'Cluster_', cl, '_markers.csv'), de_markrers)
}


# DE between certain clusters 
de_markrers <- FindMarkers(integrated_obj, assay = 'RNA', ident.1 = 0, ident.2 = 8, test.use = 't')
write.csv(file = paste0(optimal_clustering_dir, '/', 'de_res_0_vs_8.csv'), de_markrers)


####################################################################################
####################################################################################
####################################################################################
# annotate clusters ----
Idents(integrated_obj) <- cluster_name

new_annotation_df <- data.frame(cluster_id = unique(as.character(Idents(integrated_obj))),
                                initial_annotation = '')
new_annotation_df[new_annotation_df$cluster_id == '0', 'initial_annotation'] <- 'CAF'
new_annotation_df[new_annotation_df$cluster_id == '1', 'initial_annotation'] <- 'Epithelial_cancer'
new_annotation_df[new_annotation_df$cluster_id == '2', 'initial_annotation'] <- 'Epithelial_cancer'
new_annotation_df[new_annotation_df$cluster_id == '3', 'initial_annotation'] <- 'T_cells'
new_annotation_df[new_annotation_df$cluster_id == '4', 'initial_annotation'] <- 'Epithelial_cancer'
new_annotation_df[new_annotation_df$cluster_id == '5', 'initial_annotation'] <- 'T_cells'
new_annotation_df[new_annotation_df$cluster_id == '6', 'initial_annotation'] <- 'Macrophage'
new_annotation_df[new_annotation_df$cluster_id == '7', 'initial_annotation'] <- 'Myoepithelial'
new_annotation_df[new_annotation_df$cluster_id == '8', 'initial_annotation'] <- 'CAF'
new_annotation_df[new_annotation_df$cluster_id == '9', 'initial_annotation'] <- 'Endothelial'
new_annotation_df[new_annotation_df$cluster_id == '10', 'initial_annotation'] <- 'Epithelial_cancer'
new_annotation_df[new_annotation_df$cluster_id == '11', 'initial_annotation'] <- 'Epithelial_basal'
new_annotation_df[new_annotation_df$cluster_id == '12', 'initial_annotation'] <- 'PVL'
new_annotation_df[new_annotation_df$cluster_id == '13', 'initial_annotation'] <- 'DC_B_cells'
new_annotation_df[new_annotation_df$cluster_id == '14', 'initial_annotation'] <- 'Epithelial_cancer'
new_annotation_df[new_annotation_df$cluster_id == '15', 'initial_annotation'] <- 'Epithelial_luminal'
new_annotation_df[new_annotation_df$cluster_id == '16', 'initial_annotation'] <- 'MAST'


# annotate the object
new_annotations <- new_annotation_df$initial_annotation
names(new_annotations) <- new_annotation_df$cluster_id

integrated_obj <- RenameIdents(integrated_obj, new_annotations)
integrated_obj$initial_annotation <- Idents(integrated_obj)


# export cell type annotations as csv file for easier modification and assessemnt
write.csv(file = paste0(optimal_clustering_dir, '/', 'initial_annotation.csv'), integrated_obj$initial_annotation)


# plot a umap with new annotations
Idents(integrated_obj) <- 'initial_annotation'
pdf(file = paste0(optimal_clustering_dir, '/', 'umap_initial_annotation.pdf'), width = 10, height = 7)
    print(
        DimPlot(integrated_obj, reduction = "umap", group.by = 'initial_annotation', label = T, repel = T)
    )
dev.off()


####################################################################################
####################################################################################
####################################################################################
# save annotated data
saveRDS(file = paste0(data_dir, '/', sample_id, '_cellbender_filtered_integration_initial_annotation.rds'), integrated_obj)
```
