## Supplementary material for "snPATHO-seq: unlocking the pathology archives": scripts: 03_subcluster_myeloid_and_B_cells.R

```
# aim ----
    # subcluster myeloid cells


# define environment ----
library(Seurat)
library(tidyverse)
library(RColorBrewer)


# arguments ----
obj_file_path <- '/share/ScratchGeneral/tonwan/projects/snPATHO-Seq/data/snPATHO_seq/25k_downsampled_processed_objs/integrated/4066_cellbender_filtered_integration_initial_annotation.rds'

sample_id <- '4066'

out_dir <- '/share/ScratchGeneral/tonwan/projects/snPATHO-Seq/results/integration/25k_downsampled/cellbender'


# load data ----
obj <- readRDS(file = obj_file_path)

subset_obj <- subset(obj, subset = initial_annotation %in% c('DC_B_cells'))


# split data by sample id
obj_list <- list()
for (ds in unique(subset_obj$sample_id)) {
    obj_list[[ds]] <- subset(subset_obj, subset = sample_id == ds)
}


####################################################################################################
####################################################################################################
####################################################################################################
# redo integration
# load function
source('/share/ScratchGeneral/tonwan/projects/snPATHO-Seq/scripts/function_scripts/01_Seurat_integration.R')


# integrate data 
integrated_obj <- seurat_integration(obj_list,
                                    normalisation_method = 'RNA',
                                    nfeatue_each_dataset = 500,
                                    nfeatures_common = 500,
                                    vars_to_regress = NULL,
                                    integration_method = 'cca',
                                    nthread = NULL)


####################################################################################################
####################################################################################################
####################################################################################################
# check integration quality and annotate clusters
working_dir <- paste0(out_dir, '/', sample_id, '/', 'subclustering', '/', 'myeloid_reintegrate')
dir.create(working_dir, recursive = T)

# plot check batch factors and QC values
pdf(paste0(working_dir, '/', 'umap_batch_factors.pdf'))
    print(
        DimPlot(integrated_obj, reduction = "umap", group.by = 'sample_id')
    )
    print(
        FeaturePlot(integrated_obj, features = "nCount_RNA", reduction = 'umap') +
            scale_color_continuous(trans='log2', type = 'viridis')
    )
    print(
        FeaturePlot(integrated_obj, features = "nFeature_RNA", reduction = 'umap') +
            scale_color_continuous(trans='log2', type = 'viridis')
    )
    print(
        FeaturePlot(integrated_obj, features = "percent.mt", reduction = 'umap')
    )
dev.off()


# split umap by sample
pdf(file = paste0(working_dir, '/', 'umap_split_by_sample_id.pdf'),
    width = 5 * length(unique(integrated_obj$sample_id)),
    height = 5)
    print(
        DimPlot(integrated_obj, reduction = "umap", group.by = 'sample_id', 
            split.by = 'sample_id', ncol = 3)
    )
dev.off()


####################################################################################################
####################################################################################################
####################################################################################################
# plot some markers 
marker_dir <- paste0(working_dir, '/', 'markers')
dir.create(marker_dir, recursive = T)

cDC1_markers <- c('CLEC9A', 'IRF8', 'CADM1')
cDC2_markers <- c('CD1C', 'FCER1A', 'CD1E', 'CCL17', 'HLA-DQA1', 'CLEC10A', 'ENHO', 'PLD4', 'GSN', 'SLC38A1', 'NDRG2', 'AFF3')
cDC3_markers <- c('BIRC3', 'CCR7', 'IL7R', 'LAMP3')
pDC_markers <- c('JCHAIN', 'GZMB', 'IRF7')

b_cells <- c('MS4A1', 'CD79A', 'CD19', 'IGHM', 'IGHA1', 'IGHA2', 'IGHG1', 'IGHD', 'TNFRSF17')

b_intermediate <- c('MS4A1', 'TNFRSF13B', 'IGHM', 'IGHD', 'AIM2', 'CD79A', 'LINC01857', 'RALGPS2', 'BANK1', 'CD79B')
plasmablast <- c('IGHA2', 'MZB1', 'TNFRSF17', 'DERL3', 'TXNDC5', 'TNFRSF13B', 'POU2AF1', 'CPNE5', 'HRASLS2', 'NT5DC2')

other_markers <- c('ERBB2', 'MT-CO3', 'KRT8', 'KRT18')

markers <- c(cDC1_markers, cDC2_markers, cDC3_markers, pDC_markers, 
    b_cells, b_intermediate, plasmablast,
    'MKI67', 'CD68', 'CD14', other_markers)


DefaultAssay(integrated_obj) <- 'RNA'
to_plot <- markers[markers %in% rownames(integrated_obj)]
for (g in to_plot) {
    pdf(paste0(marker_dir, '/', g, '.pdf'))
        print(FeaturePlot(integrated_obj, features = g, reduction = 'umap', order = T))
    dev.off()
}
 


####################################################################################################
####################################################################################################
####################################################################################################
# find optimal clustering resolution ----
clust_dir <- paste0(working_dir, '/', 'cluster_optimisation')
dir.create(clust_dir, recursive = T)

# load function
source('/share/ScratchGeneral/tonwan/projects/snPATHO-Seq/scripts/function_scripts/02_Seurat_clustering_optimisation.R')

# make sure the default assay is the integrated assay
DefaultAssay(integrated_obj) <- 'integrated'

# get distance matrix for silhouette score calcualtion
dist.matrix <- dist(x = Embeddings(integrated_obj, reduction = "pca"))

# cluster and find optimal clustering resolution
integrated_obj <- clustering_resolution_optimisation(integrated_obj,
                                        resolution_range = seq(0.2, 2, by = 0.1),
                                        plot_dir = clust_dir,
                                        batch_factor = "sample_id",
                                        run_silhouette = TRUE,
                                        dist.matrix = dist.matrix)


####################################################################################################
####################################################################################################
####################################################################################################
# annotate subclusters
optimal_clustering_dir <- paste0(working_dir, '/', 'optimal_cluster')
dir.create(optimal_clustering_dir, recursive = T)


opt_res <- 0.3
cluster_name <- grep(paste0('res.', opt_res), colnames(integrated_obj@meta.data), value = T)[1]


# make a plot with label at the optimal cluster resolution ----
pdf(file = paste0(optimal_clustering_dir, '/', 'umap_', opt_res, '.pdf'))
    print(
        DimPlot(integrated_obj, reduction = "umap", group.by = cluster_name, label = T, repel = T)
    )
dev.off()


####################################################################################
####################################################################################
####################################################################################
# annotate clusters ----
Idents(integrated_obj) <- cluster_name
annotation_col_name <- 'new_DC_B_annotation'

new_annotation_df <- data.frame(cluster_id = unique(as.character(Idents(integrated_obj))))
new_annotation_df[[annotation_col_name]] <- ''

new_annotation_df[new_annotation_df$cluster_id == '0', annotation_col_name] <- 'B_cells'
new_annotation_df[new_annotation_df$cluster_id == '1', annotation_col_name] <- 'B_cells'
new_annotation_df[new_annotation_df$cluster_id == '2', annotation_col_name] <- 'DC'
new_annotation_df[new_annotation_df$cluster_id == '3', annotation_col_name] <- 'DC'
new_annotation_df[new_annotation_df$cluster_id == '4', annotation_col_name] <- 'DC'


# annotate the object
new_annotations <- new_annotation_df[[annotation_col_name]]
names(new_annotations) <- new_annotation_df$cluster_id

integrated_obj <- RenameIdents(integrated_obj, new_annotations)
integrated_obj[[annotation_col_name]] <- Idents(integrated_obj)


# export cell type annotations as csv file for easier modification and assessemnt
write.csv(file = paste0(optimal_clustering_dir, '/', annotation_col_name, '.csv'), integrated_obj[[annotation_col_name]])


# plot a umap with new annotations
Idents(integrated_obj) <- annotation_col_name
pdf(file = paste0(optimal_clustering_dir, '/', 'umap_', annotation_col_name, '.pdf'), width = 10, height = 7)
    print(
        DimPlot(integrated_obj, reduction = "umap", group.by = annotation_col_name, label = T, repel = T)
    )
dev.off()
```
