## Supplementary material for "snPATHO-seq: unlocking the pathology archives": scripts: 04_finalise_annotation.R

```
# aim ----
    # update annotation based on subclustering


# define environment ----
library(Seurat)
library(tidyverse)


# arguments ----
data_obj_file_path <- '/share/ScratchGeneral/tonwan/projects/snPATHO-Seq/data/snPATHO_seq/25k_downsampled_processed_objs/integrated/4066_cellbender_filtered_integration_initial_annotation.rds'

out_dir <- '/share/ScratchGeneral/tonwan/projects/snPATHO-Seq/results/integration/25k_downsampled/cellbender/4066/final_annotation'
dir.create(out_dir, recursive = TRUE)


#####################################################################################################################
#####################################################################################################################
#####################################################################################################################
# load data ----
obj <- readRDS(file = data_obj_file_path)


new_annotations <- read.csv(file = '/share/ScratchGeneral/tonwan/projects/snPATHO-Seq/results/integration/25k_downsampled/cellbender/4066/subclustering/myeloid_reintegrate/optimal_cluster/new_DC_B_annotation.csv',
    row.names = 1)
colnames(new_annotations) <- c('new_annotation')

new_meta_data <- base::merge(obj@meta.data, new_annotations, by = 0, all.x = TRUE)
new_meta_data$new_annotation <- ifelse(is.na(new_meta_data$new_annotation), as.character(new_meta_data$initial_annotation), new_meta_data$new_annotation)
new_meta_data <- column_to_rownames(new_meta_data, var = 'Row.names')
new_meta_data <- new_meta_data[rownames(obj@meta.data), ]


obj$major_annotation <- new_meta_data$new_annotation


#####################################################################################################################
#####################################################################################################################
#####################################################################################################################
# save object with final annotation
saveRDS(file = '/share/ScratchGeneral/tonwan/projects/snPATHO-Seq/data/snPATHO_seq/25k_downsampled_processed_objs/integrated/4066_cellbender_filtered_integration_final_annotation.rds', obj)
```
