## Supplementary material for "snPATHO-seq: unlocking the pathology archives": scripts: 05_paper_figures.R

```
# aim ----
    # polish plots for paper


# load other packages
library(Seurat)
library(tidyverse)
library(RColorBrewer)

library(ComplexHeatmap)
library(circlize)


# arguments ----
data_obj_file_path <- '/share/ScratchGeneral/tonwan/projects/snPATHO-Seq/data/snPATHO_seq/25k_downsampled_processed_objs/integrated/4066_cellbender_filtered_integration_final_annotation.rds'

out_dir <- '/share/ScratchGeneral/tonwan/projects/snPATHO-Seq/results/integration/25k_downsampled/cellbender/4066/final_annotation'


# load data ----
obj <- readRDS(file = data_obj_file_path)


# rename processing method to make the naming more consistent
obj$processing_method <- ifelse(obj$processing_method == 'FFPE_snPATHO', 'snPATHO',
    ifelse(obj$processing_method == 'SNAP_snPATHO', 'FLEX', '3p'))


# load predefined colors
source('/share/ScratchGeneral/tonwan/projects/snPATHO-Seq/scripts/colors.R')

# only keep colors for cell types present
cell_type_colors <- cell_type_colors[names(cell_type_colors) %in% unique(obj$major_annotation)]
cell_type_colors <- cell_type_colors[order(names(cell_type_colors))]

# snRNA-seq workflow colors
workflow_colors <- workflow_colors[order(names(workflow_colors))]


########################################################################################################################
########################################################################################################################
########################################################################################################################
# plot umap with modified cell type colors
plot <- DimPlot(obj, group.by = 'major_annotation', cols = cell_type_colors) +
            labs(title = NULL)
plot <- LabelClusters(plot = plot, id = "major_annotation", repel = TRUE, box = TRUE, force = 1, color = 'white')

pdf(file = paste0(out_dir, '/', 'umap_high_level_annotation_paper_figure.pdf'), width = 8)
    print(
        plot
    )     
dev.off()


pdf(file = paste0(out_dir, '/', 'umap_split_by_sample_paper_figure.pdf'),
    width = 15,
    height = 5)
    print(
        DimPlot(obj, reduction = "umap", group.by = 'processing_method', 
            split.by = 'processing_method', ncol = 3, cols = workflow_colors) +
            labs(title = NULL)
    )
dev.off()


########################################################################################################################
########################################################################################################################
########################################################################################################################
# run DE between annotated cell types
Idents(obj) <- 'major_annotation'
de_res <- FindAllMarkers(obj, assay = 'RNA', test.use = 't', only.pos = TRUE)
write.csv(file = paste0(out_dir, '/', 'de_between_major_cell_types.csv'), de_res)


# plot expression of top DE genes between clusters - annotate canonical cell type markers
# get top 200 DE genes from each cluster to plot
top_markers <- de_res %>% 
    group_by(cluster) %>%
    filter(p_val_adj < 0.05) %>%
    slice_max(avg_log2FC, n = 200) %>% 
    pull(gene)


# get mean expresion of the selected genes for each processing method 
plot_mtx <- as.data.frame(AverageExpression(obj,assays = 'RNA', slot = 'data', features = top_markers, group.by = c('sample_id', 'major_annotation'))[[1]])
plot_mtx$gene_names <- rownames(plot_mtx)
plot_mtx_long <- gather(plot_mtx, key = 'annotation', value = 'mean_expression', -gene_names)

# annotate the expression dataframe
plot_mtx_long$processing_method <- 
    ifelse(plot_mtx_long$annotation %in% grep('FFPE', plot_mtx_long$annotation, value = TRUE, ignore.case = TRUE), 'snPATHO', 
    ifelse(plot_mtx_long$annotation %in% grep('GEX', plot_mtx_long$annotation, value = TRUE, ignore.case = TRUE), '3p',
    ifelse(plot_mtx_long$annotation %in% grep('SNAPfix', plot_mtx_long$annotation, value = TRUE, ignore.case = TRUE), 'FLEX', 'unlabelled')))
plot_mtx_long$cell_type <- unlist(lapply(str_split(plot_mtx_long$annotation, '\\_'), function(x) paste(x[-1], collapse = '_')))


# scale the gene expression within each processing method
plot_mtx_long <- plot_mtx_long %>% 
        group_by(gene_names, processing_method) %>%
        mutate(scale(mean_expression)) %>% 
        as.data.frame() 
  
plot_order <- plot_mtx_long %>% 
    arrange(factor(cell_type, levels = c('Epithelial_luminal', 'Epithelial_cancer', 'PVL', 'Epithelial_basal', 
        'Endothelial', 'CAF', 'Myoepithelial',  'DC', 'B_cells', 'Macrophage', 'MAST', 'T_cells'))) %>% 
    pull(annotation) %>% 
    unique()


scaled_mtx <- spread(plot_mtx_long[, c('gene_names', 'annotation', 'scale(mean_expression)')],
        value = 'scale(mean_expression)', key = 'annotation')
scaled_mtx <- column_to_rownames(scaled_mtx, var = 'gene_names')
# cap the scaled expression at -5 to 5
scaled_mtx[scaled_mtx > 5] <- 5
scaled_mtx[scaled_mtx < -5] <- -5
scaled_mtx <- scaled_mtx[rowSums(is.na(scaled_mtx)) == 0, ] 
scaled_mtx <- scaled_mtx[, plot_order]

# make heatmap annotation
annot_df <- plot_mtx_long[, c('annotation', 'processing_method', 'cell_type')]
annot_df <- distinct(annot_df)
annot_df <- annot_df[match(colnames(scaled_mtx), annot_df$annotation), ]


# set colors
col_fun <- colorRamp2(c(-5, 0, 5), 
                      c("#2166AC", "white", "#B2182B"))

# set annotation colors & make annotation ----
source('/share/ScratchGeneral/tonwan/projects/snPATHO-Seq/scripts/colors.R')
workflow_colors <- workflow_colors[names(workflow_colors) %in% unique(annot_df$processing_method)]
cell_type_colors <- cell_type_colors[names(cell_type_colors) %in% unique(annot_df$cell_type)]

# make annotation
left_annot <- HeatmapAnnotation(
        Workflows = annot_df$processing_method,
        Cell_types = annot_df$cell_type,
        which = 'row',
        col = list(Workflows = workflow_colors,
                Cell_types = cell_type_colors), 
        annotation_name_gp = grid::gpar(fontsize = 12),
        annotation_legend_param = list(
            Workflows = list(title_gp = gpar(fontsize = 12, fontface = "bold"), 
                                    labels_gp = gpar(fontsize = 12)),
            Cell_types = list(title_gp = gpar(fontsize = 12, fontface = "bold"), 
                                    labels_gp = gpar(fontsize = 12))))


# annotate selected genes 
genes_to_plot <- c(
    'EPCAM', 'ERBB2', 'KRT8', 'MKI67',
    'KRT19', 'LTF', 'ESR1',
    'KRT6B', 'KRT5', 'KRT14',
    'MYH11', 'MYLK', 'ACTA2',
    'COL1A1', 'PDGFRA', 'PDGFRB',
    'RGS5', 'MCAM',
    'PECAM1', 'VWF', 'FLT1',
    'TRAC', 'CD3D', 'CD4', 'CD8A', 'FOXP3',
    'MS4A1', 'CD79A', 'CD19', 'JCHAIN',
    'CLEC9A', 'CD1C', 'CCR7',
    'ITGAX', 'CD68', 'CD14', 'FCGR3A', 'CD163', 'SPP1',
    'HDC', 'GATA2', 'CPA3', 'MS4A2'
)

# make sure the selected markers are in the matrix
all_markers <- genes_to_plot[genes_to_plot %in% rownames(scaled_mtx)]

# get location of the genes to be annotated
mtx_index <- seq(1:nrow(scaled_mtx))
names(mtx_index) <- rownames(scaled_mtx)
position_in_matrix <- mtx_index[names(mtx_index) %in% all_markers]

bottom_annotation <- HeatmapAnnotation(markers = anno_mark(at = position_in_matrix, 
        labels = names(position_in_matrix),
        which = "column", side = "bottom"))

# plot heatmap ----
plot <- Heatmap(t(scaled_mtx),
                name = 'Scaled mean expression',
                heatmap_legend_param = list(title_gp = gpar(fontsize = 12, fontface = "bold")),
                col = col_fun,
                left_annotation = left_annot,
                bottom_annotation = bottom_annotation,
                show_column_names= FALSE,
                show_row_names = FALSE,
                cluster_rows = FALSE,
                cluster_columns = TRUE,
                split = annot_df$processing_method,
                column_names_gp = gpar(fontsize = 6),
                row_names_gp = gpar(fontsize = 8))


pdf(file = paste0(out_dir, '/', 'top_DE_gene_canonical_marker_heatmap', '.pdf'),
    width = 14, height = 7)
    draw(plot)
dev.off()
```
