## Supplementary material for "snPATHO-seq: unlocking the pathology archives": scripts: 02_cell_type_annotation_1.1.R

```
# aim ----
    # annotate cell types


# define environment ----
library(Seurat)
library(tidyverse)
library(RColorBrewer)


# arguments ----
data_dir <- '/share/ScratchGeneral/tonwan/projects/snPATHO-Seq/data/snPATHO_seq/25k_downsampled_processed_objs/integrated'
obj_file <- '4399_cellbender_filtered_integration.rds'

sample_id <- '4399'

out_dir <- '/share/ScratchGeneral/tonwan/projects/snPATHO-Seq/results/integration/25k_downsampled/cellbender'


# load data ----
integrated_obj <- readRDS(file = paste0(data_dir, '/', obj_file))


# check the expression of selected cell type markers 
marker_dir <- paste0(out_dir, '/', sample_id, '/', 'markers')
dir.create(marker_dir, recursive = T)

markers <- c('HGF', 'HPX', 'ALB', 'APOE',
    'SPTA1', 'HBB', 'SPTB',
    'ABCC1', 'ALDH1A2', 'HNF1B', 'SOX9', 'ONECUT1', 'ANXA4'
    )

DefaultAssay(integrated_obj) <- 'RNA'
to_plot <- markers[markers %in% rownames(integrated_obj)]
for (g in to_plot) {
    pdf(paste0(marker_dir, '/', g, '.pdf'))
        print(FeaturePlot(integrated_obj, features = g, reduction = 'umap', order = T))
    dev.off()
}


cholangiocyte_markers <- c('ANXA4', 'KRT7', 'DEFB1', 'FXYD2', 'TM4SF4', 'KRT19', 'KRT8', 'TACSTD2', 'EGRI1', 'SPP1')
pdf(paste0(marker_dir, '/', deparse(substitute(cholangiocyte_markers)), '.pdf'))
    for (g in cholangiocyte_markers) {
        if (g %in% c(rownames(integrated_obj))) {
            print(FeaturePlot(integrated_obj, features = g, reduction = 'umap', order = T))
        }
    }
dev.off()


hepatocyte_markers <- c('FABP1', 'CYP2E1', 'HP', 'ORM1', 'APOA2', 'ADH4', 'TTR', 'APOA1', 'APOC1', 'APOC3')
pdf(paste0(marker_dir, '/', deparse(substitute(hepatocyte_markers)), '.pdf'))
    for (g in hepatocyte_markers) {
        if (g %in% c(rownames(integrated_obj))) {
            print(FeaturePlot(integrated_obj, features = g, reduction = 'umap', order = T))
        }
    }
dev.off()


stellate_cell_markers <- c('IGFBP3', 'DCN', 'CXCL14', 'COLEC11', 'BGN', 'CXCL12', 'ANGPTL6', 'TMEM56', 'ECM1', 'COL3A1')
pdf(paste0(marker_dir, '/', deparse(substitute(stellate_cell_markers)), '.pdf'))
    for (g in stellate_cell_markers) {
        if (g %in% c(rownames(integrated_obj))) {
            print(FeaturePlot(integrated_obj, features = g, reduction = 'umap', order = T))
        }
    }
dev.off()


portal_endothelial_markers <- c('CLEC14A', 'TM4SF1', 'ID1', 'PECAM1', 'DLL4', 'MGP', 'SPARCL1', 'SLC9A3R2', 'RAMP2', 'VWF')
pdf(paste0(marker_dir, '/', deparse(substitute(portal_endothelial_markers)), '.pdf'))
    for (g in portal_endothelial_markers) {
        if (g %in% c(rownames(integrated_obj))) {
            print(FeaturePlot(integrated_obj, features = g, reduction = 'umap', order = T))
        }
    }
dev.off()


central_vein_endo_markers <- c('PRSS23', 'SELE', 'RSPO3', 'LXH6', 'RAMP3', 'LIFR', 'LYPD2', 'NTS', 'IFI27', 'DNASE1L3')
pdf(paste0(marker_dir, '/', deparse(substitute(central_vein_endo_markers)), '.pdf'))
    for (g in central_vein_endo_markers) {
        if (g %in% c(rownames(integrated_obj))) {
            print(FeaturePlot(integrated_obj, features = g, reduction = 'umap', order = T))
        }
    }
dev.off()


portal_fibroblast_markers <- c('THY1', 'CCL2', 'PTGDS', 'ELN', 'MFAP4', 'IGFBP7', 'FBLN1', 'COL1A2', 'ACTA2', 'IGFBP5')
pdf(paste0(marker_dir, '/', deparse(substitute(portal_fibroblast_markers)), '.pdf'))
    for (g in portal_fibroblast_markers) {
        if (g %in% c(rownames(integrated_obj))) {
            print(FeaturePlot(integrated_obj, features = g, reduction = 'umap', order = T))
        }
    }
dev.off()


########################################################################################################################
########################################################################################################################
########################################################################################################################
optimal_clustering_dir <- paste0(out_dir, '/', sample_id, '/', 'optimal_cluster')
dir.create(optimal_clustering_dir, recursive = T)


opt_res <- 1.1
cluster_name <- grep(paste0('res.', opt_res), colnames(integrated_obj@meta.data), value = T)[1]


# make a plot with label at the optimal cluster resolution ----
pdf(file = paste0(optimal_clustering_dir, '/', 'umap_', opt_res, '.pdf'))
    print(
        DimPlot(integrated_obj, reduction = "umap", group.by = cluster_name, label = T, repel = T)
    )
dev.off()


#######################################################################################################################
#######################################################################################################################
#######################################################################################################################
subcluster_dir <- paste0(optimal_clustering_dir, '/', 'subclusters')
dir.create(subcluster_dir, recursive = T)


Idents(integrated_obj) <- cluster_name


# subcluster a few clusters ----
integrated_obj <- FindSubCluster(
    integrated_obj,
    cluster = '15',
    graph.name = 'integrated_snn',
    subcluster.name = "C15_subcluster",
    resolution = 0.5,
    algorithm = 1)

pdf(file = paste0(subcluster_dir, '/', 'C15_subclusters.pdf'), width = 20, 
    height = ceiling(length(unique(integrated_obj$C15_subcluster))/5)*4)
    print(
        DimPlot(integrated_obj, group.by = 'C15_subcluster', split.by = 'C15_subcluster', ncol = 5)
        
    )
    print(DimPlot(integrated_obj, group.by = 'C15_subcluster'))
dev.off()


integrated_obj <- FindSubCluster(
    integrated_obj,
    cluster = '22',
    graph.name = 'integrated_snn',
    subcluster.name = "C22_subcluster",
    resolution = 0.5,
    algorithm = 1)

pdf(file = paste0(subcluster_dir, '/', 'C22_subcluster.pdf'), width = 20, 
    height = ceiling(length(unique(integrated_obj$C22_subcluster))/5)*4)
    print(
        DimPlot(integrated_obj, group.by = 'C22_subcluster', split.by = 'C22_subcluster', ncol = 5)
        
    )
    print(DimPlot(integrated_obj, group.by = 'C22_subcluster'))
dev.off()


integrated_obj <- FindSubCluster(
    integrated_obj,
    cluster = '18',
    graph.name = 'integrated_snn',
    subcluster.name = "C18_subcluster",
    resolution = 0.5,
    algorithm = 1)

pdf(file = paste0(subcluster_dir, '/', 'C18_subcluster.pdf'), width = 20, 
    height = ceiling(length(unique(integrated_obj$C18_subcluster))/5)*4)
    print(
        DimPlot(integrated_obj, group.by = 'C18_subcluster', split.by = 'C18_subcluster', ncol = 5)
        
    )
    print(DimPlot(integrated_obj, group.by = 'C18_subcluster'))
dev.off()


# modified annotation
integrated_obj$modified_cluster_1.1 <- 
    ifelse(as.character(integrated_obj@meta.data[[cluster_name]]) == '22', as.character(integrated_obj@meta.data[['C22_subcluster']]),
    ifelse(as.character(integrated_obj@meta.data[[cluster_name]]) == '18', as.character(integrated_obj@meta.data[['C18_subcluster']]),
    ifelse(as.character(integrated_obj@meta.data[[cluster_name]]) == '15', as.character(integrated_obj@meta.data[['C15_subcluster']]),
    as.character(integrated_obj@meta.data[[cluster_name]]))))


pdf(file = paste0(optimal_clustering_dir, '/', 'umap_', opt_res, '_subclustered.pdf'))
    print(
        DimPlot(integrated_obj, reduction = "umap", group.by = 'modified_cluster_1.1', label = T, repel = T)
    )
dev.off()


####################################################################################
####################################################################################
####################################################################################
# annotate clusters ----
Idents(integrated_obj) <- 'modified_cluster_1.1'

new_annotation_df <- data.frame(cluster_id = unique(as.character(Idents(integrated_obj))),
                                initial_annotation = '')
new_annotation_df[new_annotation_df$cluster_id == '0', 'initial_annotation'] <- 'Epithelial_cancer'
new_annotation_df[new_annotation_df$cluster_id == '1', 'initial_annotation'] <- 'Epithelial_cancer'
new_annotation_df[new_annotation_df$cluster_id == '2', 'initial_annotation'] <- 'Epithelial_cancer'
new_annotation_df[new_annotation_df$cluster_id == '3', 'initial_annotation'] <- 'Epithelial_cancer'
new_annotation_df[new_annotation_df$cluster_id == '4', 'initial_annotation'] <- 'Epithelial_cancer'
new_annotation_df[new_annotation_df$cluster_id == '5', 'initial_annotation'] <- 'Epithelial_cancer'
new_annotation_df[new_annotation_df$cluster_id == '6', 'initial_annotation'] <- 'Epithelial_cancer'
new_annotation_df[new_annotation_df$cluster_id == '7', 'initial_annotation'] <- 'CAF'
new_annotation_df[new_annotation_df$cluster_id == '8', 'initial_annotation'] <- 'Endothelial'
new_annotation_df[new_annotation_df$cluster_id == '9', 'initial_annotation'] <- 'Epithelial_cancer'
new_annotation_df[new_annotation_df$cluster_id == '10', 'initial_annotation'] <- 'Epithelial_cancer'
new_annotation_df[new_annotation_df$cluster_id == '11', 'initial_annotation'] <- 'Epithelial_cancer'
new_annotation_df[new_annotation_df$cluster_id == '12', 'initial_annotation'] <- 'Endothelial'
new_annotation_df[new_annotation_df$cluster_id == '13', 'initial_annotation'] <- 'PVL'
new_annotation_df[new_annotation_df$cluster_id == '14', 'initial_annotation'] <- 'Epithelial_cancer'
new_annotation_df[new_annotation_df$cluster_id == '15_0', 'initial_annotation'] <- 'Hepatocyte'
new_annotation_df[new_annotation_df$cluster_id == '15_1', 'initial_annotation'] <- 'Hepatocyte'
new_annotation_df[new_annotation_df$cluster_id == '15_2', 'initial_annotation'] <- 'Hepatocyte'
new_annotation_df[new_annotation_df$cluster_id == '15_3', 'initial_annotation'] <- 'Hepatocyte'
new_annotation_df[new_annotation_df$cluster_id == '15_4', 'initial_annotation'] <- 'Cholangiocyte'
new_annotation_df[new_annotation_df$cluster_id == '16', 'initial_annotation'] <- 'CAF'
new_annotation_df[new_annotation_df$cluster_id == '17', 'initial_annotation'] <- 'Macrophage'
new_annotation_df[new_annotation_df$cluster_id == '18_0', 'initial_annotation'] <- 'Epithelial_cancer'
new_annotation_df[new_annotation_df$cluster_id == '18_1', 'initial_annotation'] <- 'Epithelial_cancer'
new_annotation_df[new_annotation_df$cluster_id == '18_2', 'initial_annotation'] <- 'Epithelial_cancer'
new_annotation_df[new_annotation_df$cluster_id == '18_3', 'initial_annotation'] <- 'Epithelial_cancer'
new_annotation_df[new_annotation_df$cluster_id == '18_4', 'initial_annotation'] <- 'RBCs'
new_annotation_df[new_annotation_df$cluster_id == '19', 'initial_annotation'] <- 'PVL'
new_annotation_df[new_annotation_df$cluster_id == '20', 'initial_annotation'] <- 'Epithelial_cancer'
new_annotation_df[new_annotation_df$cluster_id == '21', 'initial_annotation'] <- 'Epithelial_cancer'
new_annotation_df[new_annotation_df$cluster_id == '22_0', 'initial_annotation'] <- 'LSEC'
new_annotation_df[new_annotation_df$cluster_id == '22_1', 'initial_annotation'] <- 'Lymphatic_endothelial'
new_annotation_df[new_annotation_df$cluster_id == '23', 'initial_annotation'] <- "Mixed_lymphocytes"


# annotate the object
new_annotations <- new_annotation_df$initial_annotation
names(new_annotations) <- new_annotation_df$cluster_id

integrated_obj <- RenameIdents(integrated_obj, new_annotations)
integrated_obj$initial_annotation <- Idents(integrated_obj)


# export cell type annotations as csv file for easier modification and assessemnt
write.csv(file = paste0(optimal_clustering_dir, '/', 'initial_annotation.csv'), integrated_obj$initial_annotation)


# plot a umap with new annotations
Idents(integrated_obj) <- 'initial_annotation'
pdf(file = paste0(optimal_clustering_dir, '/', 'umap_initial_annotation.pdf'), width = 10, height = 7)
    print(
        DimPlot(integrated_obj, reduction = "umap", group.by = 'initial_annotation', label = T, repel = T)
    )
dev.off()


integrated_obj$major_annotation <- integrated_obj$initial_annotation


####################################################################################
####################################################################################
####################################################################################
# save annotated data
saveRDS(file = '/share/ScratchGeneral/tonwan/projects/snPATHO-Seq/data/snPATHO_seq/25k_downsampled_processed_objs/integrated/4399_cellbender_filtered_integration_final_annotation.rds',
    integrated_obj)
```
