## Supplementary material for "snPATHO-seq: unlocking the pathology archives": scripts: 02_cell_type_annotation_1.1.R

```
# aim ----
    # annotate cell types


# define environment ----
library(Seurat)
library(tidyverse)
library(RColorBrewer)


# arguments ----
data_dir <- '/share/ScratchGeneral/tonwan/projects/snPATHO-Seq/data/snPATHO_seq/25k_downsampled_processed_objs/integrated'
obj_file <- '4411_cellbender_filtered_integration.rds'

sample_id <- '4411'

out_dir <- '/share/ScratchGeneral/tonwan/projects/snPATHO-Seq/results/integration/25k_downsampled/cellbender'


# load data ----
integrated_obj <- readRDS(file = paste0(data_dir, '/', obj_file))


# check the expression of selected cell type markers 
marker_dir <- paste0(out_dir, '/', sample_id, '/', 'markers')
dir.create(marker_dir, recursive = T)

markers <- c('PROX1', 'HGF', 'HPX', 'ALB', 'CLEC4G', 'APOE',
    'SPTA1', 'HBB', 'SPTB',
    'ABCC1', 'ALDH1A2', 'HNF1B', 'SOX9', 'ONECUT1',
    'ASGR1')

DefaultAssay(integrated_obj) <- 'RNA'
to_plot <- markers[markers %in% rownames(integrated_obj)]
for (g in to_plot) {
    pdf(paste0(marker_dir, '/', g, '.pdf'))
        print(FeaturePlot(integrated_obj, features = g, reduction = 'umap', order = T))
    dev.off()
}


top_periportal_hep_genes <- c('ALB', 'GC', 'APOH', 'FGG', 'SERPINC1', 'BCHE', 'FGB',
    'CAT', 'CFHR1', 'UGT2B4')

pdf(paste0(marker_dir, '/', deparse(substitute(top_periportal_hep_genes)), '.pdf'))
    for (g in top_periportal_hep_genes) {
        if (g %in% c(rownames(integrated_obj))) {
            print(FeaturePlot(integrated_obj, features = g, reduction = 'umap', order = T))
        }
    }
dev.off()


top_central_hep_genes <- c('PPARA', 'MAT1A', 'RPL36', 'MT1H', 'RPLP1', 'SLTM',
    'PCK2', 'RPLP2', 'GLTSCR2', 'HIPK2')

pdf(paste0(marker_dir, '/', deparse(substitute(top_central_hep_genes)), '.pdf'))
    for (g in top_central_hep_genes) {
        if (g %in% c(rownames(integrated_obj))) {
            print(FeaturePlot(integrated_obj, features = g, reduction = 'umap', order = T))
        }
    }
dev.off()


Cholangiocyte <- c('ANXA4', 'KRT7', 'DEFB1', 'FXYD2', 'TM4SF4', 'KRT19', 'KRT8', 'TACSTD2', 'EGRI1', 'SPP1')

pdf(paste0(marker_dir, '/', deparse(substitute(Cholangiocyte)), '.pdf'))
    for (g in Cholangiocyte) {
        if (g %in% c(rownames(integrated_obj))) {
            print(FeaturePlot(integrated_obj, features = g, reduction = 'umap', order = T))
        }
    }
dev.off()


astrocyte <- c('SLC1A2', 'ADGRV1', 'SLC1A3', 'GPC5', 'RNF219.AS1', 'ARHGAP24', 'CST3', 'HPSE2', 'AQP4', 'COL5A3')

pdf(paste0(marker_dir, '/', deparse(substitute(astrocyte)), '.pdf'))
    for (g in astrocyte) {
        if (g %in% c(rownames(integrated_obj))) {
            print(FeaturePlot(integrated_obj, features = g, reduction = 'umap', order = T))
        }
    }
dev.off()


########################################################################################################################
########################################################################################################################
########################################################################################################################
optimal_clustering_dir <- paste0(out_dir, '/', sample_id, '/', 'optimal_cluster')
dir.create(optimal_clustering_dir, recursive = T)


opt_res <- 1.1
cluster_name <- grep(paste0('res.', opt_res), colnames(integrated_obj@meta.data), value = T)[1]


# make a plot with label at the optimal cluster resolution ----
pdf(file = paste0(optimal_clustering_dir, '/', 'umap_', opt_res, '.pdf'))
    print(
        DimPlot(integrated_obj, reduction = "umap", group.by = cluster_name, label = T, repel = T)
    )
dev.off()


# check markers for selected clusters 
de_dir <- paste0(optimal_clustering_dir, '/', 'de_markers')
dir.create(de_dir, recursive = T)

Idents(integrated_obj) <- cluster_name

for (cl in c(22, 25)) {
    de_markrers <- FindMarkers(integrated_obj, ident.1 = cl, test.use = 't', only.pos = TRUE)
    write.csv(file = paste0(de_dir, '/', 'Cluster_', cl, '_markers.csv'), de_markrers)
}


####################################################################################
####################################################################################
####################################################################################
# annotate clusters ----
Idents(integrated_obj) <- cluster_name

new_annotation_df <- data.frame(cluster_id = unique(as.character(Idents(integrated_obj))),
                                initial_annotation = '')
new_annotation_df[new_annotation_df$cluster_id == '0', 'initial_annotation'] <- 'Epithelial_cancer'
new_annotation_df[new_annotation_df$cluster_id == '1', 'initial_annotation'] <- 'Epithelial_cancer'
new_annotation_df[new_annotation_df$cluster_id == '2', 'initial_annotation'] <- 'Epithelial_cancer'
new_annotation_df[new_annotation_df$cluster_id == '3', 'initial_annotation'] <- 'Hepatocyte'
new_annotation_df[new_annotation_df$cluster_id == '4', 'initial_annotation'] <- 'Epithelial_cancer'
new_annotation_df[new_annotation_df$cluster_id == '5', 'initial_annotation'] <- 'Hepatocyte'
new_annotation_df[new_annotation_df$cluster_id == '6', 'initial_annotation'] <- 'Hepatocyte'
new_annotation_df[new_annotation_df$cluster_id == '7', 'initial_annotation'] <- 'Epithelial_cancer'
new_annotation_df[new_annotation_df$cluster_id == '8', 'initial_annotation'] <- 'Epithelial_cancer'
new_annotation_df[new_annotation_df$cluster_id == '9', 'initial_annotation'] <- 'Hepatocyte'
new_annotation_df[new_annotation_df$cluster_id == '10', 'initial_annotation'] <- 'Epithelial_cancer'
new_annotation_df[new_annotation_df$cluster_id == '11', 'initial_annotation'] <- 'Endothelial'
new_annotation_df[new_annotation_df$cluster_id == '12', 'initial_annotation'] <- 'Epithelial_cancer'
new_annotation_df[new_annotation_df$cluster_id == '13', 'initial_annotation'] <- 'Epithelial_cancer'
new_annotation_df[new_annotation_df$cluster_id == '14', 'initial_annotation'] <- 'Macrophage'
new_annotation_df[new_annotation_df$cluster_id == '15', 'initial_annotation'] <- 'LSEC'
new_annotation_df[new_annotation_df$cluster_id == '16', 'initial_annotation'] <- 'Epithelial_cancer'
new_annotation_df[new_annotation_df$cluster_id == '17', 'initial_annotation'] <- 'Epithelial_cancer'
new_annotation_df[new_annotation_df$cluster_id == '18', 'initial_annotation'] <- 'CAF'
new_annotation_df[new_annotation_df$cluster_id == '19', 'initial_annotation'] <- 'RBCs'
new_annotation_df[new_annotation_df$cluster_id == '20', 'initial_annotation'] <- 'Hepatocyte'
new_annotation_df[new_annotation_df$cluster_id == '21', 'initial_annotation'] <- 'Endothelial_proliferative'
new_annotation_df[new_annotation_df$cluster_id == '22', 'initial_annotation'] <- 'Epithelial_cancer'
new_annotation_df[new_annotation_df$cluster_id == '23', 'initial_annotation'] <- 'Mixed_lymphocytes'
new_annotation_df[new_annotation_df$cluster_id == '24', 'initial_annotation'] <- 'Cholangiocyte'
new_annotation_df[new_annotation_df$cluster_id == '25', 'initial_annotation'] <- 'Epithelial_cancer'


# annotate the object
new_annotations <- new_annotation_df$initial_annotation
names(new_annotations) <- new_annotation_df$cluster_id

integrated_obj <- RenameIdents(integrated_obj, new_annotations)
integrated_obj$initial_annotation <- Idents(integrated_obj)


# export cell type annotations as csv file for easier modification and assessemnt
write.csv(file = paste0(optimal_clustering_dir, '/', 'initial_annotation.csv'), integrated_obj$initial_annotation)


# plot a umap with new annotations
Idents(integrated_obj) <- 'initial_annotation'
pdf(file = paste0(optimal_clustering_dir, '/', 'umap_initial_annotation.pdf'), width = 10, height = 7)
    print(
        DimPlot(integrated_obj, reduction = "umap", group.by = 'initial_annotation', label = T, repel = T)
    )
dev.off()


####################################################################################
####################################################################################
####################################################################################
# save annotated data
saveRDS(file = paste0(data_dir, '/', sample_id, '_cellbender_filtered_integration_initial_annotation.rds'), integrated_obj)
```
