## Supplementary material for "snPATHO-seq: unlocking the pathology archives": scripts: summarise_CNV.R

```
# aim ----
    # gather all inferCNV HMM predictions
    # plot all CNV results together


# define environment ----
library(Seurat)
library(tidyverse)

library(ggrepel)
library(cowplot) 


# Common arguments ----
gene_order_file <- '/share/ScratchGeneral/tonwan/projects/snPATHO-Seq/resources/Javier_gene_ordering_hg38_gencode_v27.txt'

out_dir <- '/share/ScratchGeneral/tonwan/projects/snPATHO-Seq/results/integration/25k_downsampled/cellbender/CNV_summary'


# load gene order file ----
gene_order <- read.delim(gene_order_file, header = FALSE)
colnames(gene_order) <- c('Gene_name', 'Chromosome', 'start', 'end')
gene_order$gene_positions <- rownames(gene_order)
gene_order$gene_positions <- as.integer(gene_order$gene_positions)
rownames(gene_order) <- gene_order$Gene_name


# load all HMM results from all samples ----
# 4066
CNV_HMM_res_path <- '/share/ScratchGeneral/tonwan/projects/snPATHO-Seq/results/integration/25k_downsampled/cellbender/4066/cnv_HMM_res'
samples <- dir(CNV_HMM_res_path)

snRNA_obj_file <- '/share/ScratchGeneral/tonwan/projects/snPATHO-Seq/data/snPATHO_seq/25k_downsampled_processed_objs/integrated/4066_cellbender_filtered_integration_final_annotation.rds'
obj <- readRDS(file = snRNA_obj_file)

res_list <- list()
for (s in samples) {
    HMM_res <- read.delim(paste0(CNV_HMM_res_path, '/', s, '/', 'infercnv.20_HMM_predHMMi6.hmm_mode-samples.Pnorm_0.5.repr_intensities.observations.txt'),
        sep = ' ', header = TRUE)

    colnames(HMM_res) <- gsub('\\.', '\\-', colnames(HMM_res))
    res_list[[s]] <- HMM_res
}

# remove barcodes containing normal epithelials 
barcodes_to_remove <- rownames(obj@meta.data[obj$major_annotation %in% c('Epithelial_basal', 'Epithelial_luminal', 'Myoepithelial'), ])

filtered_res_list_4066 <- lapply(res_list, function(x){
    x <- x[, !(colnames(x) %in% barcodes_to_remove)]
})


# 4399
CNV_HMM_res_path <- '/share/ScratchGeneral/tonwan/projects/snPATHO-Seq/results/integration/25k_downsampled/cellbender/4399/cnv_HMM_res'
samples <- dir(CNV_HMM_res_path)

snRNA_obj_file <- '/share/ScratchGeneral/tonwan/projects/snPATHO-Seq/data/snPATHO_seq/25k_downsampled_processed_objs/integrated/4399_cellbender_filtered_integration_final_annotation.rds'
obj <- readRDS(file = snRNA_obj_file)

res_list <- list()
for (s in samples) {
    HMM_res <- read.delim(paste0(CNV_HMM_res_path, '/', s, '/', 'infercnv.20_HMM_predHMMi6.hmm_mode-samples.Pnorm_0.5.repr_intensities.observations.txt'),
        sep = ' ', header = TRUE)

    colnames(HMM_res) <- gsub('\\.', '\\-', colnames(HMM_res))
    res_list[[s]] <- HMM_res
}

# remove barcodes containing normal epithelials 
barcodes_to_remove <- rownames(obj@meta.data[obj$major_annotation %in% c('Cholangiocyte'), ])

filtered_res_list_4399 <- lapply(res_list, function(x){
    x <- x[, !(colnames(x) %in% barcodes_to_remove)]
})


# 4411
CNV_HMM_res_path <- '/share/ScratchGeneral/tonwan/projects/snPATHO-Seq/results/integration/25k_downsampled/cellbender/4411/cnv_HMM_res'
samples <- dir(CNV_HMM_res_path)

snRNA_obj_file <- '/share/ScratchGeneral/tonwan/projects/snPATHO-Seq/data/snPATHO_seq/25k_downsampled_processed_objs/integrated/4411_cellbender_filtered_integration_final_annotation.rds'
obj <- readRDS(file = snRNA_obj_file)

res_list <- list()
for (s in samples) {
    HMM_res <- read.delim(paste0(CNV_HMM_res_path, '/', s, '/', 'infercnv.20_HMM_predHMMi6.hmm_mode-samples.Pnorm_0.5.repr_intensities.observations.txt'),
        sep = ' ', header = TRUE)

    colnames(HMM_res) <- gsub('\\.', '\\-', colnames(HMM_res))
    res_list[[s]] <- HMM_res
}

# remove barcodes containing normal epithelials 
barcodes_to_remove <- rownames(obj@meta.data[obj$major_annotation %in% c('Cholangiocyte'), ])

filtered_res_list_4411 <- lapply(res_list, function(x){
    x <- x[, !(colnames(x) %in% barcodes_to_remove)]
})


#######################################################################################################################
#######################################################################################################################
#######################################################################################################################
# summarise data
# concatenate all lists ----
merged_list <- c(filtered_res_list_4066, filtered_res_list_4399, filtered_res_list_4411)


# summarise all CNV results together for plotting ----
summarised_CNV_profiles <- lapply(seq_along(merged_list), function(x){
    dataset_name <- names(merged_list)[[x]]
    working_cnv_res <- merged_list[[x]]

    # get mean CNV prediction per gene across all cells within each cluster
    per_gene_average <- apply(working_cnv_res, 1, mean)
    # round up to 0.5
    per_gene_average <- plyr::round_any(per_gene_average, 0.5)
    # turn results into a dataframe and merge with the gene order file
    summarised_cnv_res <- as.data.frame(per_gene_average)
    colnames(summarised_cnv_res) <- 'CNV_HMM_prediction'
    summarised_cnv_res <- base::merge(gene_order, summarised_cnv_res, by = 0, all = TRUE)
    # make a label for genes without any prediction
    summarised_cnv_res$detection_status <- ifelse(is.na(summarised_cnv_res$CNV_HMM_prediction), 'no_prediction_made', 'predicted')
    # replace genes without any prediction with 1
    summarised_cnv_res$CNV_HMM_prediction[is.na(summarised_cnv_res$CNV_HMM_prediction)] <- 1
    summarised_cnv_res <- column_to_rownames(summarised_cnv_res, var = 'Row.names')
    # order the results by position
    summarised_cnv_res <- summarised_cnv_res %>% 
        arrange(gene_positions) %>% 
        as.data.frame()
    summarised_cnv_res$dataset_label <- dataset_name

    return(summarised_cnv_res)
})

# turn processing results into a dataframe for plotting
merged_res_df <- Reduce(rbind, summarised_CNV_profiles)


saveRDS(file = paste0(out_dir, '/', 'summarised_CNV_res_dataframe.rds'), merged_res_df)
write.csv(file = paste0(out_dir, '/', 'summarised_CNV_res_dataframe.csv'), merged_res_df)


#######################################################################################################################
#######################################################################################################################
#######################################################################################################################
# plot line/path plot for predicted CNV profiles 

# format metadata
merged_res_df$sample <- 
    ifelse(merged_res_df$dataset_label %in% c('4066FFPE', '4066GEX', '4066SNAPFix'), '4066',
    ifelse(merged_res_df$dataset_label %in% c('4399FFPE_run1', '4399FFPE_run2', '4399GEX', '4399SNAPFix'), '4399',
    ifelse(merged_res_df$dataset_label %in% c('4411FFPE', '4411GEX', '4411SNAPFix'), '4411', 'NA')))

merged_res_df$processing_method <- 
    ifelse(merged_res_df$dataset_label %in% c('4066FFPE', '4411FFPE', '4399FFPE_run1'), 'snPATHO',
    ifelse(merged_res_df$dataset_label %in% c('4066GEX', '4399GEX', '4411GEX'), '3p',
    ifelse(merged_res_df$dataset_label %in% c('4066SNAPFix', '4399SNAPFix', '4411SNAPFix'), 'FLEX', 'NA')))


# define line colors based on CNV gain or loss ----
merged_res_df$CNV_colors <- 
    ifelse(merged_res_df$CNV_HMM_prediction < 1, 'blue',
    ifelse(merged_res_df$CNV_HMM_prediction > 1, 'red', 'white'))
# don't plot regions without CNV change
merged_res_df$CNV_alpha <- 
    ifelse(merged_res_df$CNV_HMM_prediction < 1, 1,
    ifelse(merged_res_df$CNV_HMM_prediction > 1, 1, 0))

# mock a dataframe and ggplot for the legend
CNV_color_legend_plot <- ggplot(
    data.frame(x = c(1, 2), y = c(1, 2), CNV_status = c('Gain', 'Loss')),
    aes(x = x, y = y, fill = CNV_status)) +
        geom_col() +
        scale_fill_manual(values = c('red', 'blue')) +
        labs(fill = 'CNV status')

color_legend <- get_legend(CNV_color_legend_plot)


# define chrosome separation ----
separations <- gene_order %>% 
    group_by(Chromosome) %>% 
    slice_head(n = 1) %>%
    arrange(gene_positions, decreasing = TRUE) %>% 
    pull(gene_positions)

# only keep the first 22 chrosomes as they were in the data
separations <- separations[1:22]
chrosome_labels <- paste0('Chr', seq(1:22))
chrosome_df <- data.frame(x = separations, y = 0, chrosome_labels = chrosome_labels,
    sample = '4411', processing_method = 'snPATHO')


# plot a few known drivers that might have been affected by CNV ----
drivers_to_plot <- c('ERBB2', 'MDM4',  "MYC", 'AURKA', 'VEGFA', 'RB1', 'KAT6A', 'AKT2', 'MAPK1')

selected_driver_label_df <- data.frame(
    x = merged_res_df[drivers_to_plot, 'gene_positions'],
    y = 3,
    gene_name = drivers_to_plot, 
    sample = '4066',
    processing_method = '3p')


# plot line plot representing CNV profiles ----
plot <- ggplot(merged_res_df, aes(x = gene_positions, y = CNV_HMM_prediction)) +
            geom_path(aes(colour = .data[['CNV_colors']], alpha = .data[['CNV_alpha']]), group = 1) +
            scale_colour_identity() +
            geom_vline(xintercept = separations, linetype = 'dashed', alpha = 0.4) +
            geom_hline(yintercept = 1, linewidth = 0.8) +
            geom_text_repel(data = chrosome_df, aes(x = x, y = y, label = chrosome_labels), 
                size = 3, min.segment.length = 0, direction = 'x', 
                segment.curvature = 0.2, segment.ncp = 3, segment.angle = 30) +
            geom_text_repel(data = selected_driver_label_df, aes(x = x, y = y, label = gene_name), 
                size = 3, min.segment.length = 0, direction = 'x', nudge_y = 1,
                segment.curvature = 0.2, segment.ncp = 3, segment.angle = 30) +
            facet_grid(rows = vars(sample, processing_method)) +
            theme_bw() +
            theme(axis.text.x = element_blank(),
                axis.ticks = element_blank(),
                legend.position = "none") +
            labs(x = 'Genomic Regions',
                y = 'InferCNV HMM prediction results')
            

# ggsave(file = paste0(out_dir, '/', 'test.pdf'), plot, device = 'pdf', width = 14, height = 14)


# change facet colors ----
# change facet color depending on the snRNA-seq workflow

# load predefined colors
source('/share/ScratchGeneral/tonwan/projects/snPATHO-Seq/scripts/colors.R')

# define the color vectors to change
plot_order_df <- data.frame(sample = merged_res_df$sample, processing_method = merged_res_df$processing_method)
plot_order <- plot_order_df %>% 
    distinct() %>% 
    arrange(sample, processing_method) %>% 
    pull(processing_method)
# match colors
snRNA_workflow_colors <- workflow_colors[plot_order]


# get build info of the plot
g <- ggplot_gtable(ggplot_build(plot))
# get index of the facet labels
stripr <- which(grepl('strip-r', g$layout$name))


# for this analysis:
    # facet layer will be in reverse order as the input above
        # e.g. in ggplot above, facet goes as sample, then workflow
        # in the plot, it goes workflow then sample
for (i in seq_along(stripr)) {
    # change background of the first facet factor - workflow in this case
    g$grobs[[stripr[[i]]]]$grobs[[1]]$children[[1]]$gp$fill <- snRNA_workflow_colors[[i]]
    # change font color to white to look better with colored background
    g$grobs[[stripr[[i]]]]$grobs[[1]]$children[[2]]$children[[1]]$gp$col <- 'white'
    
    # change background of the 2nd facet facor - sample in this case
    g$grobs[[stripr[[i]]]]$grobs[[2]]$children[[1]]$gp$fill <- 'white'
    
}


# put main plot and color legend together ----
plot_with_legend <- ggdraw() +
    draw_plot(g, x = 0, y = 0, width = .9, height = 1) +
    draw_plot(color_legend, x = .9, y = 0, width = .1, height = 1) +
    theme(text = element_text(size = 12))

ggsave(file = paste0(out_dir, '/', 'CNV_summary_lineplot.pdf'), plot_with_legend, device = 'pdf', width = 14, height = 12)
```
