## Supplementary material for "snPATHO-seq: unlocking the pathology archives": scripts: 4066.R

```
# aim ----
    # basic filtering and processing of Visium data


# define environment ----
library(STutility)
library(Seurat)
library(tidyverse)
library(RColorBrewer)


# arguments ----
data_dir <- '/share/ScratchGeneral/tonwan/projects/snPATHO-Seq/data/Visium/sr_counts/V42Y04-403_A1'

out_dir <- '/share/ScratchGeneral/tonwan/projects/snPATHO-Seq/results/visium_individual_sample/V42Y04-403_A1'
dir.create(out_dir, recursive = T)

out_data_dir <- '/share/ScratchGeneral/tonwan/projects/snPATHO-Seq/data/Visium/processed/STutility'

path_annotation_file <- '/share/ScratchGeneral/tonwan/projects/snPATHO-Seq/data/Visium/spot_annotation/4066_pathology_annotation.csv'


# load spaceranger data ----
samples <- file.path(data_dir, "outs", "filtered_feature_bc_matrix.h5")
imgs <- file.path(data_dir, 'outs', "spatial", "tissue_hires_image.png")
spotfiles <- file.path(data_dir, 'outs', "spatial", "tissue_positions_list.csv")
json <- file.path(data_dir, 'outs', "spatial", "scalefactors_json.json")

infoTable <- data.frame(samples, imgs, spotfiles, json) # Add additional column

obj <- InputFromTable(infoTable)
obj <- LoadImages(obj, time.resolve = FALSE)
obj <- RenameAssays(object = obj, RNA = 'Spatial')


# lock in image ratio ----
image <- GetStaffli(obj)
x_length <- image@dims[[1]]$width
y_length <- image@dims[[1]]$height

xdim <- (x_length/2000) * 10
ydim <- (y_length/2000) * 9


########################################################################################################################
########################################################################################################################
########################################################################################################################
# remove spots on the edge
    # probes often accumulte on the border of the Visium cassette causing artefacts
spots_to_remove <- rownames(image@meta.data[image@meta.data$x %in% c(0, 127) | image@meta.data$y %in% c(0, 77), , drop=F])

obj$spots_to_remove <- ifelse(rownames(obj@meta.data) %in% spots_to_remove, 'to_remove', 'keep')
obj <- SubsetSTData(obj, expression = spots_to_remove == 'keep')


# plot to check if filtering is enough
plot <- ST.FeaturePlot(obj, features = 'nFeature_Spatial',
                        pt.size = 2, pt.border = F, show.sb = F,
                        cols = c("lightgray", "mistyrose", "red", "darkred", "black")) +
                        theme(aspect.ratio = y_length / x_length)
ggsave(file = paste0(out_dir, '/', 'spatial_gene_distribution.pdf'),
       device = 'pdf',
       plot,
       width = xdim,
       height = ydim)


########################################################################################################################
########################################################################################################################
########################################################################################################################
# crop image ----
    # a whole slide scan was supplied but only a small region was targeted for Visium

# get crop window size with a bit of extra buffer - 100 pixels
crop_width <- max(obj@tools$Staffli@meta.data$pixel_x) - min(obj@tools$Staffli@meta.data$pixel_x) + 100
crop_height <- max(obj@tools$Staffli@meta.data$pixel_y) - min(obj@tools$Staffli@meta.data$pixel_y) + 100

# find centroid of the data
centroid_x <- mean(range(obj@tools$Staffli@meta.data$pixel_x))
centroid_y <- mean(range(obj@tools$Staffli@meta.data$pixel_y))

# calculate the offset
x_offset <- centroid_x - (crop_width / 2)
y_offset <- centroid_y - (crop_height / 2)

# define the crop window
crop_windows <- list("1" = paste0(crop_width, 'x', crop_height, '+', x_offset, '+', y_offset))

# crop the image
obj_cropped <- CropImages(obj, crop.geometry.list = crop_windows, time.resolve = TRUE, verbose = TRUE)


# plot H&E after cropping
pdf(file = paste0(out_dir, '/H&E_cropped.pdf'),
    width = 7, height = 7)
    print(ImagePlot(obj_cropped, 
                type = 'raw',
                method = "raster", 
                annotate = FALSE,
                fix.axes = T))
dev.off()


# plot pathological annotation
path_annotation <- read.csv(file = path_annotation_file, row.names = 1)
rownames(path_annotation) <- paste0(rownames(path_annotation), '_1')

path_annotation <- path_annotation[rownames(obj_cropped@meta.data), , drop= FALSE]

obj_cropped$path_annotation <- path_annotation[[1]]


# get coordinates of the visium spots
image <- GetStaffli(obj_cropped)
image_subset <- image@meta.data[colnames(obj_cropped), ]
x_dim <- image_subset$x 
y_dim <- image_subset$y


plot_df <- data.frame(x = x_dim,
                        y = y_dim,
                        value = obj_cropped$path_annotation)
plot_df$value <- ifelse(plot_df$value == 'exclude', 'necrosis', plot_df$value)

plot <- ggplot(plot_df, aes(x = x, y = y, color = value, fill = value)) +
            geom_point(shape = 21) +
            scale_fill_brewer(palette = 'Paired') +
            scale_color_brewer(palette = 'Paired') +
            theme_bw() +
            theme(panel.grid = element_blank(),
                axis.text = element_blank(),
                axis.ticks = element_blank(),
                panel.border = element_blank(),
                aspect.ratio = (max(y_dim)*1.7) / max(x_dim)) +
            labs(title = NULL,
                fill = 'Regional annotation',
                x = NULL,
                y = NULL) +
            guides(color = 'none')
ggsave(file = paste0(out_dir, '/', 'pathology_annotation', '.pdf'),
        device = 'pdf', plot, width = 8, height = 7)


# plot a few markers 
marker_dir <- paste0(out_dir, '/', 'markers')
dir.create(marker_dir, recursive = T)

markers <- c('BMP2', 'PTHLH',
    'ACACB','ADGRV1','ANKRD30B','CADPS2','CAPN13','CLPSL1','CPED1','CYP21A2','DNAH14','DPY19L3','EIF4G3','GCNT2','GLI3','HOXC9','INTS6','ITPR2','KRTAP2-4','LURAP1L','MAST4','MGAT4A','MPV17','MYEOV','NAV1','NFATC4','OGT','PCNX2','PLA2R1','PLK2','PRKG1','RHOBTB3','SLC4A7','SLC4A8','TBC1D30','TTC6','ZNF512B','ZNF580',
    'PTCH1')

for (m in markers) {
    plot_df <- data.frame(x = x_dim,
                        y = y_dim,
                        value = as.matrix(GetAssayData(obj_cropped, assay = 'Spatial', slot = 'data'))[m, ])
    
    plot <- ggplot(plot_df, aes(x = x, y = y, color = value, fill = value)) +
            geom_point(shape = 21) +
            scale_fill_gradient(low = '#e5e5e5', high = '#B2182B') +
            scale_color_gradient(low = '#e5e5e5', high = '#B2182B') +
            theme_bw() +
            theme(panel.grid = element_blank(),
                axis.text = element_blank(),
                axis.ticks = element_blank(),
                panel.border = element_blank(),
                aspect.ratio = (max(y_dim)*1.7) / max(x_dim)) +
            labs(title = NULL,
                fill = 'Expression',
                x = NULL,
                y = NULL) +
            guides(color = 'none')
    ggsave(file = paste0(marker_dir, '/', m, '.pdf'),
        device = 'pdf', plot, width = 8, height = 7)

}


########################################################################################################################
########################################################################################################################
########################################################################################################################
# save object ----
saveRDS(file = paste0(out_data_dir, '/', '4066FFPE.rds'), obj_cropped)
```
