## Supplementary material for "snPATHO-seq: unlocking the pathology archives": scripts: run_RCTD.R

```
# aim ---
    # run RCTD deconvolution


# define environment ----
library(STutility)
library(Seurat)
library(tidyverse)
library(RColorBrewer)

library(spacexr)
library(Matrix)


# arguments ----
# ref_obj_file <- '/share/ScratchGeneral/tonwan/projects/snPATHO-Seq/data/snPATHO_seq/25k_downsampled_processed_objs/integrated/4411_cellbender_filtered_integration_final_annotation.rds'
# ref_obj_file <- '/share/ScratchGeneral/tonwan/projects/snPATHO-Seq/data/snPATHO_seq/25k_downsampled_processed_objs/integrated/4399_cellbender_filtered_integration_final_annotation.rds'
ref_obj_file <- '/share/ScratchGeneral/tonwan/projects/snPATHO-Seq/data/snPATHO_seq/25k_downsampled_processed_objs/integrated/4066_cellbender_filtered_integration_final_annotation.rds'


# visium_obj_file <- '/share/ScratchGeneral/tonwan/projects/snPATHO-Seq/data/Visium/processed/STutility/4411FFPE.rds'
# visium_obj_file <- '/share/ScratchGeneral/tonwan/projects/snPATHO-Seq/data/Visium/processed/STutility/4399FFPE.rds'
visium_obj_file <- '/share/ScratchGeneral/tonwan/projects/snPATHO-Seq/data/Visium/processed/STutility/4066FFPE.rds'


out_dir <- '/share/ScratchGeneral/tonwan/projects/snPATHO-Seq/results/deconvolution/RCTD'
dir.create(out_dir, recursive = T)

annotation_colname <- 'major_annotation'

batch_colname <- 'sample_id'

batch <- '4066SNAPFix'
# "4411FFPE"    "4411GEX" "4411SNAPFix" 
# "4399FFPE_run1" "4399GEX" "4399SNAPFix"
# "4066FFPE"    "4066GEX" "4066SNAPFix" 

# minimal numbers of cells in single-cell reference as input
min_cell <- 3


working_out <- paste0(out_dir, '/', batch)
dir.create(working_out, recursive = T)


########################################################################################################
########################################################################################################
########################################################################################################
# prepare single cell data ----
ref_obj <- readRDS(file = ref_obj_file)

# split the reference object based on the processing method / sample id ----
subset_ref_obj <- subset(ref_obj, subset = !!sym(batch_colname) == batch)

# make sure all cell types have at least the required amount
summary_df <- as.data.frame(table(subset_ref_obj@meta.data[[annotation_colname]]))
cell_types_to_keep <- as.character(summary_df[summary_df$Freq >= min_cell, 'Var1'])
subset_ref_obj <- subset(subset_ref_obj, subset = !!sym(annotation_colname) %in% cell_types_to_keep)


sc_counts <- GetAssayData(subset_ref_obj, assay = 'RNA', slot = 'counts')
sc_cell_types <- as.character(subset_ref_obj@meta.data[, annotation_colname])
names(sc_cell_types) <- rownames(subset_ref_obj@meta.data)
sc_cell_types <- as.factor(sc_cell_types)

sc_reference <- spacexr::Reference(
    counts = sc_counts,
    cell_types = sc_cell_types)


########################################################################################################
########################################################################################################
########################################################################################################
# prepare visium data for deconvolution ----
visium_obj <- readRDS(file = visium_obj_file)
v_counts <- GetAssayData(visium_obj, assay = 'Spatial', slot = 'counts')
image_obj <- GetStaffli(visium_obj)
v_coords <- image_obj@meta.data[, c('x', 'y')]
v_nUMI <- colSums(v_counts)

## Create SpatialRNA object ----
v_RCTD_obj <- spacexr::SpatialRNA(
    coords = v_coords, 
    counts = v_counts, 
    nUMI = v_nUMI)


########################################################################################################
########################################################################################################
########################################################################################################
# run RCTD ----
RCTD_res <- spacexr::create.RCTD(v_RCTD_obj, 
                                sc_reference,
                                CELL_MIN_INSTANCE = min_cell,
                                max_cores = 4)
RCTD_res <- spacexr::run.RCTD(RCTD_res, doublet_mode = 'full')

saveRDS(file = paste0(working_out, '/', batch, '_RCTD_results.rds'), RCTD_res)


########################################################################################################
########################################################################################################
########################################################################################################
# plot to check RCTD results ----

## add raw and scaled weights to visium object ----
raw_weights <- RCTD_res@results$weights
norm_weights <- spacexr::normalize_weights(raw_weights) 
norm_weights <- t(norm_weights)
scaled_weights_assay <- CreateAssayObject(data = as.matrix(norm_weights))
visium_obj[['RCTD']] <- scaled_weights_assay


# make spatial plot for results ----
plot_dir <- here(working_out, 'spatial_plots')
dir.create(plot_dir, recursive = T)


for (ct in rownames(visium_obj[['RCTD']])) {
  DefaultAssay(visium_obj) <- 'RCTD'
  plot <- ST.FeaturePlot(visium_obj, features = ct, 
                        grid.ncol = 1, show.sb = F,
                        pt.size = 2, pt.border = F,
                        cols = c("lightgray", "mistyrose", "red", "darkred", "black"))
  pdf(file = paste0(plot_dir, '/', ct, '_sclaed_porportion.pdf'),
      width = 6, height = 5)
      print(plot)
  dev.off()
}
```
