## Supplementary material for "snPATHO-seq: unlocking the pathology archives": scripts: 02_integrate_each_sample.R

```
# aim ----
    # take pre-processed data
    # integrate all data
    # derive joint cell type annotations
    # split by dataset


# define environment ----
.libPaths(
    c(
        '/share/ScratchGeneral/tonwan/projects/snPATHO-Seq/renv/library/R-4.1/x86_64-conda-linux-gnu',
        .libPaths()
    )
)
library(Seurat)
library(tidyverse)
library(RColorBrewer)


# arguments ----
data_dir <- '/share/ScratchGeneral/tonwan/projects/snPATHO-Seq/data/external_data/FHIL_Data/processed_data'

out_dir <- '/share/ScratchGeneral/tonwan/projects/snPATHO-Seq/results/snRNA_external_data/FHIL_kidney/integration'
dir.create(out_dir, recursive = T)

out_data_dir <- '/share/ScratchGeneral/tonwan/projects/snPATHO-Seq/data/external_data/FHIL_Data/integrated'
dir.create(out_data_dir, recursive = T)


##################################################################################################
##################################################################################################
##################################################################################################
# load data
obj_list <- list()
obj_list[[1]] <- readRDS(file = paste0(data_dir, '/', 'Kidney_nuclei_clustered.rds'))
obj_list[[2]] <- readRDS(file = paste0(data_dir, '/', 'Kidney_whole_cell_clustered.rds'))


# integrate objects
source('/share/ScratchGeneral/tonwan/projects/snPATHO-Seq/scripts/function_scripts/01_Seurat_integration.R')

integrated_obj <- seurat_integration(obj_list,
                                    normalisation_method = 'RNA',
                                    nfeatue_each_dataset = 2000,
                                    nfeatures_common = 2000,
                                    vars_to_regress = NULL,
                                    integration_method = 'cca',
                                    nthread = NULL)
    


# plot to check integration results

# load predefined colors
source('/share/ScratchGeneral/tonwan/projects/snPATHO-Seq/scripts/colors.R')
# only keep the selected workflow colors
working_colors <- workflow_colors[names(workflow_colors) %in% unique(integrated_obj$processing_method)]


# plot QC metrics
pdf(paste0(out_dir, '/', 'umap_batch_factors.pdf'))
    print(
        DimPlot(integrated_obj, reduction = "umap", group.by = 'processing_method')
    )
    print(
        FeaturePlot(integrated_obj, features = "nCount_RNA", reduction = 'umap') +
                scale_color_continuous(trans='log2', type = 'viridis')
    )
    print(
        FeaturePlot(integrated_obj, features = "nFeature_RNA", reduction = 'umap') +
                scale_color_continuous(trans='log2', type = 'viridis')
    )
    print(
        FeaturePlot(integrated_obj, features = "percent.mt", reduction = 'umap')
    )
dev.off()

# split UMAP by processing methods
pdf(file = paste0(out_dir, '/', 'umap_split_by_sample_paper_figure.pdf'),
        width = 10,
        height = 5)
        print(
            DimPlot(integrated_obj, reduction = "umap", group.by = 'processing_method', 
                split.by = 'processing_method', ncol = 2, cols = working_colors) +
                labs(title = NULL)
        )
dev.off()

    
    
    
    
    
    
    
    


##################################################################################################
##################################################################################################
##################################################################################################
# find a good clustering resolution
clust_dir <- paste0(out_dir, '/', 'cluster_optimisation')
dir.create(clust_dir, recursive = T)

# load function
source('/share/ScratchGeneral/tonwan/projects/snPATHO-Seq/scripts/function_scripts/02_Seurat_clustering_optimisation.R')

# make sure the default assay is the integrated assay
DefaultAssay(integrated_obj) <- 'integrated'
# cleanup metadta
integrated_obj@meta.data <- integrated_obj@meta.data[, c('orig.ident', 'nCount_RNA', 'nFeature_RNA', 'percent.mt',
                                                            'dataset_id', 'processing_method', 'sample_id')]
# cluster and find optimal clustering resolution
integrated_obj <- clustering_resolution_optimisation(integrated_obj,
                                        resolution_range = seq(0.2, 2, by = 0.1),
                                        plot_dir = clust_dir,
                                        batch_factor = "processing_method",
                                        run_silhouette = FALSE,
                                        dist.matrix = NULL)


##################################################################################################
##################################################################################################
##################################################################################################
# save processed data
saveRDS(file = paste0(out_data_dir, '/', 'Kidney_integrated.rds'), integrated_obj)
```
