## Supplementary material for "snPATHO-seq: unlocking the pathology archives": scripts: 01_integration.R

```
# aim ----
    # dimension reduction and clustering based FLEX and 3p compairson using PBMC data


# define environment ----
library(Seurat)
library(tidyverse)
library(RColorBrewer)


# arguments ----
data_dir <- '/share/ScratchGeneral/tonwan/projects/snPATHO-Seq/data/external_data/FHIL_Data/Aligned_30K_downsampled'

out_dir <- '/share/ScratchGeneral/tonwan/projects/snPATHO-Seq/results/snRNA_external_data/FHIL_PBMC'

out_data_dir <- '/share/ScratchGeneral/tonwan/projects/snPATHO-Seq/data/external_data/FHIL_Data/processed_data'


# load data ----
obj_list <- list()

mtx <- Read10X(paste0(data_dir, '/FRP_PBMC1/outs/per_sample_outs/FRP_PBMC1/count/sample_filtered_feature_bc_matrix'))
obj <- CreateSeuratObject(counts = mtx)
obj$sample_id <- 'FLEX_exp_1'
obj_list[['FLEX_exp_1']] <- obj


mtx <- Read10X(paste0(data_dir, '/FRP_PBMC2/outs/per_sample_outs/FRP_PBMC2/count/sample_filtered_feature_bc_matrix'))
obj <- CreateSeuratObject(counts = mtx)
obj$sample_id <- 'FLEX_exp_2'
obj_list[['FLEX_exp_2']] <- obj


mtx <- Read10X(paste0(data_dir, '/V31_PBMC1/outs/filtered_feature_bc_matrix'))
obj <- CreateSeuratObject(counts = mtx)
obj$sample_id <- '3p_exp_1'
obj_list[['3p_exp_1']] <- obj


mtx <- Read10X(paste0(data_dir, '/V31_PBMC2/outs/filtered_feature_bc_matrix'))
obj <- CreateSeuratObject(counts = mtx)
obj$sample_id <- '3p_exp_2'
obj_list[['3p_exp_2']] <- obj


#####################################################################################################################
#####################################################################################################################
#####################################################################################################################
# filtering
filtered_obj_list <- lapply(obj_list, function(x){
    x <- subset(x, subset = nFeature_RNA > 200)
    x <- subset(x, subset = nFeature_RNA < 8000)
    x[["percent.mt"]] <- PercentageFeatureSet(x, pattern = "^MT-")
    x <- subset(x, subset = percent.mt < 10)
})


#####################################################################################################################
#####################################################################################################################
#####################################################################################################################
# integration
source('/share/ScratchGeneral/tonwan/projects/snPATHO-Seq/scripts/function_scripts/01_Seurat_integration.R')

# integrate data 
integrated_obj <- seurat_integration(filtered_obj_list,
                                    normalisation_method = 'RNA',
                                    nfeatue_each_dataset = 2000,
                                    nfeatures_common = 2000,
                                    vars_to_regress = NULL,
                                    integration_method = 'cca',
                                    nthread = NULL)


####################################################################################################
####################################################################################################
####################################################################################################
working_dir <- paste0(out_dir, '/', 'dimension_reduction')
dir.create(working_dir, recursive = T)

# check integration quality and annotate clusters
integrated_obj$cells_with_low_gene_count <- ifelse(integrated_obj$nFeature_RNA < 500, 'yes', 'no')
integrated_obj$cells_with_high_gene_count <- ifelse(integrated_obj$nFeature_RNA > 5000, 'yes', 'no')

# plot check batch factors and QC values
pdf(paste0(working_dir, '/', 'umap_batch_factors.pdf'))
    print(
        DimPlot(integrated_obj, reduction = "umap", group.by = 'sample_id')
    )
    print(
        FeaturePlot(integrated_obj, features = "nCount_RNA", reduction = 'umap') +
            scale_color_continuous(trans='log2', type = 'viridis')
    )
    print(
        FeaturePlot(integrated_obj, features = "nFeature_RNA", reduction = 'umap') +
            scale_color_continuous(trans='log2', type = 'viridis')
    )
    print(
        FeaturePlot(integrated_obj, features = "percent.mt", reduction = 'umap')
    )
    print(
        DimPlot(integrated_obj, reduction = "umap", group.by = 'cells_with_low_gene_count')
    )
    print(
        DimPlot(integrated_obj, reduction = "umap", group.by = 'cells_with_high_gene_count')
    )
dev.off()


# split umap by sample
pdf(file = paste0(working_dir, '/', 'umap_split_by_sample_id.pdf'),
    width = 20,
    height = 5)
    print(
        DimPlot(integrated_obj, reduction = "umap", group.by = 'sample_id', 
            split.by = 'sample_id', ncol = 4)
    )
dev.off()


####################################################################################################
####################################################################################################
####################################################################################################
# find optimal clustering resolution ----
clust_dir <- paste0(out_dir, '/', 'dimension_reduction', '/', 'cluster_optimisation')
dir.create(clust_dir, recursive = T)

# load function
source('/share/ScratchGeneral/tonwan/projects/snPATHO-Seq/scripts/function_scripts/02_Seurat_clustering_optimisation.R')

# make sure the default assay is the integrated assay
DefaultAssay(integrated_obj) <- 'integrated'

# cluster and find optimal clustering resolution
integrated_obj <- clustering_resolution_optimisation(integrated_obj,
                                        resolution_range = seq(0.2, 2, by = 0.1),
                                        plot_dir = clust_dir,
                                        batch_factor = "sample_id",
                                        run_silhouette = FALSE,
                                        dist.matrix = NULL)


####################################################################################################
####################################################################################################
####################################################################################################
# save processed data
saveRDS(file = paste0(out_data_dir, '/', 'PBMC_integrated_obj.rds'), integrated_obj)
```
