## Supplementary material for "snPATHO-seq: unlocking the pathology archives": scripts: 02_manual_cell_type_annotation.R

```
# aim ----
    # annotate cell types by top markers


# define environment ----
library(Seurat)
library(tidyverse)
library(RColorBrewer)


# arguments ----
data_dir <- '/share/ScratchGeneral/tonwan/projects/snPATHO-Seq/data/external_data/FHIL_Data/processed_data'

out_dir <- '/share/ScratchGeneral/tonwan/projects/snPATHO-Seq/results/snRNA_external_data/FHIL_PBMC'
dir.create(out_dir, recursive = T)


####################################################################################################
####################################################################################################
####################################################################################################
# load data ----
obj <- readRDS(paste0(data_dir, '/', 'PBMC_integrated_obj.rds'))


####################################################################################################
####################################################################################################
####################################################################################################
# plot some markers ----
marker_dir <- paste0(out_dir, '/', 'markers')
dir.create(marker_dir, recursive = T)

proliferating <- c('MKI67')
t_cell_general <- c('CD3D', 'CD3E', 'CD4', 'CD8A', 'CD8B')
ab_t_cells <- c('TRAC', 'TRBC1')
gd_t_cells <- c('TRDC', 'TRGC1', 'TRGC2', 'KLRC1', 'NKG7', 'TRDV2', 'CD7', 'TRGV9', 'KLRD1', 'KLRG1')
MAIT <- c('KLRB1', 'TRAV1-2', 'TRAJ33', 'IFNG', 'NKG7', 'GZMK', 'IL7R', 'SLC4A10', 'GZMA', 'CXCR6', 'PRSS35', 'RBM24', 'NCR3')
dnT <- c('PTPN3', 'MIR4422HG', 'NUCB2', 'CAV1', 'DTHD1', 'GZMA', 'MYB', 'FXYD2', 'GZMK', 'AC004585.1')

CD8_t_cells <- c('CD8A', 'CD8B')
naive_CD8 <- c('CCR7', 'IL7R', 'S100B', 'RGS10', 'NOSIP', 'LINC02446', 'LEF1', 'CRTAM', 'OXNAD1')
CD8_TCM <- c('IL7R', 'CD27', 'SELL', 'CCR7', 'ANXA1', 'KRT1', 'LINC02446', 'YBX3', 'NELL2', 'LDHB')
CD8_TEM <- c('IL7R', 'CX3CR1', 'GZMA', 'PRF1', 'CCR5', 'CCL5', 'GZMH', 'KLRD1', 'NKG7', 'GZMK', 'CST7', 'TRGC2')

CD4_t_cells <- c('CD4')
naive_CD4 <- c('CCR7', 'TCF7', 'IL7R', 'FHIT', 'LEF1', 'MAL', 'NOSIP', 'LDHB', 'PIK3IP1')
CD4_TCM <- c('IL2RA', 'SELL', 'CCR7', 'IL7R', 'TMSB10', 'ITGB1', 'LTB', 'AQP3', 'LDHB', 'IL32', 'MAL')
CD4_TEM <- c('IL2RA', 'CCR5', 'IL7R', 'CCL5', 'FYB1', 'GZMK', 'IL32', 'GZMA', 'KLRB1', 'LTB', 'AQP3')
CD4_CTL <- c('GZMH', 'FGFBP2', 'ITGB1', 'GZMA', 'CST7', 'GNLY', 'B2M', 'IL32', 'NKG7')
Treg <- c('IL2RA', 'FOXP3')

NK <- c('NCAM1', 'CD244', 'GNLY', 'TYROBP', 'NKG7', 'FCER1G', 'GZMB', 'TRDC', 'PRF1', 'FGFBP2', 'SPON2', 'KLRF1')
NK_CD56_bright <- c('XCL2', 'FCER1G', 'SPINK2', 'TRDC', 'KLRC1', 'XCL1', 'SPTSSB', 'PPP1R9A', 'NCAM1', 'TNFRSF11A')

CD14_monocytes <- c('CD14', 'S100A9', 'CTSS', 'S100A8', 'LYZ', 'VCAN', 'S100A12', 'IL1B', 'G0S2', 'FCN1')
CD16_monocytes <- c('FCGR3A', 'CDKN1C', 'PTPRC', 'LST1', 'IER5', 'MS4A7', 'RHOC', 'IFITM3', 'AIF1', 'HES4')

ASDC <- c('PPP1R14A', 'LILRA4', 'AXL', 'IL3RA', 'SCT', 'SCN9A', 'LGMN', 'DNASE1L3', 'CLEC4C', 'GAS6')
cDC1 <- c('CLEC9A', 'XCR1', 'DNASE1L3', 'C1orf54', 'IDO1', 'CLNK', 'CADM1', 'FLT3', 'ENPP1', 'NDRG2')
cDC2 <- c('CD1C', 'FCER1A', 'HLA-DQA1', 'CLEC10A', 'ENHO', 'PLD4', 'GSN', 'SLC38A1', 'NDRG2', 'AFF3')
pDC <- c('IL3RA', 'TLR7', 'TLR9', 'JCHAIN', 'ITM2C', 'PLD4', 'SERPINF1', 'LILRA4', 'TPM2', 'MZB1', 'SPIB', 'IRF4', 'SMPD3')

HSPC <- c('SPINK2', 'PRSS57', 'CYTL1', 'EGFL7', 'GATA2', 'CD34', 'SMIM24', 'AVP', 'MYB', 'LAPTM4B')
ILC <- c('KIT', 'TRDC', 'TTLL10', 'LINC01229', 'SOX4', 'KLRB1', 'TNFRSF18', 'TNFRSF4', 'IL1R1', 'HPGDS')


markers <- c(proliferating, t_cell_general, ab_t_cells, gd_t_cells, MAIT, dnT, 
    CD8_t_cells, naive_CD8, CD8_TCM, CD8_TEM,
    CD4_t_cells, naive_CD4, CD4_TCM, CD4_TEM, CD4_CTL, Treg,
    NK, NK_CD56_bright, CD14_monocytes, CD16_monocytes,
    ASDC, cDC1, cDC2, pDC, HSPC, ILC
)


DefaultAssay(obj) <- 'RNA'
to_plot <- markers[markers %in% rownames(obj)]
for (g in to_plot) {
    pdf(paste0(marker_dir, '/', g, '.pdf'))
        print(FeaturePlot(obj, features = g, reduction = 'umap', order = T))
    dev.off()
}
 


####################################################################################################
####################################################################################################
####################################################################################################
# annotate subclusters
optimal_clustering_dir <- paste0(out_dir, '/', 'optimal_cluster')
dir.create(optimal_clustering_dir, recursive = T)


opt_res <- 0.8
cluster_name <- grep(paste0('res.', opt_res), colnames(obj@meta.data), value = T)[1]


# make a plot with label at the optimal cluster resolution ----
pdf(file = paste0(optimal_clustering_dir, '/', 'umap_', opt_res, '.pdf'))
    print(
        DimPlot(obj, reduction = "umap", group.by = cluster_name, label = T, repel = T)
    )
dev.off()


# check top markers of a few clusters 
de_dir <- paste0(optimal_clustering_dir, '/', 'de_res')
dir.create(de_dir, recursive = T)

cl_to_check <- c(5, 19, 8, 10, 15)

Idents(obj) <- cluster_name
for (cl in cl_to_check) {
    de_res <- FindMarkers(obj, ident.1 = cl, only.pos = TRUE)
    de_res <- de_res %>% 
        filter(p_val_adj < 0.05)
    write.csv(file = paste0(de_dir, '/', cl, '_de_markers.csv'), de_res)
}


####################################################################################
####################################################################################
####################################################################################
# annotate clusters ----
Idents(obj) <- cluster_name
annotation_col_name <- 'manual_annotation'

new_annotation_df <- data.frame(cluster_id = unique(as.character(Idents(obj))))
new_annotation_df[[annotation_col_name]] <- ''

new_annotation_df[new_annotation_df$cluster_id == '0', annotation_col_name] <- 'CD4_naive'
new_annotation_df[new_annotation_df$cluster_id == '1', annotation_col_name] <- 'B_naive'
new_annotation_df[new_annotation_df$cluster_id == '2', annotation_col_name] <- 'CD14_monocyte'
new_annotation_df[new_annotation_df$cluster_id == '3', annotation_col_name] <- 'NK'
new_annotation_df[new_annotation_df$cluster_id == '4', annotation_col_name] <- 'CD8_TEM_CD4_CTL'
new_annotation_df[new_annotation_df$cluster_id == '5', annotation_col_name] <- 'CD4_TCM'
new_annotation_df[new_annotation_df$cluster_id == '6', annotation_col_name] <- 'CD8_naive'
new_annotation_df[new_annotation_df$cluster_id == '7', annotation_col_name] <- 'B_intermediate_memory'
new_annotation_df[new_annotation_df$cluster_id == '8', annotation_col_name] <- 'CD4_TCM'
new_annotation_df[new_annotation_df$cluster_id == '9', annotation_col_name] <- 'CD8_TCM_CD8_naive'
new_annotation_df[new_annotation_df$cluster_id == '10', annotation_col_name] <- 'CD4_TEM'
new_annotation_df[new_annotation_df$cluster_id == '11', annotation_col_name] <- 'CD14_monocyte'
new_annotation_df[new_annotation_df$cluster_id == '12', annotation_col_name] <- 'MAIT'
new_annotation_df[new_annotation_df$cluster_id == '13', annotation_col_name] <- 'Treg'
new_annotation_df[new_annotation_df$cluster_id == '14', annotation_col_name] <- 'CD8_NK_mixed_CTL_population'
new_annotation_df[new_annotation_df$cluster_id == '15', annotation_col_name] <- 'CD4_TCM'
new_annotation_df[new_annotation_df$cluster_id == '16', annotation_col_name] <- 'CD8_TCM'
new_annotation_df[new_annotation_df$cluster_id == '17', annotation_col_name] <- 'CD16_monocyte'
new_annotation_df[new_annotation_df$cluster_id == '18', annotation_col_name] <- 'CD4_naive'
new_annotation_df[new_annotation_df$cluster_id == '19', annotation_col_name] <- 'CD4_TCM'
new_annotation_df[new_annotation_df$cluster_id == '20', annotation_col_name] <- 'CD8_naive'
new_annotation_df[new_annotation_df$cluster_id == '21', annotation_col_name] <- 'cDC'
new_annotation_df[new_annotation_df$cluster_id == '22', annotation_col_name] <- 'NK_CD56bright_HSPC'
new_annotation_df[new_annotation_df$cluster_id == '23', annotation_col_name] <- 'CD4_naive'
new_annotation_df[new_annotation_df$cluster_id == '24', annotation_col_name] <- 'platelet'
new_annotation_df[new_annotation_df$cluster_id == '25', annotation_col_name] <- 'CD14_monocyte'
new_annotation_df[new_annotation_df$cluster_id == '26', annotation_col_name] <- 'CD14_monocyte'
new_annotation_df[new_annotation_df$cluster_id == '27', annotation_col_name] <- 'ILC'
new_annotation_df[new_annotation_df$cluster_id == '28', annotation_col_name] <- 'pDC'


# annotate the object
new_annotations <- new_annotation_df[[annotation_col_name]]
names(new_annotations) <- new_annotation_df$cluster_id

obj <- RenameIdents(obj, new_annotations)
obj@meta.data[[annotation_col_name]] <- Idents(obj)


# export cell type annotations as csv file for easier modification and assessemnt
write.csv(file = paste0(optimal_clustering_dir, '/', 'manual_annotation.csv'), obj@meta.data[[annotation_col_name]])


# plot a umap with new annotations
Idents(obj) <- annotation_col_name
pdf(file = paste0(optimal_clustering_dir, '/', 'umap_manual_annotation.pdf'), width = 10, height = 7)
    print(
        DimPlot(obj, reduction = "umap", group.by = annotation_col_name, label = T, repel = T)
    )
dev.off()


####################################################################################
####################################################################################
####################################################################################
# save annotated object
saveRDS(file = paste0(data_dir, '/', 'PBMC_integrated_manual_annotation.rds'), obj)
```
