## Supplementary material for "snPATHO-seq: unlocking the pathology archives": scripts: subcluster_B.R

```
# aim ----
    # subcluster B cells to better differentiate them
    # redo integration for these cells


# define environment ----
library(Seurat)
library(tidyverse)
library(RColorBrewer)


# arguments ----
data_dir <- '/share/ScratchGeneral/tonwan/projects/snPATHO-Seq/data/external_data/FHIL_Data/processed_data'


out_dir <- '/share/ScratchGeneral/tonwan/projects/snPATHO-Seq/results/snRNA_external_data/FHIL_PBMC'
dir.create(out_dir, recursive = T)


####################################################################################################
####################################################################################################
####################################################################################################
# load data ----
obj <- readRDS(paste0(data_dir, '/', 'PBMC_integrated_manual_annotation.rds'))


# subset to only keep cells of interest
subset_obj <- subset(obj, subset = manual_annotation %in% c('B_intermediate_memory', 'B_naive'))


####################################################################################################
####################################################################################################
####################################################################################################
# simply recluster data
working_dir <- paste0(out_dir, '/', 'reclustering', '/', 'B_cells')
dir.create(working_dir, recursive = T)


# simply recluster cells ----
DefaultAssay(subset_obj) <- 'RNA'
subset_obj <- FindVariableFeatures(subset_obj, nfeatures = 5000)
featueres <- VariableFeatures(subset_obj)

DefaultAssay(subset_obj) <- 'integrated'
subset_obj <- ScaleData(subset_obj, features = featueres)
subset_obj <- RunPCA(subset_obj, features = featueres)
subset_obj <- RunUMAP(subset_obj, dims = 1:30)
subset_obj <- FindNeighbors(subset_obj, dims = 1:30)


# find optimal clustering resolution
clustering_optimisation_dir <- paste0(working_dir, '/', 'clustering_optimisation')
dir.create(clustering_optimisation_dir, recursive = T)

for (res in seq(0.2, 2, by = 0.1)) {
    subset_obj <- FindClusters(subset_obj, resolution = res)
    cluster_name <- grep(paste0('res.', res), colnames(subset_obj@meta.data), value = T)[1]
    # make a plot with label at the optimal cluster resolution ----
    pdf(file = paste0(clustering_optimisation_dir, '/', 'umap_', res, '.pdf'))
        print(
            DimPlot(subset_obj, reduction = "umap", group.by = cluster_name, label = T, repel = T)
        )
    dev.off()

    pdf(file = paste0(clustering_optimisation_dir, '/', 'umap_split_by_clusters_', res, '.pdf'), 
        width = 20, 
        height = ceiling(length(unique(subset_obj@meta.data[[cluster_name]])) / 4) * 5)
        print(
            DimPlot(subset_obj, reduction = "umap", group.by = cluster_name, 
                split.by = cluster_name, ncol = 4)
        )
    dev.off()
}


# check batch 
pdf(file = paste0(working_dir, '/', 'batch.pdf'))
    print(
        DimPlot(subset_obj, group.by = 'sample_id')
    )
dev.off()


# plot some markers 
marker_dir <- paste0(working_dir, '/', 'markers')
dir.create(marker_dir, recursive = T)

intermediate_B <- c('MS4A1', 'TNFRSF13B', 'IGHM', 'IGHD', 'AIM2', 'CD79A', 'LINC01857', 'RALGPS2', 'BANK1', 'CD79B')
memory_B <- c('MS4A1', 'COCH', 'AIM2', 'BANK1', 'SSPN', 'CD79A', 'TEX9', 'RALGPS2', 'TNFRSF13C', 'LINC01781')
naive_B <- c('IGHM', 'IGHD', 'CD79A', 'IL4R', 'MS4A1', 'CXCR4', 'BTG1', 'TCL1A', 'CD79B', 'YBX3')
plasmablast <- c('IGHA2', 'MZB1', 'TNFRSF17', 'DERL3', 'TXNDC5', 'TNFRSF13B', 'POU2AF1', 'CPNE5', 'HRASLS2', 'NT5DC2')


markers <- c(intermediate_B, memory_B, naive_B, plasmablast, 'TRAC')

DefaultAssay(subset_obj) <- 'RNA'
to_plot <- unique(markers[markers %in% rownames(subset_obj)])
for (g in to_plot) {
    pdf(paste0(marker_dir, '/', g, '.pdf'))
        print(FeaturePlot(subset_obj, features = g, reduction = 'umap', order = T))
    dev.off()
}


####################################################################################################
####################################################################################################
####################################################################################################
# redo cluster annotation - on subset object
Idents(subset_obj) <- grep(paste0('res.', 0.2), colnames(subset_obj@meta.data), value = T)[1]
annotation_col_name <- 'modified_annotation_by_subclustering'

new_annotation_df <- data.frame(cluster_id = unique(as.character(Idents(subset_obj))))
new_annotation_df[[annotation_col_name]] <- ''

new_annotation_df[new_annotation_df$cluster_id == '0', annotation_col_name] <- 'B_naive'
new_annotation_df[new_annotation_df$cluster_id == '1', annotation_col_name] <- 'B_intermediate'
new_annotation_df[new_annotation_df$cluster_id == '2', annotation_col_name] <- 'B_memory'
new_annotation_df[new_annotation_df$cluster_id == '3', annotation_col_name] <- 'doublet'
new_annotation_df[new_annotation_df$cluster_id == '4', annotation_col_name] <- 'B_naive'


# annotate the object
new_annotations <- new_annotation_df[[annotation_col_name]]
names(new_annotations) <- new_annotation_df$cluster_id

subset_obj <- RenameIdents(subset_obj, new_annotations)
subset_obj@meta.data[[annotation_col_name]] <- Idents(subset_obj)


# export cell type annotations as csv file for easier modification and assessemnt
write.csv(file = paste0(working_dir, '/', 'modified_annotation.csv'), 
    subset_obj@meta.data[, annotation_col_name, drop = F])


# plot a umap with new annotations
Idents(subset_obj) <- annotation_col_name
pdf(file = paste0(working_dir, '/', 'umap_modified_annotation.pdf'), width = 10, height = 7)
    print(
        DimPlot(subset_obj, reduction = "umap", group.by = annotation_col_name, label = T, repel = T)
    )
dev.off()
```
