## Supplementary material for "snPATHO-seq: unlocking the pathology archives": scripts: 04_final_annotation.R

```
# aim ----
    # merge annotation modified through subclustering


# define environment ----
library(Seurat)
library(tidyverse)
library(RColorBrewer)


# arguments ----
data_obj_file <- '/share/ScratchGeneral/tonwan/projects/snPATHO-Seq/data/external_data/FHIL_Data/processed_data/PBMC_integrated_manual_annotation.rds'


# load object ----
obj <- readRDS(file = data_obj_file)


################################################################################################################
################################################################################################################
################################################################################################################
# modify old annotation ----
new_CD8_TCM_CD8_naive_annotation <- read.csv(file = '/share/ScratchGeneral/tonwan/projects/snPATHO-Seq/results/snRNA_external_data/FHIL_PBMC/reclustering/CD8_TCM_CD8_naive/modified_annotation.csv',
    row.names = 1)
new_B_intermediate_memory_annotation <- read.csv(file = '/share/ScratchGeneral/tonwan/projects/snPATHO-Seq/results/snRNA_external_data/FHIL_PBMC/reclustering/B_cells/modified_annotation.csv',
    row.names = 1)
new_NK_CD56bright_HSPC_annotation <- read.csv(file = '/share/ScratchGeneral/tonwan/projects/snPATHO-Seq/results/snRNA_external_data/FHIL_PBMC/reclustering/NK_CD56bright_HSPC/modified_annotation.csv',
    row.names = 1)
new_CD8_NK_mixed_CTL_population_annotation <- read.csv(file = '/share/ScratchGeneral/tonwan/projects/snPATHO-Seq/results/snRNA_external_data/FHIL_PBMC/reclustering/CD8_NK_mixed_CTL_population/modified_annotation.csv',
    row.names = 1)
new_CD8_TEM_CD4_CTL_annotation <- read.csv(file = '/share/ScratchGeneral/tonwan/projects/snPATHO-Seq/results/snRNA_external_data/FHIL_PBMC/reclustering/CD8_TEM_CD4_CTL/modified_annotation.csv',
    row.names = 1)


new_annotation_df <- do.call(rbind, 
    list(new_CD8_TCM_CD8_naive_annotation,
        new_B_intermediate_memory_annotation,
        new_NK_CD56bright_HSPC_annotation,
        new_CD8_NK_mixed_CTL_population_annotation,
        new_CD8_TEM_CD4_CTL_annotation))
 
new_annotation_df <- base::merge(obj@meta.data[, 'manual_annotation', drop = F], new_annotation_df, by = 0, all.x = TRUE)
new_annotation_df <- column_to_rownames(new_annotation_df, var = 'Row.names')
new_annotation_df <- new_annotation_df[rownames(obj@meta.data), ]
new_annotation_df$modified_annotation_by_subclustering <- ifelse(is.na(new_annotation_df$modified_annotation_by_subclustering), as.character(new_annotation_df$manual_annotation), new_annotation_df$modified_annotation_by_subclustering)


obj$new_annotation <- new_annotation_df$modified_annotation_by_subclustering
    


################################################################################################################
################################################################################################################
################################################################################################################
# save modified data object 
saveRDS(file = '/share/ScratchGeneral/tonwan/projects/snPATHO-Seq/data/external_data/FHIL_Data/processed_data/PBMC_integrated_annotation_modifed_by_subclustering.rds',
    obj)
```
