## Supplementary material for "snPATHO-seq: unlocking the pathology archives": scripts: 05_paper_figures.R

```
# aim ----
    # make plots for paper


# define environment ----
library(Seurat)
library(tidyverse)
library(RColorBrewer)

library(ComplexHeatmap)
library(circlize)


# arguments ----
data_obj_file <- '/share/ScratchGeneral/tonwan/projects/snPATHO-Seq/data/external_data/FHIL_Data/processed_data/PBMC_integrated_annotation_modifed_by_subclustering.rds'

out_dir <- '/share/ScratchGeneral/tonwan/projects/snPATHO-Seq/results/snRNA_external_data/FHIL_PBMC/final_annotation'


# load data ----
obj <- readRDS(file = data_obj_file)


################################################################################################################
################################################################################################################
################################################################################################################
# plot UMI / gene per cell

# define workflow colors
colors <- c("#1B9E77", "#D95F02")

# plot UMI per cell
plot <- ggplot(merged_obj@meta.data, aes(x = unique_id, y = nCount_RNA, fill = processing_method)) +
    geom_violin() +
    scale_fill_manual(values = colors) +
    geom_boxplot(fill = 'white', width=0.1, outlier.size = 0) +
    facet_grid(cols = vars(sample_id), scales="free") +
    stat_summary(fun = 'median', colour = "black", size = 10,
                 geom = "text", aes(y = stage(start = nCount_RNA, after_stat = 120000), label = round(after_stat(y)))) +
    scale_y_log10() +
    labs(x = 'Samples',
        y = 'UMIs detected per cell',
        fill = 'Processing method',
        title = NULL) +
    theme_bw() +
    theme(text = element_text(size = 30),
        axis.text.x = element_blank())

ggsave(file = paste0(out_dir, '/', 'UMI_per_cell.pdf'), plot, width = 14, height = 7, device = 'pdf')


# plot gene per cell
plot <- ggplot(merged_obj@meta.data, aes(x = unique_id, y = nFeature_RNA, fill = processing_method)) +
    geom_violin() +
    scale_fill_manual(values = colors) +
    geom_boxplot(fill = 'white', width=0.1, outlier.size = 0) +
    facet_grid(cols = vars(sample_id), scales="free") +
    stat_summary(fun = 'median', colour = "black", size = 10,
                 geom = "text", aes(y = stage(start = nFeature_RNA, after_stat = 10000), label = round(after_stat(y)))) +
    # scale_y_log10() +
    labs(x = 'Samples',
        y = 'Genes detected per cell',
        fill = 'Processing method',
        title = NULL) +
    theme_bw() +
    theme(text = element_text(size = 30),
        axis.text.x = element_blank())

ggsave(file = paste0(out_dir, '/', 'Gene_per_cell.pdf'), plot, width = 14, height = 7, device = 'pdf')


################################################################################################################
################################################################################################################
################################################################################################################
# plot umap


# use customised function to pre set color for each cell type before plotting
source('/share/ScratchGeneral/tonwan/projects/snPATHO-Seq/scripts/function_scripts/08_prepare_data_for_complex_heatmap.R')

cell_type_colors <- set_colors(obj@meta.data, 'new_annotation', n = length(unique(obj$new_annotation)))


# plot UMAP
pdf(file = paste0(out_dir, '/', 'umap_final_annotation.pdf'), width = 10)
    print(
        DimPlot(obj, group.by = 'new_annotation', label = TRUE, repel = TRUE, cols = as.character(cell_type_colors)) +
            labs(title = NULL)
    )
dev.off()


# plot UMAP split by sample id and processing methods
colors <- c("#1B9E77", "#D95F02")

obj$processing_method <- ifelse(obj$sample_id %in% c('FLEX_exp_1', 'FLEX_exp_2'), 'FLEX', '3p')
obj$plot_id <- ifelse(obj$sample_id == 'FLEX_exp_1', 'Sample_1_FLEX',
    ifelse(obj$sample_id == 'FLEX_exp_2', 'Sample_2_FLEX',
    ifelse(obj$sample_id == '3p_exp_1', 'Sample_1_3p',
    ifelse(obj$sample_id == '3p_exp_2', 'Sample_2_3p', 'NULL'))))


pdf(file = paste0(out_dir, '/', 'umap_split_by_id_&_processing_method.pdf'), width = 20, height = 5)
    print(
        DimPlot(obj, group.by = 'processing_method', split.by = 'plot_id', cols = colors) +
            labs(title = NULL)
    )
dev.off()


################################################################################################################
################################################################################################################
################################################################################################################
# check top DE markers ----
Idents(obj) <- 'new_annotation'
de_res <- FindAllMarkers(obj, assay = 'RNA', test.use = 't', only.pos = TRUE)
write.csv(file = paste0(out_dir, '/', 'de_results.csv'), de_res)


# plot expression of top DE genes between clusters - annotate canonical cell type markers
# get top 200 DE genes from each cluster to plot
top_markers <- de_res %>% 
    group_by(cluster) %>%
    filter(p_val_adj < 0.05) %>%
    slice_max(avg_log2FC, n = 200) %>% 
    pull(gene) %>% 
    unique()


# get mean expresion of the selected genes for each processing method 
plot_mtx <- as.data.frame(AverageExpression(obj, assays = 'RNA', slot = 'data', features = top_markers, group.by = c('processing_method', 'new_annotation'))[[1]])
plot_mtx$gene_names <- rownames(plot_mtx)
plot_mtx_long <- gather(plot_mtx, key = 'annotation', value = 'mean_expression', -gene_names)

# annotate the expression dataframe
plot_mtx_long$processing_method <- unlist(lapply(str_split(plot_mtx_long$annotation, '_'), '[[', 1))
plot_mtx_long$cell_type <- unlist(lapply(str_split(plot_mtx_long$annotation, '\\_'), function(x) paste(x[-1], collapse = '_')))


# scale the gene expression within each processing method
plot_mtx_long <- plot_mtx_long %>% 
        group_by(gene_names, processing_method) %>%
        mutate(scale(mean_expression)) %>% 
        as.data.frame() 
  
# plot_order <- plot_mtx_long %>% 
#     arrange(as.factor(cell_type)) %>% 
#     pull(annotation) %>% 
#     unique()

plot_order <- plot_mtx_long %>% 
    arrange(factor(cell_type, 
        levels = c(
            'HSPC', 'cDC', 'CD14_monocyte', 'CD16_monocyte', 'pDC', 'platelet', 'ILC',
            'Treg', 'CD4_naive', 'CD8_naive', 'CD4_TCM', 'CD8_TCM', 'CD4_TEM', 'CD8_TEM',
            'CD4_CTL', 'MAIT', 'NK_CD56_bright', 'NK', 'CTL_CCR7_high', 'B_naive', 
            'B_intermediate', 'B_memory', 'doublet'))) %>% 
    pull(annotation) %>% 
    unique()


scaled_mtx <- spread(plot_mtx_long[, c('gene_names', 'annotation', 'scale(mean_expression)')],
        value = 'scale(mean_expression)', key = 'annotation')
scaled_mtx <- column_to_rownames(scaled_mtx, var = 'gene_names')
# cap the scaled expression at -5 to 5
scaled_mtx[scaled_mtx > 5] <- 5
scaled_mtx[scaled_mtx < -5] <- -5
scaled_mtx[is.na(scaled_mtx)] <- 0
scaled_mtx <- scaled_mtx[, plot_order]

# make heatmap annotation
annot_df <- plot_mtx_long[, c('annotation', 'processing_method', 'cell_type')]
annot_df <- distinct(annot_df)
annot_df <- annot_df[match(colnames(scaled_mtx), annot_df$annotation), ]


# set colors
col_fun <- colorRamp2(c(-5, 0, 5), 
                      c("#2166AC", "white", "#B2182B"))
source('/share/ScratchGeneral/tonwan/projects/snPATHO-Seq/scripts/colors.R')
workflow_colors <- workflow_colors[names(workflow_colors) %in% unique(annot_df$processing_method)]

# make annotation
left_annot <- HeatmapAnnotation(
        Workflows = annot_df$processing_method,
        which = 'row',
        col = list(Workflows = workflow_colors), 
        annotation_name_gp = grid::gpar(fontsize = 12),
        annotation_legend_param = list(
            Workflows = list(title_gp = gpar(fontsize = 12, fontface = "bold"), 
                                    labels_gp = gpar(fontsize = 12))))


# annotate selected genes 
genes_to_plot <- de_res %>% 
    group_by(cluster) %>%
    filter(p_val_adj < 0.05) %>%
    slice_max(avg_log2FC, n = 5) %>% 
    pull(gene)

# make sure the selected markers are in the matrix
all_markers <- genes_to_plot[genes_to_plot %in% rownames(scaled_mtx)]

# get location of the genes to be annotated
mtx_index <- seq(1:nrow(scaled_mtx))
names(mtx_index) <- rownames(scaled_mtx)
position_in_matrix <- mtx_index[names(mtx_index) %in% all_markers]

bottom_annotation <- HeatmapAnnotation(
    markers = anno_mark(
        at = position_in_matrix, 
        labels = names(position_in_matrix),
        which = "column", side = "bottom",
        labels_gp = gpar(fontsize = 8)))


# plot heatmap ----
plot <- Heatmap(t(scaled_mtx),
                name = 'Scaled mean expression',
                heatmap_legend_param = list(title_gp = gpar(fontsize = 12, fontface = "bold")),
                col = col_fun,
                left_annotation = left_annot,
                bottom_annotation = bottom_annotation,
                show_column_names= FALSE,
                show_row_names = TRUE,
                cluster_rows = FALSE,
                cluster_columns = TRUE,
                split = annot_df$processing_method,
                column_names_gp = gpar(fontsize = 6),
                row_names_gp = gpar(fontsize = 8))


pdf(file = paste0(out_dir, '/', 'top_DE_gene_heatmap_plot_200_annotate_5', '.pdf'),
    width = 14, height = 14)
    draw(plot)
dev.off()
```
