## Supplementary material for "snPATHO-seq: unlocking the pathology archives": scripts: 06_annotated_Hao_etal_with_inhouse_data.R

```
# aim ----
    # use current 3p and FLEX cell type signatures to annotate 3p data from Hao_et_al_2021 
    # use SCN for annotation
    # then compare predicted label to ground truth


# define environment ----
library(Seurat)
library(tidyverse)
library(RColorBrewer)


# arguments ----
training_data_file <- "/share/ScratchGeneral/tonwan/projects/snPATHO-Seq/data/external_data/FHIL_Data/processed_data/PBMC_integrated_annotation_modifed_by_subclustering.rds"

query_data_file <- '/share/ScratchGeneral/tonwan/projects/snPATHO-Seq/data/external_data/Hao_et_al_2021_PBMC/baseline_3p_obj.rds'

out_dir <- '/share/ScratchGeneral/tonwan/projects/snPATHO-Seq/results/snRNA_external_data/FHIL_PBMC/SCN_Hao_et_al_2021_annotation'


# load data ----
training_obj <- readRDS(file = training_data_file)

query_obj <- readRDS(file = query_data_file)


# load FLEX reference to get the whole panel
Flex_probe_ref <- read.csv(file = '/share/ScratchGeneral/tonwan/projects/snPATHO-Seq/resources/Chromium_Human_Transcriptome_Probe_Set_v1.0_GRCh38-2020-A.csv',
    skip = 5)
FLEX_genes <- Flex_probe_ref[Flex_probe_ref$included == TRUE, 'probe_id']
FLEX_genes <- unique(unlist(lapply(str_split(FLEX_genes, '\\|'), '[[', 2)))


##############################################################################################################################
##############################################################################################################################
##############################################################################################################################
# load SCN function
source('/share/ScratchGeneral/tonwan/projects/snPATHO-Seq/scripts/function_scripts/05_SCN_annotation.R')


# run SCN using FLEX data only
working_out <- paste0(out_dir, '/', 'FLEX')
dir.create(working_out, recursive = T)

flex_data_subset <- subset(training_obj, subset = sample_id %in% c("FLEX_exp_1", "FLEX_exp_2"))

run_SCN(training_data = flex_data_subset,
    query_data = query_obj,
    out_dir = working_out,
    annotation_col = 'new_annotation',
    common_genes = FLEX_genes,
    seed = 100,
    output_label = 'FLEX')


# plot confusion matrix

# plot confusion matrix - FLEX data
predicted_metadata <- read.csv(file = paste0(working_out, '/', 'FLEX', "_meta_data_table.csv"), row.names = 1)

plot_df <- as.data.frame(table(predicted_metadata$celltype.l2, predicted_metadata$category))
plot_df <- plot_df %>% 
    group_by(Var1) %>% 
    mutate(ground_truth_sum = sum(Freq)) %>% 
    ungroup() %>% 
    mutate(percent_of_ground_truth = Freq / ground_truth_sum)

# order the plot
plot_df$Var1 <- factor(plot_df$Var1, levels = c('Plasmablast', 'B memory', 'B intermediate', 'B naive',
    'CD8 Naive', 'CD8 TCM', 'CD8 TEM', 'CD8 Proliferating',
    'CD4 Naive', 'CD4 TCM', 'CD4 TEM', 'CD4 CTL', 'CD4 Proliferating', 'Treg',
    'MAIT', 'gdT', 'dnT',
    'NK', 'NK_CD56bright', 'NK Proliferating',
    'CD14 Mono', 'CD16 Mono', 'cDC1', 'cDC2', 'pDC', 'ASDC',
    'HSPC', 'ILC', 'Platelet', 'Eryth', 'Doublet'))
       
plot_df$Var2 <- factor(plot_df$Var2, levels = c('B_memory', 'B_intermediate', 'B_naive',
    'CD8_naive', 'CD8_TCM', 'CD8_TEM',
    'CD4_naive', 'CD4_TCM', 'CD4_TEM', 'CD4_CTL', 'Treg',
    'MAIT', 'CTL_CCR7_high', 
    'NK', 'NK_CD56_bright', 
    'CD14_monocyte', 'CD16_monocyte', 'cDC', 'pDC',
    'HSPC', 'ILC', 'platelet', 'doublet', 'rand'))
                   


plot <- ggplot(plot_df, aes(x = Var1, y = Var2, fill = percent_of_ground_truth, label = Freq)) +
    geom_tile() +
    coord_fixed(ratio = length(unique(plot_df$Var2)) / length(unique(plot_df$Var1))) +
    scale_fill_distiller(palette="Blues", direction=1) +
    labs(x = 'Ground truth labels',
        y = 'Predicted labels',
        fill = "Proportion",
        title = 'Prediction of Hao_et_al_2021 with FLEX signatures') +
    geom_text(color="black", size = 2) +
    theme(axis.text.x = element_text(angle = 45, hjust=1, size = 16),
            axis.text.y = element_text(hjust=1, size = 16),
            legend.text = element_text(size = 20),
            title = element_text(size = 24))

ggsave(file = paste0(working_out, '/', 'FLEX', '_confusion_matrix.pdf'),
    width = 16,
    height = 10,
    plot,
    device = 'pdf')


#############################
#############################
# run SCN using 3p data only ----
working_out <- paste0(out_dir, '/', 'GEX')
dir.create(working_out, recursive = T)

GEX_data_subset <- subset(training_obj, subset = sample_id %in% c("3p_exp_1", "3p_exp_2"))

common_genes <- intersect(rownames(GEX_data_subset), rownames(query_obj))

run_SCN(training_data = GEX_data_subset,
    query_data = query_obj,
    out_dir = working_out,
    annotation_col = 'new_annotation',
    common_genes = common_genes,
    seed = 100,
    output_label = 'GEX')


# plot confusion matrix - GEX data
predicted_metadata <- read.csv(file = paste0(working_out, '/', 'GEX', "_meta_data_table.csv"), row.names = 1)

plot_df <- as.data.frame(table(predicted_metadata$celltype.l2, predicted_metadata$category))
plot_df <- plot_df %>% 
    group_by(Var1) %>% 
    mutate(ground_truth_sum = sum(Freq)) %>% 
    ungroup() %>% 
    mutate(percent_of_ground_truth = Freq / ground_truth_sum)

# order the plot
plot_df$Var1 <- factor(plot_df$Var1, levels = c('Plasmablast', 'B memory', 'B intermediate', 'B naive',
    'CD8 Naive', 'CD8 TCM', 'CD8 TEM', 'CD8 Proliferating',
    'CD4 Naive', 'CD4 TCM', 'CD4 TEM', 'CD4 CTL', 'CD4 Proliferating', 'Treg',
    'MAIT', 'gdT', 'dnT',
    'NK', 'NK_CD56bright', 'NK Proliferating',
    'CD14 Mono', 'CD16 Mono', 'cDC1', 'cDC2', 'pDC', 'ASDC',
    'HSPC', 'ILC', 'Platelet', 'Eryth', 'Doublet'))
       
plot_df$Var2 <- factor(plot_df$Var2, levels = c('B_memory', 'B_intermediate', 'B_naive',
    'CD8_naive', 'CD8_TCM', 'CD8_TEM',
    'CD4_naive', 'CD4_TCM', 'CD4_TEM', 'CD4_CTL', 'Treg',
    'MAIT', 'CTL_CCR7_high', 
    'NK', 'NK_CD56_bright', 
    'CD14_monocyte', 'CD16_monocyte', 'cDC', 'pDC',
    'HSPC', 'platelet', 'doublet', 'rand'))
                   


plot <- ggplot(plot_df, aes(x = Var1, y = Var2, fill = percent_of_ground_truth, label = Freq)) +
    geom_tile() +
    coord_fixed(ratio = length(unique(plot_df$Var2)) / length(unique(plot_df$Var1))) +
    scale_fill_distiller(palette="Blues", direction=1) +
    labs(x = 'Ground truth labels',
        y = 'Predicted labels',
        fill = "Proportion",
        title = 'Prediction of Hao_et_al_2021 with 3p signatures') +
    geom_text(color="black", size = 2) +
    theme(axis.text.x = element_text(angle = 45, hjust=1, size = 16),
            axis.text.y = element_text(hjust=1, size = 16),
            legend.text = element_text(size = 20),
            title = element_text(size = 24))

ggsave(file = paste0(working_out, '/', 'GEX', '_confusion_matrix.pdf'),
    width = 16,
    height = 10,
    plot,
    device = 'pdf')
```
