## Supplementary material for "snPATHO-seq: unlocking the pathology archives": scripts: 01_Run_NMF.R

```
# aim ----
    # run NMF to generate gene modules
    # merge similar modules

.libPaths(
    c(
        '/share/ScratchGeneral/tonwan/projects/snPATHO-Seq/renv/library/R-4.2/x86_64-conda-linux-gnu',
        .libPaths()
    )
)

library(Seurat)
library(tidyverse)
library(NMF)

library(bigmemory)


# arguments ----
# obj_file <- '/share/ScratchGeneral/tonwan/projects/snPATHO-Seq/data/snPATHO_seq/25k_downsampled_processed_objs/integrated/4066_cellbender_filtered_integration_final_annotation.rds'
obj_file <- '/share/ScratchGeneral/tonwan/projects/snPATHO-Seq/data/snPATHO_seq/25k_downsampled_processed_objs/integrated/4399_cellbender_filtered_integration_final_annotation.rds'
# obj_file <- '/share/ScratchGeneral/tonwan/projects/snPATHO-Seq/data/snPATHO_seq/25k_downsampled_processed_objs/integrated/4411_cellbender_filtered_integration_final_annotation.rds'

id <- '4399SNAPFix'
# 4066FFPE 4066GEX 4066SNAPFix
# 4399FFPE_run1 4399FFPE_run2 4399GEX 4399SNAPFix
# 4411FFPE 4411GEX 4411SNAPFix

out_dir <- '/share/ScratchGeneral/tonwan/projects/snPATHO-Seq/results/gene_module_analysis/4399_snap_new_dim_reduction'
working_out <- paste0(out_dir, '/', id)
dir.create(working_out, recursive = T)


# load data ----
obj <- readRDS(file = obj_file)


# run NMF on cancer gene matrix ----
subset_obj <- subset(obj, subset = sample_id == id & 
    major_annotation == 'Epithelial_cancer')
mtx <- GetAssayData(subset_obj, assay = 'RNA', slot = 'data')

# only keep top 7000 genes in the dataset
genes_to_keep <- names(rowSums(mtx)[order(rowSums(mtx), decreasing = TRUE)][1:7000])
mtx_subset <- mtx[genes_to_keep, ]

# make sure there's no gene with only 0 counts
mtx_subset <- mtx_subset[rowSums(mtx_subset) != 0,]

# scale data gene expression matrix
centered_mtx <- apply(mtx_subset, 1, scale)
centered_mtx[centered_mtx < 0] <- 0
centered_mtx[is.na(centered_mtx)] <- 0
rownames(centered_mtx) <- colnames(mtx)
centered_mtx <- t(centered_mtx)


# run NMF
NMF_res <- nmf(x = centered_mtx, rank = 4:9, method="snmf/r", nrun = 10, .opt='v3P')
# save NMF results
saveRDS(file = paste0(working_out, '/', 'NMF_results.rds'), NMF_res)
```
