## Supplementary material for "snPATHO-seq: unlocking the pathology archives": scripts: 02_format_raw_NMF_results.R

```
# aim ----
    # format raw NMF results and name them in the same way as Itay's team does


# define environment ----
library(Seurat)
library(tidyverse)
library(NMF)


# arguments ----
NMF_res_dir <- '/share/ScratchGeneral/tonwan/projects/snPATHO-Seq/results/gene_module_analysis'

samples <- dir(NMF_res_dir)[dir(NMF_res_dir) != 'archive']

cell_type <- 'Epithelial_cancer'


# make a list containing all NMF programs defined ----
merged_NMF_list <- list()

for (s in samples) {
    # load NMF results
    NMF_result_file <- paste0(NMF_res_dir, '/', s, '/', 'NMF_results.rds')
    NMF_results <- readRDS(file = NMF_result_file)

    # create a uniformed name for the NMF analysis result
    entry_name <- paste0(s, '_', cell_type, '_rank4_9_nruns10.RDS')

    # merge all programs defined within each sample
    working_list <- list()

    for (n in names(NMF_results$fit)) {
        working_res <- paste('NMF_results$fit$', n, '@fit@W', sep = "'")
        working_matrix <- eval(parse(text = working_res))

        module_index <- seq(1:ncol(working_matrix))
        module_names <- paste0(entry_name, '.', n, '.', module_index)
        colnames(working_matrix) <- module_names
        working_list[[n]] <- working_matrix
    }
    merged_NMF_mtx <- Reduce(cbind, working_list)

    # add merged NMF matrix to a list
    merged_NMF_list[[entry_name]] <- merged_NMF_mtx
}

# save merged list
saveRDS(file = paste0(NMF_res_dir, '/', 'merged_', cell_type, "_module_list.rds"), merged_NMF_list)


# # NMF_result_file <- '/share/ScratchGeneral/tonwan/projects/snPATHO-Seq/results/gene_module_analysis/4066FFPE/NMF_results.rds'

# # sample_id <- '4066FFPE'


# # load NMF results ----
# NMF_results <- readRDS(file = NMF_result_file)


# # name and merge NMF programs from different parameters together
# merged_NMF_list <- list()

# for (n in names(NMF_results$fit)) {
#     working_res <- paste('NMF_results$fit$', n, '@fit@W', sep = "'")
#     working_matrix <- eval(parse(text = working_res))

#     module_index <- seq(1:ncol(working_matrix))
#     module_names <- paste0(entry_name, '.', n, '.', module_index)
#     colnames(working_matrix) <- module_names
#     merged_NMF_list[[n]] <- working_matrix
# }

# merged_NMF_mtx <- Reduce(cbind, merged_NMF_list)
```
