## Supplementary material for "snPATHO-seq: unlocking the pathology archives": scripts: 03_get_robust_NMF.R

```
# aim ----
    # define robust NMF programs from the results


# define environment ----
library(Seurat)
library(tidyverse)


# arguments ----
NMF_list_file <- '/share/ScratchGeneral/tonwan/projects/snPATHO-Seq/results/gene_module_analysis/merged_Epithelial_cancer_module_list.rds'

out_dir <- '/share/ScratchGeneral/tonwan/projects/snPATHO-Seq/results/gene_module_analysis'

Min_intersect_initial <- 10    # the minimal intersection cutoff for defining the first NMF program in a cluster
Min_intersect_cluster <- 5    # the minimal intersection cutoff for adding a new NMF to the forming cluster 
Min_group_size        <- 3     # the minimal group size to consider for defining the first NMF program in a cluster 


####################################################################################################################################
####################################################################################################################################
####################################################################################################################################
# load NMF results
NMF_list <- readRDS(file = NMF_list_file)
# only keep top 50 genes of each program
nmf_programs <- lapply(NMF_list, function(x) apply(x, 2, function(y) names(sort(y, decreasing = T))[1:50]))


# select robust NMF within each sample ----
# load predefined function
source('/share/ScratchGeneral/tonwan/projects/snPATHO-Seq/scripts/function_scripts/Itay_2023_gene_module_paper/robust_nmf_programs.R')

robust_NMF_names <- robust_nmf_programs(nmf_programs, intra_min = 35, intra_max = 10, inter_filter=T, inter_min = 10)

filtered_nmf_programs <- lapply(nmf_programs, function(x) x[, is.element(colnames(x), robust_NMF_names),drop=F])
filtered_nmf_programs <- do.call(cbind, filtered_nmf_programs)

# save robust NMF programs
saveRDS(file = paste0(out_dir, '/', 'robust_NMF_programs.rds'), filtered_nmf_programs)
```
