## Supplementary material for "snPATHO-seq: unlocking the pathology archives": scripts: 01_run_NMF.R

```
# aim ----
    # run NMF to generate gene modules
    # merge similar modules

.libPaths(
    c(
        '/share/ScratchGeneral/tonwan/projects/snPATHO-Seq/renv/library/R-4.2/x86_64-conda-linux-gnu',
        .libPaths()
    )
)

library(Seurat)
library(tidyverse)
library(NMF)

library(bigmemory)


# arguments ----
dataset_label <- '4066FFPE'
# 4399FFPE 4066FFPE

obj_file <- paste0('/share/ScratchGeneral/tonwan/projects/snPATHO-Seq/data/Visium/processed/STutility/', dataset_label, '.rds')

cnv_res_dir <- paste0('/share/ScratchGeneral/tonwan/projects/snPATHO-Seq/results/visium_CNV/', dataset_label)

out_dir <- paste0('/share/ScratchGeneral/tonwan/projects/snPATHO-Seq/results/visium_gene_module_analysis/', dataset_label)
dir.create(out_dir, recursive = T)


# load data ----
obj <- readRDS(file = obj_file)
# make sure the data is normalised
obj <- NormalizeData(obj)

# load CNV prediction results
cnv_spots_df <- read.csv(file = paste0(cnv_res_dir, '/', 'dendrogram_plots/spots_with_CNV_changes.csv'), row.names = 1)
if (is.null(cnv_spots_df$label)) {
    cnv_spots_df$label <- 'population_1'
}

# split data by CNV clones and only run NMF on cancer populations
for (p in unique(cnv_spots_df$label)) {
    working_spots <- cnv_spots_df[cnv_spots_df$label == p, 'sub_phylo_id']
    subset_obj <- subset(obj, cells = working_spots)

    # get data matrix
    mtx <- GetAssayData(subset_obj, assay = 'Spatial', slot = 'data')

    # only keep top 7000 genes in the dataset
    genes_to_keep <- names(rowSums(mtx)[order(rowSums(mtx), decreasing = TRUE)][1:7000])
    mtx_subset <- mtx[genes_to_keep, ]

    # make sure there's no gene with only 0 counts
    mtx_subset <- mtx_subset[rowSums(mtx_subset) != 0,]

    # scale data gene expression matrix
    centered_mtx <- apply(mtx_subset, 1, scale)
    centered_mtx[centered_mtx < 0] <- 0
    centered_mtx[is.na(centered_mtx)] <- 0
    rownames(centered_mtx) <- colnames(mtx)
    centered_mtx <- t(centered_mtx)

    # run NMF
    NMF_res <- nmf(x = centered_mtx, rank = 4:9, method="snmf/r", nrun = 10, .opt='v3P')
    # save NMF results
    saveRDS(file = paste0(out_dir, '/', 'NMF_results.rds'), NMF_res)

}
```
