## Supplementary material for "snPATHO-seq: unlocking the pathology archives": scripts: 03_heatmap_overview_with_Gavish.R

```
# aim ----
    # check overlapping of gene modules derived from the current dataset with published gene modules


# define environment ----
library(tidyverse)
library(readxl)

library(viridis)
library(ComplexHeatmap)
library(circlize)

library(ape)
library(phylogram)


# arguments ----
snRNA_nmf_file <- '/share/ScratchGeneral/tonwan/projects/snPATHO-Seq/results/gene_module_analysis/robust_NMF_programs.rds'
visium_nmf_file <- '/share/ScratchGeneral/tonwan/projects/snPATHO-Seq/results/visium_gene_module_analysis/robust_NMF_programs.rds'

Gavish_meta_program_file <- '/share/ScratchGeneral/tonwan/projects/snPATHO-Seq/resources/meta_programs.xlsx'

out_dir <- '/share/ScratchGeneral/tonwan/projects/snPATHO-Seq/results/gene_module_analysis/snRNA_visium_together'
dir.create(out_dir, recursive = T)


# load data ----
snRNA_nmf <- readRDS(file = snRNA_nmf_file)
snRNA_nmf <- as.data.frame(snRNA_nmf)

visium_nmf <- readRDS(file = visium_nmf_file)
visium_nmf <- as.data.frame(visium_nmf)

Gavish_meta_programs <- read_excel(Gavish_meta_program_file)
Gavish_meta_programs <- as.data.frame(Gavish_meta_programs)


all_program_df <- cbind(Gavish_meta_programs, snRNA_nmf, visium_nmf)


########################################################################################################################
########################################################################################################################
########################################################################################################################
# check numbers of overlapping genes between robust NMF programs
nmf_intersect <- apply(all_program_df , 2, function(x) apply(all_program_df , 2, function(y){
    return(length(intersect(x,y)))
})) 


# hierchal clustering of the programs
nmf_intersect_hc <- hclust(as.dist(50-nmf_intersect), method="average") 


# plot dendrogram ----
phylo_tree <- as.phylo(as.dendrogram(nmf_intersect_hc))

# plot dendrogram
pdf(file = paste0(out_dir, '/', 'Phylo_tree_FLEX_gene_filtered.pdf'),
    height = 25, width = 15)
    plot(phylo_tree, show.tip.label = TRUE, show.node.label = FALSE)
    tiplabels() 
dev.off() 

# export the order of the heatmap plot
write.csv(file = paste0(out_dir, '/', 'heatmap_GEX_FLEX_common_genes_plot_order.csv'), as.data.frame(phylo_tree$tip.label))


# prepare heatmap annotation ----
annotation_df <- data.frame(programs = colnames(nmf_intersect))
annotation_df$sample <- ifelse(annotation_df$programs %in% grep('4066', annotation_df$programs, value = T), '4066',
    ifelse(annotation_df$programs %in% grep('4399', annotation_df$programs, value = T), '4399',
    ifelse(annotation_df$programs %in% grep('4411', annotation_df$programs, value = T), '4411', 'Gavish_et_al_2023')))

annotation_df$processing_method <- ifelse(annotation_df$programs %in% grep('FFPE', annotation_df$programs, value = T), 'snPATHO',
    ifelse(annotation_df$programs %in% grep('SNAP', annotation_df$programs, value = T), 'FLEX',
    ifelse(annotation_df$programs %in% grep('GEX', annotation_df$programs, value = T), '3p', 'Gavish_et_al_2023')))

annotation_df$processing_method <- ifelse(annotation_df$programs %in% grep('spatial', annotation_df$programs, value = T), 'Spatial', annotation_df$processing_method)


# plot heatmap ----
col_fun <- colorRamp2(c(0, 5, 10, 15, 20, 25), 
                      c('white', rev(inferno(n = 6))[2:length(rev(inferno(n = 6)))]))


# load preset colors for snRNA-seq workflows
source('/share/ScratchGeneral/tonwan/projects/snPATHO-Seq/scripts/colors.R')
workflow_colors <- workflow_colors[unique(annotation_df$processing_method)]
sample_colors <- sample_colors[unique(annotation_df$sample)]


# make annotation
top_annot <- HeatmapAnnotation(
    Samples = annotation_df$sample,
    Workflows = annotation_df$processing_method,
    which = 'column',
    col = list(Samples = sample_colors,
                Workflows = workflow_colors), 
    annotation_name_gp = grid::gpar(fontsize = 16),
    annotation_legend_param = list(
        Samples = list(title_gp = gpar(fontsize = 16, fontface = "bold"), 
                                    labels_gp = gpar(fontsize = 16)),
        Workflows = list(title_gp = gpar(fontsize = 16, fontface = "bold"), 
                                    labels_gp = gpar(fontsize = 16))))


plot <- Heatmap(nmf_intersect,
                name = 'Overlapping',
                heatmap_legend_param = list(title_gp = gpar(fontsize = 16, fontface = "bold")),
                col = col_fun,
                top_annotation = top_annot,
                show_column_names= FALSE,
                show_row_names = FALSE,
                cluster_rows = as.dendrogram(nmf_intersect_hc),
                cluster_columns = as.dendrogram(nmf_intersect_hc),
                column_names_gp = gpar(fontsize = 14),
                row_names_gp = gpar(fontsize = 14))

pdf(file = paste0(out_dir, '/', 'Gene_module_overlapping_heatmap', '.pdf'),
    width = 12, height = 10)
    draw(plot, merge_legend = TRUE)
dev.off()
```
