## Supplementary material for "snPATHO-seq: unlocking the pathology archives": scripts: 04_summarise_core_genes_from_robust_NMF.R

```
# aim ----
    # after clustering with Gavish_et_al_2023 meta programs, get core genes from different clusters
        # due to small sample size in the current study, any gene present in more than 2 robust NMF programs within the same cluster is considered as a core gene


# define environment ----
library(tidyverse)
library(readxl)


# arguments ----
snRNA_nmf_file <- '/share/ScratchGeneral/tonwan/projects/snPATHO-Seq/results/gene_module_analysis/robust_NMF_programs.rds'
spatial_nmf_file <- '/share/ScratchGeneral/tonwan/projects/snPATHO-Seq/results/visium_gene_module_analysis/robust_NMF_programs.rds'

out_dir <- '/share/ScratchGeneral/tonwan/projects/snPATHO-Seq/results/gene_module_analysis/core_genes'
dir.create(out_dir, recursive = T)


# load data ----
snRNA_nmf <- readRDS(file = snRNA_nmf_file)
snRNA_nmf <- as.data.frame(snRNA_nmf)

spatial_nmf <- readRDS(file = spatial_nmf_file)
spatial_nmf <- as.data.frame(spatial_nmf)

all_nmf <- cbind(snRNA_nmf, spatial_nmf)


nmf_cell_cycle <- c(
    '4066GEX_Epithelial_cancer_rank4_9_nruns10.RDS.4.2',
    '4411GEX_Epithelial_cancer_rank4_9_nruns10.RDS.5.2',
    '4399FFPE_run1_Epithelial_cancer_rank4_9_nruns10.RDS.4.3',
    '4399SNAPFix_Epithelial_cancer_rank4_9_nruns10.RDS.6.6',
    '4066FFPE_Epithelial_cancer_rank4_9_nruns10.RDS.4.1',
    '4066SNAPFix_Epithelial_cancer_rank4_9_nruns10.RDS.5.3',
    '4411FFPE_Epithelial_cancer_rank4_9_nruns10.RDS.5.3',
    '4411SNAPFix_Epithelial_cancer_rank4_9_nruns10.RDS.5.5',
    '4066FFPE_Epithelial_cancer_spatial_rank4_9_nruns10.RDS.8.8',
    '4399FFPE_Epithelial_cancer_spatial_rank4_9_nruns10.RDS.6.5'
    )

nmf_4399 <- c(
    '4399GEX_Epithelial_cancer_rank4_9_nruns10.RDS.9.8',
    '4399FFPE_run1_Epithelial_cancer_rank4_9_nruns10.RDS.9.5',
    '4399SNAPFix_Epithelial_cancer_rank4_9_nruns10.RDS.7.2'
    )

nmf_4411_1 <- c('4411GEX_Epithelial_cancer_rank4_9_nruns10.RDS.7.5',
    '4411FFPE_Epithelial_cancer_rank4_9_nruns10.RDS.7.6',
    '4411SNAPFix_Epithelial_cancer_rank4_9_nruns10.RDS.7.2')

nmf_4066 <- c('4066GEX_Epithelial_cancer_rank4_9_nruns10.RDS.8.7',
    '4066FFPE_Epithelial_cancer_rank4_9_nruns10.RDS.6.1',
    '4066SNAPFix_Epithelial_cancer_rank4_9_nruns10.RDS.6.1')

nmf_mixed_1 <- c(
    '4066GEX_Epithelial_cancer_rank4_9_nruns10.RDS.8.8',
    '4399FFPE_run1_Epithelial_cancer_rank4_9_nruns10.RDS.5.1',
    '4066FFPE_Epithelial_cancer_rank4_9_nruns10.RDS.9.1',
    '4066SNAPFix_Epithelial_cancer_rank4_9_nruns10.RDS.8.1',
    '4411FFPE_Epithelial_cancer_rank4_9_nruns10.RDS.9.1',
    '4411SNAPFix_Epithelial_cancer_rank4_9_nruns10.RDS.8.6'
)

nmf_4411_2 <- c('4411GEX_Epithelial_cancer_rank4_9_nruns10.RDS.7.6',
    '4411FFPE_Epithelial_cancer_rank4_9_nruns10.RDS.6.2',
    '4411SNAPFix_Epithelial_cancer_rank4_9_nruns10.RDS.7.5')

nmf_4411_3 <- c('4411GEX_Epithelial_cancer_rank4_9_nruns10.RDS.6.2',
    '4411FFPE_Epithelial_cancer_rank4_9_nruns10.RDS.7.3',
    '4411SNAPFix_Epithelial_cancer_rank4_9_nruns10.RDS.7.3')

nmf_stress <- c('4399GEX_Epithelial_cancer_rank4_9_nruns10.RDS.7.2',
    '4399FFPE_run1_Epithelial_cancer_rank4_9_nruns10.RDS.7.5',
    '4399SNAPFix_Epithelial_cancer_rank4_9_nruns10.RDS.8.8')

nmf_hypoxia <- c('4399GEX_Epithelial_cancer_rank4_9_nruns10.RDS.9.3',
    '4399FFPE_run1_Epithelial_cancer_rank4_9_nruns10.RDS.6.4',
    '4399SNAPFix_Epithelial_cancer_rank4_9_nruns10.RDS.9.3',
    '4411GEX_Epithelial_cancer_rank4_9_nruns10.RDS.6.1',
    '4411FFPE_Epithelial_cancer_rank4_9_nruns10.RDS.7.4',
    '4411SNAPFix_Epithelial_cancer_rank4_9_nruns10.RDS.7.7',
    '4066GEX_Epithelial_cancer_rank4_9_nruns10.RDS.6.6',
    '4066FFPE_Epithelial_cancer_rank4_9_nruns10.RDS.7.5',
    '4066SNAPFix_Epithelial_cancer_rank4_9_nruns10.RDS.7.2',
    '4066FFPE_Epithelial_cancer_spatial_rank4_9_nruns10.RDS.7.5',
    '4399FFPE_Epithelial_cancer_spatial_rank4_9_nruns10.RDS.7.7',
    '4411FFPE_Epithelial_cancer_spatial_rank4_9_nruns10.RDS.5.4')


nmf_mixed_2 <- c(
    '4399FFPE_Epithelial_cancer_spatial_rank4_9_nruns10.RDS.7.6',
    '4411FFPE_Epithelial_cancer_spatial_rank4_9_nruns10.RDS.5.1',
    '4411FFPE_Epithelial_cancer_rank4_9_nruns10.RDS.7.7',
    '4411SNAPFix_Epithelial_cancer_rank4_9_nruns10.RDS.6.4'
)

nmf_mixed_3 <- c(
    '4411FFPE_Epithelial_cancer_rank4_9_nruns10.RDS.8.8',
    '4411SNAPFix_Epithelial_cancer_rank4_9_nruns10.RDS.7.1',
    '4399GEX_Epithelial_cancer_rank4_9_nruns10.RDS.4.3',
    '4411GEX_Epithelial_cancer_rank4_9_nruns10.RDS.8.2',
    '4066FFPE_Epithelial_cancer_rank4_9_nruns10.RDS.4.4',
    '4066SNAPFix_Epithelial_cancer_rank4_9_nruns10.RDS.5.5',
    '4399FFPE_run1_Epithelial_cancer_rank4_9_nruns10.RDS.4.4',
    '4399SNAPFix_Epithelial_cancer_rank4_9_nruns10.RDS.4.3'
)

nmf_EMT <- c(
    '4066FFPE_Epithelial_cancer_spatial_rank4_9_nruns10.RDS.8.3',
    '4399GEX_Epithelial_cancer_rank4_9_nruns10.RDS.7.6',
    '4066FFPE_Epithelial_cancer_spatial_rank4_9_nruns10.RDS.4.3',
    '4399FFPE_Epithelial_cancer_spatial_rank4_9_nruns10.RDS.4.3'
)


all_clusters <- c("nmf_4066", "nmf_4399", "nmf_4411_1", "nmf_4411_2", "nmf_4411_3", "nmf_cell_cycle",     
    "nmf_EMT", "nmf_hypoxia", "nmf_mixed_1", 'nmf_mixed_2', 'nmf_mixed_3', 'nmf_stress')


for (c in all_clusters) {
    working_nmf <- all_nmf[, eval(parse(text = c))]
    working_nmf_long <- gather(working_nmf, key = 'module_names', value = 'genes')
    detection_freq <- table(working_nmf_long$genes)
    core_genes <- names(detection_freq[detection_freq >= 2])
    write.csv(file = paste0(out_dir, '/', c, '.csv'), core_genes)
}
```
