## Supplementary material for "snPATHO-seq: unlocking the pathology archives": scripts: 05_Map_GM_on_UMAP.R

```
# aim ----
    # score GM on all samples
    # plot GM scores


# define environment ----
library(Seurat)

library(RColorBrewer)
library(clusterProfiler)
library(tidyverse)


# arguments ----
snRNA_obj_dir <- '/share/ScratchGeneral/tonwan/projects/snPATHO-Seq/data/snPATHO_seq/25k_downsampled_processed_objs/integrated'

nmf_gene_list_dir <- '/share/ScratchGeneral/tonwan/projects/snPATHO-Seq/results/gene_module_analysis/core_genes'

samples <- c('4066', '4399', '4411')


out_dir <- '/share/ScratchGeneral/tonwan/projects/snPATHO-Seq/results/gene_module_analysis/map_GM'
dir.create(out_dir, recursive = T)


# load object ----
obj_list <- list()
for (s in samples) {
    obj <- readRDS(file = paste0(snRNA_obj_dir, '/', s, '_cellbender_filtered_integration_final_annotation.rds'))
    obj_list[[s]] <- obj
}


# load NMF gene lists ----
nmf_gene_list <- list()
for (m in list.files(nmf_gene_list_dir, '.csv')) {
    working_nmf <- read.csv(file = paste0(nmf_gene_list_dir, '/', m), row.names = 1)
    output_name <- unlist(lapply(str_split(m, '\\.csv'), '[[', 1))
    nmf_gene_list[[output_name]] <- working_nmf[[1]]
}


# load genes from Gavish_et_al_2023 meta programs
Gavish_G2M <- c('TOP2A','UBE2C','HMGB2','NUSAP1','CENPF','CCNB1','TPX2','CKS2','BIRC5','PRC1','PTTG1','KPNA2','MKI67','CDC20','CDK1','CCNB2','CDKN3','SMC4','NUF2','ARL6IP1','CKAP2','ASPM','PLK1','CKS1B','CCNA2','AURKA','MAD2L1','GTSE1','HMMR','UBE2T','CENPE','CENPA','KIF20B','AURKB','CDCA3','CDCA8','UBE2S','KNSTRN','KIF2C','PBK','TUBA1B','DLGAP5','TACC3','STMN1','DEPDC1','ECT2','CENPW','ZWINT','HIST1H4C','KIF23')
Gavish_G1S <- c('PCNA','RRM2','FEN1','GINS2','TYMS','MCM3','GMNN','HIST1H4C','CLSPN','ATAD2','TK1','KIAA0101','DUT','HELLS','MCM7','UBE2T','MCM4','CENPU','DHFR','ZWINT','ASF1B','MCM5','DNAJC9','RFC4','HMGB2','CDC6','RRM1','ORC6','CDK1','RAD51AP1','RNASEH2A','CHAF1A','CENPK','CDCA5','SLBP','MCM6','TMEM106C','CENPM','MYBL2','E2F1','USP1','DNMT1','PKMYT1','MAD2L1','PSMC3IP','CDCA4','RFC2','CDC45','UHRF1','MCM2')
Gavish_hypoxia <- c('NDRG1','BNIP3','SLC2A1','P4HA1','VEGFA','ENO2','ADM','HILPDA','PGK1','BNIP3L','FAM162A','PLOD2','EGLN3','LDHA','MT1X','NRN1','DDIT4','DDIT3','ERO1A','ANGPTL4','IGFBP3','PFKP','INSIG2','NDUFA4L2','PDK1','CA9','AKAP12','C4ORF3','IGFBP5','GBE1','LGALS3','TMEM45A','WSB1','GPI','S100A10','SLC16A3','EPB41L4A-AS1','ALDOA','ANKRD37','CAV1','AK4','DNAJB9','ERRFI1','MT2A','SLC3A2','VIM','ENO1','PFKFB3','HK2','CA12')
Gavish_stress <- c('ATF3','EGR1','FOS','PPP1R15A','JUN','DUSP1','JUNB','FOSB','IER2','GADD45B','BTG2','ZFP36','DNAJB1','NR4A1','SERTAD1','DDIT3','KLF6','NFKBIA','HSPA1A','HSPA1B','CYR61','MAFF','NFKBIZ','IER3','CCNL1','MCL1','RHOB','SOCS3','IRF1','CDKN1A','HES1','ID2','KLF4','KLF10','CXCL2','ID1','TOB1','TRIB1','HSPH1','RASD1','SAT1','DDIT4','EGR2','HERPUD1','PLK2','DNAJA1','HSPA6','PMAIP1','CITED2','ID3')
Gavish_stress_invitro <- c('DDIT3','SLC3A2','TRIB3','ASNS','HERPUD1','PPP1R15A','EPB41L4A-AS1','SNHG12','GARS','GADD45A','MTHFD2','XBP1','CEBPG','TAF1D','NUPR1','SARS','SNHG8','ZFAS1','DDIT4','PHGDH','SHMT2','TXNIP','ATF4','HSPA5','GDF15','CARS','MAP1LC3B','SQSTM1','YARS','DNAJB9','PSAT1','GADD45B','ATF3','SNHG15','RSRC2','CEBPB','ARF4','EIF1B','OSER1','BTG1','SNHG7','WARS','ZFAND2A','C6ORF48','STC2','ZNF622','HSPA9','PDRG1','UPP1','CCDC174')
Gavish_EMT_1 <- c('COL1A2','SPARC','COL1A1','COL3A1','IGFBP7','TAGLN','C1S','FN1','MGP','TIMP1','TIMP3','BGN','C1R','VIM','COL6A2','COL4A2','COL6A1','MMP2','DCN','COL4A1','FSTL1','NNMT','TPM1','TSC22D3','CCDC80','CTGF','CXCL12','ID3','LUM','MYL9','PRSS23','TGFBI','TPM2','CAV1','CYR61','IGFBP5','LGALS1','SPARCL1','A2M','ACTA2','CALD1','COL6A3','CTSK','FBLN1','IGFBP4','POSTN','RARRES2','SERPING1','SNAI2','TCF4')
Gavish_EMT_2 <- c('LAMC2','LAMB3','LAMA3','EMP3','TGFBI','TNFRSF12A','FLNA','PLAU','PMEPA1','CAV1','KRT17','PRSS23','LGALS1','VIM','S100A2','COL17A1','FSTL3','CST6','IL32','PDLIM7','SERPINE1','ANXA3','IGFBP7','DCBLD2','F3','MT2A','TNC','ITGB6','KRT7','PLAUR','PRKCDBP','SERPINE2','ITGB1','RBP1','ITGA3','CDKN1A','THBS1','FN1','MMP7','TGM2','C15ORF48','ITGA6','PTHLH','TPM1','CDA','IGFBP6','PDPN','RHOD','INHBA','ITGAV')
Gavish_EMT_3 <- c('ANXA2','KRT19','S100A6','TNFRSF12A','KRT7','LGALS1','ANXA1','TM4SF1','S100A10','LGALS3','S100A16','TAGLN2','LMNA','CRIP1','EZR','KRT18','S100A11','S100A4','EMP3','HSPB1','KRT8','TUBA1A','VIM','IL32','KLF6','ANXA3','SH3BGRL3','TMSB4X','C19ORF33','CKB','CLIC1','EMP1','IFI27','ISG15','MDK','MMP7','PHLDA2','PLP2','CLDN4','MYADM','RHOC','S100A14','CD24','CRIP2','MYL12A','RAB11FIP1','S100P','STMN1','FXYD3','FLNA')
Gavish_EMT_4 <- c('KRT15','CXCL14','ALDH3A1','DAPL1','DST','CCL2','TXNIP','C12ORF75','COL17A1','DCN','HOPX','MOXD1','SDCBP','SEPP1','THBS2','TIMP1','ASS1','GLUL','NFIB','NTRK2','SERPINH1','SOX4','ANTXR1','C1R','C1S','C6ORF48','CAV1','FTH1','IGFBP5','NCOA7','NINJ1','PNRC1','SERPINE2','SERPINF1','SLC47A2','SNAI2','SOCS3','SPARC','TSC22D1','ASPN','LTF','SFRP1','TP53AIP1','CD74','EGR1','BCAM','SESN3','IFITM1','HTRA1','IFIT3')

Gavish_list <- list()
for (t in c('Gavish_G2M', 'Gavish_G1S', 'Gavish_hypoxia', 'Gavish_stress', 
    'Gavish_stress_invitro', 'Gavish_EMT_1', 'Gavish_EMT_2', 'Gavish_EMT_3', 'Gavish_EMT_4')) {
    Gavish_list[[t]] <- eval(parse(text = t))
}


# merge all gene lists together
all_gene_list <- c(nmf_gene_list, Gavish_list)


# get module scores
for (o in names(obj_list)) {
    working_obj <- obj_list[[o]]

    working_out <- paste0(out_dir, '/', o, '_seurat_module_scores')
    dir.create(working_out, recursive = T)

    for (n in names(all_gene_list)) {
        annotated_obj <- AddModuleScore(
            object = working_obj,
            features = all_gene_list[n],
            assay = 'RNA',
            name = n)
        annotated_obj@meta.data[[n]] <- annotated_obj@meta.data[[paste0(n, '1')]] 

        pdf(file = paste0(working_out, '/', n, '.pdf'))
            print(
                FeaturePlot(annotated_obj, features = n, order = TRUE) +
                    scale_color_gradient2(low = '#1878b2', mid = '#e5e5e5', high = '#B2182B')
            )
        dev.off()
    } 
}
```
