## Supplementary material for "snPATHO-seq: unlocking the pathology archives": scripts: 4066_cell_assignment_DE.R

```
# aim ----
    # use gene module scores to select cells
        # only keep cancer cells
        # define cells using GM score > 0.5
    


# define environment ----
library(Seurat)
library(tidyverse)

library(RColorBrewer)


# arguments ----
obj_file <- '/share/ScratchGeneral/tonwan/projects/snPATHO-Seq/data/snPATHO_seq/25k_downsampled_processed_objs/integrated/4066_cellbender_filtered_integration_final_annotation.rds'

nmf_gene_list_dir <- '/share/ScratchGeneral/tonwan/projects/snPATHO-Seq/results/gene_module_analysis/core_genes'

out_dir <- '/share/ScratchGeneral/tonwan/projects/snPATHO-Seq/results/gene_module_analysis/4066_unique_GMs'
dir.create(out_dir, recursive = T)

min_score <- 0.5


################################################################################################################
################################################################################################################
################################################################################################################
# get genes from the robust NMF programs
programs <- c('nmf_4066', 'nmf_hypoxia', 'nmf_cell_cycle', 'nmf_stress')

NMF_list <- list()
for (p in programs) {
    genes <- read.csv(file = paste0(nmf_gene_list_dir, '/', p, '.csv'), row.names = 1)
    NMF_list[[p]] <- genes[[1]]
}


# get module scores for each gene modules
obj <- readRDS(file = obj_file)

for (t in names(NMF_list)) {
    obj <- AddModuleScore(
        object = obj,
        features = NMF_list[t],
        assay = 'RNA',
        name = t)
    obj@meta.data[[t]] <- obj@meta.data[[paste0(t, '1')]] 
    obj@meta.data <- obj@meta.data[, !(colnames(obj@meta.data) == paste0(t, '1'))]
}


# check assignmetn ----
# only keep cancer cells 
subset_obj <- subset(obj, subset = major_annotation == 'Epithelial_cancer')


# get maximum score each cell has been assigned
score_columns <- grep('NMF', colnames(subset_obj@meta.data), value = T, ignore.case = T)
max_scores <- apply(subset_obj@meta.data[, score_columns], 1, function(x) max(x))
subset_obj$max_scores <- max_scores
# remove cells with no score over minimal requirement
subset_obj <- subset(subset_obj, subset = max_scores > min_score)
# assign cells based on maximum scores
assignment <- apply(subset_obj@meta.data[, score_columns], 1, function(x){
    x <- as.numeric(x)
    score_columns[which(x == max(x))]
})
subset_obj$assigment <- assignment


# plot to check assignment ----
pdf(file = paste0(out_dir, '/', 'GM_assignment.pdf'), width = 8, height = 7)
    print(
        DimPlot(subset_obj, group.by = 'assigment')
    )
dev.off()


saveRDS(file = paste0(out_dir, '/', '4066_GM_assigned.rds'), subset_obj)


# plot assignment on all cells - all other cells label as unassigned
meta_df <- obj@meta.data
subset_meta_df <- subset_obj@meta.data
merged_meta_df <- base::merge(meta_df, subset_meta_df[, 'assigment', drop = FALSE], by = 0, all.x = T)
merged_meta_df$assigment <- ifelse(is.na(merged_meta_df$assigment), 'no_assignment', merged_meta_df$assigment)

merged_meta_df <- column_to_rownames(merged_meta_df, var = 'Row.names')
merged_meta_df <- merged_meta_df[match(rownames(obj@meta.data), rownames(merged_meta_df)),]
obj@meta.data <- merged_meta_df

colors <- brewer.pal(name = 'Dark2', n = 4)
colors <- c(colors, '#e5e5e5')
names(colors) <- c('nmf_cell_cycle', 'nmf_hypoxia', 'nmf_stress',
    'nmf_4066', 'no_assignment')


pdf(file = paste0(out_dir, '/', 'GM_assignment_paper.pdf'), width = 8, height = 7)
    print(
        DimPlot(obj, group.by = 'assigment', cols = colors, shuffle = TRUE)
    )
dev.off()


################################################################################################################
################################################################################################################
################################################################################################################
# DE between cancer populations 
de_obj <- subset(obj, subset = major_annotation == 'Epithelial_cancer')

Idents(de_obj) <- 'assigment'
de_res <- FindAllMarkers(de_obj, assay = 'RNA', slot = "data", logfc.threshold = 0, only.pos = FALSE, return.thresh = Inf)
saveRDS(file = paste0(out_dir, '/', 'DE_between_GM_cells.rds'), de_res)
```
