## Supplementary material for "snPATHO-seq: unlocking the pathology archives": scripts: 4066_pathway_analysis.R

```
# aim ----
    # check top pathway between cells annotated by gene modules


# define environment ----
library(Seurat)
library(tidyverse)

library(clusterProfiler)
library(org.Hs.eg.db)
library(fgsea)

library(RColorBrewer)


# arguments ----
de_res_file <- '/share/ScratchGeneral/tonwan/projects/snPATHO-Seq/results/gene_module_analysis/4066_unique_GMs/DE_between_GM_cells.rds'

obj_file <- '/share/ScratchGeneral/tonwan/projects/snPATHO-Seq/data/snPATHO_seq/25k_downsampled_processed_objs/integrated/4066_cellbender_filtered_integration_final_annotation.rds'

out_dir <- '/share/ScratchGeneral/tonwan/projects/snPATHO-Seq/results/gene_module_analysis/4066_unique_GMs'
dir.create(out_dir, recursive = T)

reactome_gs <- '/share/ScratchGeneral/tonwan/projects/snPATHO-Seq/resources/pathway_gmt/c2.cp.reactome.v2022.1.Hs.entrez.gmt'


##############################################################################################################################
##############################################################################################################################
##############################################################################################################################
# run GSEA using DE results

# load DE results 
de_res <- readRDS(file = de_res_file)


# convert gene symbols to gene entrez ID 
entrez_ID_df <- bitr(de_res$gene, fromType="SYMBOL", toType="ENTREZID", OrgDb="org.Hs.eg.db")
de_res_entrez <- base::merge(de_res, entrez_ID_df, by.x = 'gene', by.y = 'SYMBOL', all.x = TRUE)

# sort genes by FC
de_res_entrez <- de_res_entrez %>% 
    arrange(desc(avg_log2FC)) %>% 
    filter(!(is.na(ENTREZID)))

# turn results into gene lists
gene_lists <- list()
for (ds in unique(de_res_entrez$cluster)) {
    working_res <- de_res_entrez[de_res_entrez$cluster == ds, ]
    FCs <- working_res$avg_log2FC
    names(FCs) <- working_res$ENTREZID
    gene_lists[[ds]] <- FCs
}


# obtained from 
    # https://bioinformaticsbreakdown.com/how-to-gsea/
GSEA <- function(gmt_file, gene_list) {
    # load reference gmt files
    gs <- fgsea::gmtPathways(gmt_file)
    # run fgsea
    fgRes <- fgsea::fgsea(pathways = gs,
                           stats = gene_list,
                           minSize=15, ## minimum gene set size
                           maxSize=200, ## maximum gene set size
                           nperm=10000) 
    # filter results 
    fgRes_filtered <- fgRes %>% 
        as.data.frame() %>% 
        filter(padj < 0.05) %>% 
        arrange(desc(NES))

    # collapse similar pathways
    concise_pathways <- collapsePathways(data.table::as.data.table(fgRes_filtered),
                                      pathways = gs,
                                      stats = gene_list)  
    fgRes_filtered <- fgRes_filtered[fgRes_filtered$pathway %in% concise_pathways$mainPathways, ]

    # return results 
    return(fgRes_filtered)
}


# run GSEA analysis
reactome_res_list <- lapply(gene_lists, function(x){
    GSEA(gmt_file = reactome_gs,
        gene_list = x)
})


# convert leading edge gene names from entrez to symbols
reactome_res_list <- lapply(reactome_res_list, function(x){
    x$leadingEdge_gene_symbol <- lapply(x$leadingEdge, function(y){
        working_conversion_df <- entrez_ID_df[entrez_ID_df$ENTREZID %in% y, ]
        working_conversion_df <- working_conversion_df[match(y, working_conversion_df$ENTREZID),]
        leadingEdge_genes_symbols <- working_conversion_df$SYMBOL
    })
    return(x)
})


# save results 
saveRDS(file = paste0(out_dir, '/', 'GSEA_reactome_results.rds'), reactome_res_list)


##############################################################################################################################
##############################################################################################################################
##############################################################################################################################
# plot upregualted pathways ----
lapply(seq_along(reactome_res_list), function(x){
    working_results <- reactome_res_list[[x]]
    working_results <- working_results %>% 
        filter(NES > 0) %>% 
        slice_max(order_by = NES, n = 10)

    plot <- ggplot(working_results, aes(reorder(pathway, NES), NES)) +
        geom_point(aes(fill = padj, size = size), shape=21) +
        scale_fill_gradient(low = "firebrick2", high = "dodgerblue2") +
        # scale_fill_manual(values = colors) +
        scale_size_continuous(range = c(3,8)) +
        # geom_hline(yintercept = 0) +
        coord_flip() +
        labs(x="Pathway", y="Normalized Enrichment Score",
            title= names(reactome_res_list)[[x]]) + 
        theme_bw()
    ggsave(file = paste0(out_dir, '/', 'GSEA_', names(reactome_res_list)[[x]], '_only_enriched.pdf'), width = 10, height = 7, plot, device = 'pdf')
})


##############################################################################################################################
##############################################################################################################################
##############################################################################################################################
# check expression of leading edge genes on UMAP

# load object 
obj <- readRDS(file = obj_file)


# plot
marker_dir <- paste0(out_dir, '/', 'reactome_leading_edge_genes')
dir.create(marker_dir, recursive = T)

working_module <- 'nmf_4066'
working_pathway <- 'REACTOME_CA2_PATHWAY'

working_genes <- reactome_res_list[[working_module]] %>% 
    filter(pathway == working_pathway) %>% 
    pull(leadingEdge_gene_symbol) %>% 
    unlist()

DefaultAssay(obj) <- 'RNA'
pdf(file = paste0(marker_dir, '/', working_module, '_', working_pathway, '.pdf'))
    for (g in working_genes) {
        print(
            FeaturePlot(obj, features = g, order = TRUE)
        )
    }
dev.off()


working_module <- 'nmf_4066'
working_pathway <- 'REACTOME_INOSITOL_PHOSPHATE_METABOLISM'

working_genes <- reactome_res_list[[working_module]] %>% 
    filter(pathway == working_pathway) %>% 
    pull(leadingEdge_gene_symbol) %>% 
    unlist()

DefaultAssay(obj) <- 'RNA'
pdf(file = paste0(marker_dir, '/', working_module, '_', working_pathway, '.pdf'))
    for (g in working_genes) {
        print(
            FeaturePlot(obj, features = g, order = TRUE)
        )
    }
dev.off()
```
