## Supplementary material for "snPATHO-seq: unlocking the pathology archives": scripts: 4066_top_DE_gene_heatmap.R

```
# aim ----
    # plot top DE gene heatmap from each gene module


# define environment ----
library(Seurat)
library(tidyverse)

library(ComplexHeatmap)
library(circlize)


# arguments ----
de_res_file <- '/share/ScratchGeneral/tonwan/projects/snPATHO-Seq/results/gene_module_analysis/4066_unique_GMs/DE_between_GM_cells.rds'

out_dir <- '/share/ScratchGeneral/tonwan/projects/snPATHO-Seq/results/gene_module_analysis/4066_unique_GMs'


# load data ----
de_res <- readRDS(file = de_res_file)


# get top DE genes ---
# select top genes separately because I want to make sure all genes are significant
top_genes <- de_res %>% 
    filter(p_val_adj < 0.05) %>% 
    group_by(cluster) %>% 
    slice_max(order_by = avg_log2FC, n = 20) %>% 
    pull(gene)

# get fold change of the top genes as plot matrix
de_res$cluster <- as.character(de_res$cluster)

exp <- de_res %>% 
    select(avg_log2FC, cluster, gene) %>% 
    filter(gene %in% top_genes) %>% 
    spread(value = 'avg_log2FC', key = 'cluster') %>% 
    column_to_rownames(var = 'gene') %>% 
    as.data.frame()


# define heatmap colors
col_fun <- colorRamp2(c(floor(min(exp)),
                        0,
                        ceiling(max(exp))), 
                      c("#2166AC", "white", "#B2182B"))


# plot heatmap ----
plot <- Heatmap(exp,
                name = 'Fold change - log2',
                heatmap_legend_param = list(title_gp = gpar(fontsize = 12, fontface = "bold")),
                col = col_fun,
                # right_annotation = right_annot,
                # bottom_annotation = bottom_annotation,
                show_column_names= TRUE,
                show_row_names = TRUE,
                cluster_rows = TRUE,
                cluster_columns = FALSE,
                column_names_gp = gpar(fontsize = 12),
                row_names_gp = gpar(fontsize = 12))

pdf(file = paste0(out_dir, '/', 'top_DE_genes_heatmap', '.pdf'),
    width = 5, height = 14)
    draw(plot)
dev.off()
```
