## Supplementary material for "snPATHO-seq: unlocking the pathology archives": scripts: 08_change_robust_NMF_names_for_paper.R

```
# aim ----
    # change program names of the robust NMF programs for publication


# define environment ----
library(tidyverse)


# arguments ----
robust_NMF_file <- '/share/ScratchGeneral/tonwan/projects/snPATHO-Seq/results/gene_module_analysis/robust_NMF_programs.rds'


# load data and change names ----
robust_NMF <- readRDS(file = robust_NMF_file)

robust_NMF <- as.matrix(robust_NMF)


colnames(robust_NMF) <- gsub('GEX', '_3p', colnames(robust_NMF))
colnames(robust_NMF) <- gsub('SNAPFix', '_FLEX', colnames(robust_NMF))
colnames(robust_NMF) <- gsub('FFPE_run1', '_FFPE', colnames(robust_NMF))
colnames(robust_NMF) <- gsub('4066FFPE', '4066_FFPE', colnames(robust_NMF))
colnames(robust_NMF) <- gsub('4411FFPE', '4411_FFPE', colnames(robust_NMF))


write.csv(file = paste0('/share/ScratchGeneral/tonwan/projects/snPATHO-Seq/results/gene_module_analysis',
    '/', 'robust_NMF_paper_format.csv'), robust_NMF)
```
