## Supplementary material for "snPATHO-seq: unlocking the pathology archives": scripts: cellbender_remove_background_GPU.sh

```
#!/bin/bash
#$ -N cellbender
#$ -pe smp 8
#$ -l mem_requested=8G
#$ -M
#$ -m ae
#$ -V
#$ -o /share/ScratchGeneral/tonwan/projects/snPATHO-Seq/logs/cellbender/$JOB_ID.out
#$ -e /share/ScratchGeneral/tonwan/projects/snPATHO-Seq/logs/cellbender/$JOB_ID.err
#$ -l nvgpu=1
#$ -l h=epsilon*
#$ -now no


# set directories
data_dir=/share/ScratchGeneral/tonwan/projects/snPATHO-Seq/data/external_data/FHIL_Data/Aligned/Kidney_whole_cell
input_file=$data_dir/outs/multi/count/raw_feature_bc_matrix.h5

out_dir=/share/ScratchGeneral/tonwan/projects/snPATHO-Seq/data/external_data/FHIL_Data/Aligned/Kidney_whole_cell/GPU
mkdir -p $out_dir
cd $out_dir


# run cellbender
cellbender remove-background \
     --cuda \
     --input $input_file \
     --output ./cellbender_filtered_matrix.h5 \
     --expected-cells 14312 \
     --total-droplets-included 40000
     
```
