## Supplementary material for "snPATHO-seq: unlocking the pathology archives": scripts: colors.R

```
# cell type colors
cell_type_colors <- c(
    '#d8391f', '#bd6d84', '#1a5e62',
    '#81ab7d', '#213e87', '#71ad2c', 
    '#bc6a23', '#e2c4aa', '#6A3D9A',
    '#B15928', '#EA469A', "#FB9A99",
    '#88181d', '#7eb2d0', '#fcc02d',
    '#d8391f',
    '#6A3D9A', '#FB9A99', '#992CA0',
    '#213e87', '#FD9C6F', '#8ADF8D')

names <- c(
    'Epithelial_cancer', 'CAF', 'Macrophage',
    'Endothelial', 'T_cells', 'B_cells', 
    'DC', 'MAST', 'Myoepithelial',
    'PVL', 'Epithelial_basal', 'Epithelial_luminal',
    'Epithelial_KRT7+', 'Plasma', 'NK',
    "Epithelial_KRT7-",
    'Cholangiocyte', 'Hepatocyte', 'LSEC',
    'Mixed_lymphocytes', 'RBCs', 'Lymphatic_endothelial')


names(cell_type_colors) <- names
rm(names)


# snRNA-seq workflow colors
workflow_colors <- c("#0072B2", "#009E73", "#E69F00", '#00a3ff', '#CC79A7', '#a5682a')
workflow_names <- c('snPATHO', 'FLEX', '3p', 'scFFPE', 'Gavish_et_al_2023', 'Spatial')
names(workflow_colors) <- workflow_names
rm(workflow_names)


# breast cancer sample colors
sample_colors <- c('#CC79A7', '#9e79cc', '#a7cc79', '#79cc9e')
sample_names <- c('Gavish_et_al_2023', '4411', '4066', '4399')
names(sample_colors) <- sample_names
rm(sample_names)
```
