## Supplementary material for "snPATHO-seq: unlocking the pathology archives": scripts: downsample_fastq.sh

```
#! /bin/bash
#$ -S /bin/bash
#$ -pe smp 8
#$ -cwd
#$ -l tmp_requested=40G
#$ -l tmpfree=40G

# This script downsamples a folder of compressed fastq files to the requested fraction or read count

module load aletan/seqtk/1.3

set -euxo pipefail

target=$1
out=$2
frac=$3

echo "Downsample rate: $frac"

if (( $(echo "$frac > 1" | bc -l) )) ; then
    mkdir -p $out
    cp -r $target $out
    exit
fi

# Copy files to temp
echo $TMPDIR
cd $TMPDIR
for file in $target/*
do
    echo "Copying $file"
    cp -r -L $file .
done

# Unzip
for file in $TMPDIR/*.fastq.gz
do
    echo "Unzipping $file"
    gunzip $file
done


# Down-sample with random seed=100
mkdir -p $out
for file in $TMPDIR/*.fastq
do
    echo "Downsampling $file"
    mkdir -p $out/$(dirname $file)
    seqtk sample -s100 $file $frac > $out/$file
    gzip $out/$file
done
```
