## Supplementary material for "snPATHO-seq: unlocking the pathology archives": scripts: submit_downsample.sh

```
#!/bin/bash

# where the fastq files are
data_dir="/share/ScratchGeneral/tonwan/projects/snPATHO-Seq/data/snPATHO_seq/Fastq_files/polyA/4399GEX/original"

# where the down sampled fastq files should be kept
out_dir="/share/ScratchGeneral/tonwan/projects/snPATHO-Seq/data/snPATHO_seq/downsampled_fastq_files/25k_reads_per_cell/4399GEX"
mkdir -p $out_dir

cd $out_dir

# function script
script_file=/share/ScratchGeneral/tonwan/projects/snPATHO-Seq/scripts/downsample_fastq/downsample_fastq.sh

# down sample rate
fraction=$(echo 25000 / 26062 | bc -l)

qsub $script_file $data_dir $out_dir $fraction
```
