## Supplementary material for "snPATHO-seq: unlocking the pathology archives": scripts: 01_Seurat_integration.R

```
# aim ----
    # function scripts to run Seurat integration methods
        # CCA and RPCA
        # using RNA or SCT assays as input


# requirement
    # data should have been filtered before / doublet removal


# arguments
    # obj_list                      a list of seurat objects to be integrated together
    # normalisation_method          method used for normalisation, 'RNA' or 'SCT'
    # nfeatue_each_dataset          numbers of features to be selected from each dataset
    # vars_to_regress               variable features to be regressed out during data scaling
    # integration_method            method used for integration, 'cca' or 'rpca'


# example
# integrated_obj <- seurat_integration(obj_list,
#                                     normalisation_method = 'RNA',
#                                     nfeatue_each_dataset = 2000,
#                                     nfeatures_common = 2000,
#                                     vars_to_regress = NULL,
#                                     integration_method = 'cca',
#                                     nthread = NULL)


seurat_integration <- function(
    obj_list,
    normalisation_method = 'RNA',
    nfeatue_each_dataset = 4000,
    nfeatures_common = 10000,
    vars_to_regress = NULL,
    integration_method = 'cca',
    nthread = NULL,
    ...
) {
    # make sure the following packages are loaded
    library(Seurat)

    # parallelisation
    if (!is.null(nthread)) {
        library(future)
        plan("multicore", workers = nthread)
    }
    
    # check normalisation method
    if (normalisation_method == 'RNA') {
        # normalise data & select features
        normalised_list <- lapply(obj_list, function(x){
            DefaultAssay(x) <- 'RNA'
            x <- NormalizeData(x)
            x <- FindVariableFeatures(x, selection.method = "vst", nfeatures = nfeatue_each_dataset)
        })

        # find common features for integration
        features <- SelectIntegrationFeatures(object.list = normalised_list, nfeatures = nfeatures_common)

        # scale data & PCA analysis
        # only scale the selected features to speed up data processing
        scaled_list <- lapply(normalised_list, function(x){
            x <- ScaleData(x, features = features, vars.to.regress = vars_to_regress, verbose = FALSE)
            x <- RunPCA(x, features = features, verbose = FALSE)
        })

        # integrate data
        anchors <- FindIntegrationAnchors(object.list = scaled_list, 
                anchor.features = features, 
                normalization.method = 'LogNormalize',
                reduction = integration_method)
        integrated_obj <- IntegrateData(anchorset = anchors,
            normalization.method = 'LogNormalize')
        
        # scale the integrated data
        integrated_obj <- ScaleData(integrated_obj, 
            features = features, 
            vars.to.regress = vars_to_regress, 
            verbose = FALSE)

    } else if (normalisation_method == 'SCT') {
        # SCT normalise data
        normalised_list <- lapply(obj_list, function(x){
            x <- SCTransform(x, assay = "RNA", 
                vst.flavor = "v2", 
                vars.to.regress = vars_to_regress,
                variable.features.n = nfeatue_each_dataset)
        })

        # find common features
        features <- SelectIntegrationFeatures(object.list = normalised_list, nfeatures = nfeatures_common)
        normalised_list <- PrepSCTIntegration(object.list = normalised_list, anchor.features = features)

        # run PCA on each object
        normalised_list <- lapply(normalised_list, function(x) {
            x <- RunPCA(x, features = features, verbose = FALSE)
        })

        # find anchors and integrate the data
        anchors <- FindIntegrationAnchors(normalised_list, 
            normalization.method = "SCT",
            anchor.features = features,
            reduction = integration_method)
        integrated_obj <- IntegrateData(anchors, normalization.method = "SCT")
    }

    # dimension reduction on integrated data
    integrated_obj <- integrated_obj %>% 
        RunPCA(verbose = FALSE) %>% 
        RunUMAP(reduction = "pca", dims = 1:30) %>% 
        FindNeighbors(reduction = "pca", dims = 1:30)
    
    return(integrated_obj)
}
```
