## Supplementary material for "snPATHO-seq: unlocking the pathology archives": scripts: 02_Seurat_clustering_optimisation.R

```
# aim ----
    # function script to cluster cells and make plots to facilitate the selection of optimal clustering resoltion


# arguments
    # obj                       # a seurat object with nearnest neighbour found
                                    # default assay set to the one clustering shoudl be conducted e.g. integrated
    # resolution_range          # range of lovian clustering resolution to be tested
    # plot_dir                  # directory where plots will be saved
    # batch_factor              # factor such as sample to check their contribtuion to each cluster
    # run_silhouette            # if silhouette score will be calculated or not
    # dist.matrix               # distance matrix for silhouette analysis


# example 
# obj <- clustering_resolution_optimisation(obj,
#                                         resolution_range = seq(0.2, 2.5, by = 0.1),
#                                         plot_dir = "./path",
#                                         batch_factor = "sample_id",
#                                         run_silhouette = TRUE,
#                                         dist.matrix = dist.matrix)


clustering_resolution_optimisation <- function(
    obj,
    resolution_range = seq(0.2, 2.5, by = 0.1),
    plot_dir,
    batch_factor = NULL,
    run_silhouette = FALSE,
    dist.matrix
) {
    ########################################################################################################
    ########################################################################################################
    # load essential packages
    library(Seurat)
    library(tidyverse)
    library(clustree)
    
    if (run_silhouette == TRUE) {
        library(cluster)
        library(factoextra)

        silhouette_df <- data.frame(Resolution=numeric(), Average_silhouette_width=numeric())
    }


    ########################################################################################################
    ########################################################################################################
    # cluster cells as selected resolution
    for (res in resolution_range) {
        obj <- FindClusters(obj, resolution = res)

        # get name of the cluster annotation 
        cluster_name <- grep(paste0('res.', res), colnames(obj@meta.data), value = T)[1]
        Idents(obj) <- cluster_name


        # plot umap to check clustering
        pdf(file = paste0(plot_dir, '/', 'umap_', res, '.pdf'))
            print(
                DimPlot(obj, reduction = "umap", group.by = cluster_name, label = T, repel = T)
            )
        dev.off()

        # split umap by clusters - in case there are too many overlappings
        pdf(file = paste0(plot_dir, '/', 'umap_split_by_clusters_', res, '.pdf'),
            height = 4 * (ceiling(length(unique(Idents(obj))) / 5)), width = 20)
            print(
                DimPlot(obj, reduction = "umap", group.by = cluster_name, 
                    split.by = cluster_name, label = T, repel = T, ncol = 5)
            )
        dev.off()


        # check contribution of another factor such as patient to clustering 
            # as an evaluation of integration quality and identifcation of sample specific feature
        # make a data frame to store silhouette info
        if (!is.null(batch_factor)) {
            cluster_vs_batch_table <- data.frame(table(obj@meta.data[[batch_factor]],
                                                        obj@meta.data[[cluster_name]]))
            colnames(cluster_vs_batch_table) <- c(batch_factor, 'Cluster_ID', 'Frequency')

            # get proportion
            total_cell_number_each_cell_type <- data.frame(table(obj@meta.data[[cluster_name]]))
            colnames(total_cell_number_each_cell_type) <- c('Cluster_ID', 'total_cell_number')
            cluster_vs_batch_table <- base::merge(cluster_vs_batch_table, total_cell_number_each_cell_type, 
                                                    by = 'Cluster_ID')
            cluster_vs_batch_table$proportion <- cluster_vs_batch_table$Frequency / cluster_vs_batch_table$total_cell_number

            # make a barplot using actual number 
            plot <- ggplot(cluster_vs_batch_table, 
                        aes(x = Cluster_ID,
                            y = Frequency,
                            fill = .data[[batch_factor]])) +
                        geom_col(position = "dodge") +
                        scale_fill_brewer(palette="Set3") + 
                        theme_bw() +
                        theme(axis.text.x = element_text(angle = 45, vjust = 1, hjust = 1))
            ggsave(file = paste0(plot_dir, '/', 'sample_contribution_', res, '.pdf'),
                plot, width = length(unique(cluster_vs_batch_table$Cluster_ID)) * 2,
                height = 7, limitsize = FALSE)
            
            # make a barplot to reflect proportion
            plot <- ggplot(cluster_vs_batch_table, 
                        aes(x = Cluster_ID,
                            y = proportion,
                            fill = .data[[batch_factor]])) +
                        geom_col() +
                        scale_fill_brewer(palette="Set3") + 
                        theme_bw() +
                        theme(axis.text.x = element_text(angle = 45, vjust = 1, hjust = 1))
            ggsave(file = paste0(plot_dir, '/', 'sample_contribution_proportion_', res, '.pdf'),
                plot, width = length(unique(cluster_vs_batch_table$Cluster_ID)) * 1,
                height = 7, limitsize = FALSE)
        }


        # calculate silhouette score at different clustering resolution 
        if (run_silhouette == TRUE) {
            clusters <- as.numeric(as.character(obj@meta.data[[cluster_name]]))
            sil <- silhouette(x = clusters, dist = dist.matrix)
            working_sil_df <- data.frame(sil)

            silhouette_df <- rbind(silhouette_df, 
                                    data.frame(Resolution=res,
                                                Average_silhouette_width=mean(working_sil_df$sil_width)))

            pdf(file = paste0(plot_dir, '/', 'Silhouette_plot_', res, '.pdf'),
                height = length(unique(clusters)) * 0.6)
                print(plot(sil))
            dev.off()
        }
    }


    ########################################################################################################
    ########################################################################################################
    # make a clustree plot
    prefix_name <- unique(unlist(lapply(str_split(grep('res', colnames(obj@meta.data), value = T), pattern = '\\.'), '[[', 1)))
    pdf(file = paste0(plot_dir, '/', 'clustree_plot.pdf'), 
        height = length(unique(resolution_range)) * 0.5, 
        width = 10)
        print(
            clustree(obj, prefix = paste0(prefix_name, '.'))
        )
    dev.off()

    # check silhouette score vs resoltuion
    if (run_silhouette == TRUE) {
        plot <- ggplot(silhouette_df, 
                    aes(x = Resolution,
                        y = Average_silhouette_width)) +
                    geom_point() +
                    theme_bw()
        ggsave(file = paste0(plot_dir, '/', 'Silhouette_vs_resolutions.pdf'),
            plot, width = nrow(silhouette_df) * 0.5,
            height = 7, limitsize = FALSE)
    }

    # return object with clustering info
    return(obj)
}
```
