## Supplementary material for "snPATHO-seq: unlocking the pathology archives": scripts: 03_doublet_finder.R

```
# aim ----
    # run doublet finder on seurat object to identify doublets
    # dimension reduction using basic parameters in Seurat
        # RNA assay
        # lognormalise data
        # 2000 variable features
        # 30 PCs


# requirement 
    # data should have gone through minimal filtering to improve the accuracy of doublet removal


# arguments
    # obj                               # seurat data object to be processed
    # expected_doublet_rate             # expected doublet rate
    # out_dir                           # directory to store diagnostic plots


# example
# filtered_obj <- DB_finder_removal(obj,
#                                 expected_doublet_rate = 0.8,
#                                 out_dir = '/dir',
#                                 output_col_name = 'doublet_status')


DB_finder_removal <- function(
    obj,
    expected_doublet_rate = 0.8,
    out_dir, 
    output_name,
    output_col_name = 'doublet_status'
) {
    # make sure the packages are loaded
    library(DoubletFinder)

    # dimension reduction using raw data
    DefaultAssay(obj) <- 'RNA'
    normalised_obj <- obj %>% 
        NormalizeData() %>% 
        FindVariableFeatures(selection.method = "vst", nfeatures = 2000)
    features <- VariableFeatures(normalised_obj)
    normalised_obj <- normalised_obj %>% 
        ScaleData(features = features, verbose = FALSE) %>% 
        RunPCA(features = features, verbose = FALSE) %>% 
        RunUMAP(reduction = "pca", dims = 1:30)
    
    # find pK
    sweep.res.list <- paramSweep_v3(normalised_obj, PCs = 1:30, sct = FALSE)
    sweep.stats <- summarizeSweep(sweep.res.list, GT = FALSE)
    bcmvn <- find.pK(sweep.stats)
    dev.off() # to exist the device used for autoplotting
    
    pdf(file = paste0(out_dir, '/', output_name, '_pk_optimisation.pdf'), width = 7, height = 7)
        print(
            barplot(bcmvn$BCmetric, names.arg = bcmvn$pK, las=2)
        )
    dev.off()

    working_pK <- as.numeric(as.character(bcmvn[bcmvn$BCmetric == max(bcmvn$BCmetric), 'pK']))

    # run doublet removal
    nExp <- round(ncol(normalised_obj) * expected_doublet_rate)
    processed_obj <- doubletFinder_v3(normalised_obj, pN = 0.25, pK = working_pK, nExp = nExp, PCs = 1:30)

    # check results
    col_to_plot <- grep('DF', colnames(processed_obj@meta.data), value = T)
    # default column name contain parameters used, make a new column name to make it easier for downstream analysis
    processed_obj@meta.data[[output_col_name]] <- ifelse(processed_obj@meta.data[[col_to_plot]] == 'Singlet', 'Singlet', 'Doublet')
    pdf(file = paste0(out_dir, '/', output_name, '_umap_doublet_classification.pdf'), width = 7, height = 7)
        print(DimPlot(processed_obj, reduction = "umap", group.by = col_to_plot))
    dev.off()

    # check doublet score
    col_to_plot <- grep('pANN', colnames(processed_obj@meta.data), value = T)
    pdf(file = paste0(out_dir, '/', output_name, '_umap_doublet_score.pdf'), width = 7, height = 7)
        print(FeaturePlot(processed_obj, reduction = "umap", features = col_to_plot))
    dev.off()

    # output object with
    return(processed_obj)
}
```
