## Supplementary material for "snPATHO-seq: unlocking the pathology archives": scripts: 04_prepare_data_for_complex_heatmap.R

```
# aim ----
    # summarise data for complexheatmap
    # doesn't include actual plotting as it can be more manual and customised
    # this is just to standardise the data


# arguments ----
    # seurat_obj                # data object containing all the annotations and expression
    # seurat_assay              # which assay to use
    # serat_slot                # which slot to get data from
    # genes_to_plot             # genes to use for plotting
    # grouping_annotations      # annotations used for grouping samples for mean calculation
    # additional_annotations    # additional annotations

prepare_data_for_heatmap <- function(
    seurat_obj,
    seurat_assay = 'RNA',
    serat_slot = 'data',
    genes_to_plot,
    grouping_annotations,
    additional_annotations,
    ...
    ) {
    library(Seurat)
    library(tidyverse)

    DefaultAssay(seurat_obj) <- seurat_assay
    genes_not_present <- setdiff(genes_to_plot, rownames(seurat_obj))
    print(paste0("The following genes are not present in the dataset ", genes_not_present))
    genes_to_plot <- genes_to_plot[genes_to_plot %in% rownames(seurat_obj)]

    mtx <- GetAssayData(seurat_obj, assay = seurat_assay, slot = serat_slot)[genes_to_plot, ]
    mtx <- as.data.frame(mtx)
    mtx$gene_name <- rownames(mtx)
    mtx_long <- gather(mtx, value = 'expression', key = 'ID', -gene_name)

    # prepare annotations
    if (is.null(additional_annotations)) {
        annot_df <- seurat_obj@meta.data[, c(grouping_annotations), drop = FALSE]
    } else {
        annot_df <- seurat_obj@meta.data[, c(grouping_annotations, additional_annotations)]
    }

    if (length(grouping_annotations) > 1) {
        annot_df$merged_grouping_annotation <- do.call(paste, c(annot_df[grouping_annotations], sep="_"))
    } else {
        annot_df$merged_grouping_annotation <- annot_df[[grouping_annotations]]
    }
    
    
    # add annotation to the gene expression dataframe
    annotated_mtx <- base::merge(mtx_long, annot_df[, 'merged_grouping_annotation', drop = FALSE], by.x = 'ID', by.y = 0)

    # get mean expression by group
    mean_mtx <- annotated_mtx %>% 
        group_by(merged_grouping_annotation, gene_name) %>% 
        summarise(mean(expression))
    plot_mtx <- mean_mtx[, c('mean(expression)', 'gene_name', 'merged_grouping_annotation')]
    plot_mtx <- as.data.frame(spread(plot_mtx, value = 'mean(expression)', key = 'merged_grouping_annotation'))
    plot_mtx <- as.matrix(column_to_rownames(plot_mtx, var = 'gene_name'))

    # get annotation dataframe
    rownames(annot_df) <- NULL
    annot_df <- distinct(annot_df)
    rownames(annot_df) <- annot_df$merged_grouping_annotation

    # make sure they have the same orders
    plot_mtx <- plot_mtx[, annot_df$merged_grouping_annotation]

    # save matrix and annotation into a list as output
    output_list <- list(plot_mtx, annot_df)
    return(output_list)
}


# function to assign colors to annotations
    # method 1: supply a brewer palette
    # method 2: supply a vector of colors
    # method 3: no color palette is big enough, generate a random palette 

# arguments ----
    # annot_df              # dataframe containing information to annotate
    # annot_colname         # name of the column to get colors
    # palette               # which brewer palette to use
    # colors                # will only be evaluated if no brewer palette is supplied. A vector of color names


set_colors <- function(
    annot_df,
    annot_colname,
    palette = NULL,
    colors = NULL,
    n = 10,
    seed = 123458
) {
    if (!is.null(palette)) {
        colors <- brewer.pal(length(unique(annot_df[[annot_colname]])), name = palette)
        names(colors) <- unique(annot_df[[annot_colname]])
        # in case there are < 3 annotations
        colors <- colors[1:length(unique(annot_df[[annot_colname]]))]
    } else if (!is.null(colors)) {
        colors <- colors
        names(colors) <- unique(annot_df[[annot_colname]])
    } else if (!is.null(n)) {
        if (!is.null(seed)) {
            set.seed(seed)
        }
        qual_col_pals = brewer.pal.info[brewer.pal.info$category == 'qual',]
        col_vector = unlist(mapply(brewer.pal, qual_col_pals$maxcolors, rownames(qual_col_pals)))
        colors <- sample(col_vector, n)
        names(colors) <- unique(annot_df[[annot_colname]])
    }
    
    return(colors)
}
```
