## Supplementary material for "snPATHO-seq: unlocking the pathology archives": scripts: 05_SCN_annotation.R

```
# aim ----
    # function to run SCN prediction


# requirement ----
    # TrainingDataPath        # source dataset to train SCN model
    # QueryDataPath           # target dataset to predict cell type label
    # OutputDir               # location where results should be saved
    # celltype                # cell type annotation data column on the training model


run_SCN <-function(
  training_data, 
  query_data, 
  out_dir, 
  annotation_col,
  common_genes,
  seed = 100,
  output_label
){
  library(singleCellNet)
  library(Seurat)

  # load training data
  training_sample_table <- training_data@meta.data
  training_sample_table <- droplevels(training_sample_table)
  training_sample_table$cell <- rownames(training_sample_table)
  exp_train <- GetAssayData(training_data, assay = 'RNA', slot = 'counts')

  # load query data
  query_sample_table <- query_data@meta.data
  query_sample_table <- droplevels(query_sample_table)
  query_sample_table$cell <- rownames(query_sample_table)
  exp_query <- GetAssayData(query_data, assay = 'RNA', slot = 'counts')

  # only keep intersecting genes
  true_common_genes <- intersect(common_genes, rownames(exp_train))
  true_common_genes <- intersect(true_common_genes, rownames(exp_query))

  # only keep overlapping genes
  exp_train <- exp_train[true_common_genes, ]
  exp_query <- exp_query[true_common_genes, ]


  # split training data into training set and validation set
  set.seed(seed)
  stList <- splitCommon(sampTab=training_sample_table, ncells=200, dLevel=annotation_col)
  training_subset <- stList[[1]]
  training_subset_exp <- exp_train[, rownames(training_subset)]

  validation_subset <- stList[[2]]
  validation_subset_exp <- exp_train[,rownames(validation_subset)]

  # train a classifier
  classifier_file <- paste0(out_dir, '/', output_label, '_classifier.rds')
  if (file.exists(classifier_file)) {
    classifier <- readRDS(file = classifier_file)
  } else {
    print('Training classifier')
    classifier <- scn_train(
      stTrain = training_subset, 
      expTrain = training_subset_exp, 
      nTopGenes = 50, 
      nRand = 70, 
      nTrees = 1000, 
      nTopGenePairs = 25, 
      dLevel = annotation_col, 
      colName_samp = "cell")
    
    # save classifier
    saveRDS(classifier, file = classifier_file)
  }
  


  # predict label on validation set
  print('Running validation test')
  validation_test <- scn_predict(classifier[['cnProc']], validation_subset_exp, nrand = 70)
  saveRDS(validation_test, paste0(out_dir, '/', output_label, "_validation_test.rds"))
  
  # check validation results
  heldoutassessment <- assess_comm(
    ct_scores = validation_test, 
    stTrain = training_subset, 
    stQuery = validation_subset, 
    dLevelSID = "cell", 
    classTrain = annotation_col, 
    classQuery = annotation_col,
    nRand = 70)
  
  # holdout test
  pdf(file = paste0(out_dir, '/', output_label, '_validation_holdout_assessment.pdf'),
    width = ceiling(sqrt(length(unique(training_data@meta.data[[annotation_col]])))) * 4,
    height = ceiling(sqrt(length(unique(training_data@meta.data[[annotation_col]])))) * 4)
        plot_PRs(heldoutassessment)
  dev.off()

  # result heatmap
  nrand = 70
  sla = as.vector(validation_subset[[annotation_col]])
  names(sla) = as.vector(validation_subset$cell)
  slaRand = rep("rand", nrand) 
  names(slaRand) = paste("rand_", 1:nrand, sep='')
  sla = append(sla, slaRand)

  pdf(file = paste0(out_dir, '/', output_label, '_validation_heatmap.pdf'),
    width = 14, height = 10)
    sc_hmClass(validation_test, grps = sla, maxPerGrp=300, fontsize_row=7, isBig=TRUE)
  dev.off()
  


  ###################### annotate cells in query data ######################
  # run SCN on query dataset
  print('predicting label on the query dataset')

  prediction <- scn_predict(classifier[['cnProc']], exp_query, nrand = 70)
  saveRDS(prediction, paste0(out_dir, '/', output_label, "_prediction_results.rds"))

  # add predicted label to metadata
  query_sample_table_predicted <- assign_cate(
    classRes = prediction[,1:nrow(query_sample_table)], 
    sampTab = query_sample_table, 
    cThresh = 0.5)

  # save modified metadata
  write.csv(file = paste0(out_dir, '/', output_label, "_meta_data_table.csv"),
    query_sample_table_predicted)
}
```
