## Supplementary material for "snPATHO-seq: unlocking the pathology archives": scripts: 06_fgsea_analysis.R

```
# aim ----
    # run fgsea analysis

# gmt_file          gmt file path
# gene_list         gene list order from high to low with names


# obtained from 
    # https://bioinformaticsbreakdown.com/how-to-gsea/
GSEA <- function(gmt_file, gene_list, collapse_pathways = TRUE) {
    library(fgsea)
    # load reference gmt files
    gs <- fgsea::gmtPathways(gmt_file)
    # run fgsea
    fgRes <- fgsea::fgsea(pathways = gs,
                           stats = gene_list,
                           minSize=15, ## minimum gene set size
                           maxSize=200, ## maximum gene set size
                           nperm=10000) 
    # filter results 
    fgRes_filtered <- fgRes %>% 
        as.data.frame() %>% 
        filter(padj < 0.05) %>% 
        arrange(desc(NES))

    # collapse similar pathways
    if (collapse_pathways == TRUE) {
        concise_pathways <- collapsePathways(data.table::as.data.table(fgRes_filtered),
                                      pathways = gs,
                                      stats = gene_list)  
        fgRes_filtered <- fgRes_filtered[fgRes_filtered$pathway %in% concise_pathways$mainPathways, ]
    }

    # return results 
    return(fgRes_filtered)
}
```
