## Supplementary material for "snPATHO-seq: unlocking the pathology archives": scripts: robust_nmf_programs.R

```
# -------------------------------------------------------------------------------------------
# Function for selecting robust nonnegative matrix factorization (NMF) programs
# ------------------------------------------------------------------------------------------- 

# - nmf_programs = a list; each element contains a matrix with NMF programs (top 50 genes) generated for a specific cell line using different NMF factorization ranks. 
# - intra_min = minimum overlap with a program from the same cell line (for selecting robust programs)
# - intra_max = maximum overlap with a program from the same cell line (for removing redundant programs)
# - inter_filter = logical; indicates whether programs should be filtered based on their similarity to programs of other cell lines
# - inter_min = minimum overlap with a program from another cell line 

# Returns a character vector with the NMF programs selected

robust_nmf_programs <- function(nmf_programs, intra_min = 35, intra_max = 10, inter_filter=T, inter_min = 10) {
  
  # Select NMF programs based on the minimum overlap with other NMF programs from the same cell line
  intra_intersect <- lapply(nmf_programs, function(z) apply(z, 2, function(x) apply(z, 2, function(y) length(intersect(x,y))))) 
  intra_intersect_max <- lapply(intra_intersect, function(x) apply(x, 2, function(y) sort(y, decreasing = T)[2]))             
  nmf_sel <- lapply(names(nmf_programs), function(x) nmf_programs[[x]][,intra_intersect_max[[x]]>=intra_min]) 
  names(nmf_sel) <- names(nmf_programs)
  
  # Select NMF programs based on i) the maximum overlap with other NMF programs from the same cell line and
  # ii) the minimum overlap with programs from another cell line
  nmf_sel_unlist <- do.call(cbind, nmf_sel)
  inter_intersect <- apply(nmf_sel_unlist , 2, function(x) apply(nmf_sel_unlist , 2, function(y) length(intersect(x,y)))) ## calculating intersection between all programs
  
  final_filter <- NULL 
  for(i in names(nmf_sel)) {
    a <- inter_intersect[grep(i, colnames(inter_intersect), invert = T),grep(i, colnames(inter_intersect))]
    b <- sort(apply(a, 2, max), decreasing = T) # for each cell line, ranks programs based on their maximum overlap with programs of other cell lines
    if(inter_filter==T) b <- b[b>=inter_min] # selects programs with a maximum intersection of at least 10
    if(length(b) > 1) {
      c <- names(b[1]) 
      for(y in 2:length(b)) {
        if(max(inter_intersect[c,names(b[y])]) <= intra_max) c <- c(c,names(b[y])) # selects programs iteratively from top-down. Only selects programs that have a intersection smaller than 10 with a previously selected programs
      }
      final_filter <- c(final_filter, c)
    } else {
      final_filter <- c(final_filter, names(b))
    }
  }
  return(final_filter)                                                      
}
```
