## Supplementary material for "snPATHO-seq: unlocking the pathology archives": scripts: plot_qc_metrics_for_all_data.R

```
# aim ----
    # plot QC metrics for all cellbender filtered data


# define environment ----
library(Seurat)
library(tidyverse)
library(RColorBrewer)

library(patchwork)
# library(ggpubr)

# arguments ----
out_dir <- '/share/ScratchGeneral/tonwan/projects/snPATHO-Seq/results/QC_plots'


################################################################################################################
################################################################################################################
################################################################################################################
# load all datasets with annotated cell types 
    # except for breast cancer scFFPE data - load cellranger outputs

summary_list <- list()


# breast cancer data - snRNA datasets
brca_data_dir <- '/share/ScratchGeneral/tonwan/projects/snPATHO-Seq/data/snPATHO_seq/25k_downsampled_processed_objs/integrated'

for (s in c('4066', '4399', '4411')) {
    obj <- readRDS(file = paste0(brca_data_dir, '/', s, '_cellbender_filtered_integration_final_annotation.rds'))
    df <- obj@meta.data[, c('nCount_RNA', 'nFeature_RNA', 'major_annotation', 'processing_method')]
    df$processing_method <- ifelse(df$processing_method == 'FFPE_snPATHO', 'snPATHO',
        ifelse(df$processing_method == 'SNAP_snPATHO', 'FLEX', '3p'))
    colnames(df) <- c('UMI', 'Gene', 'Annotation', 'Workflow')
    df$sample_id <- s
    df$processing_site <- 'Australia'
    summary_list[[s]] <- df
    rm(obj)
}


# BrCa scFFPE data - from cellranger outputs
cr_output_dir <- '/share/ScratchGeneral/tonwan/projects/snPATHO-Seq/data/snPATHO_seq/Aligned/FRP/scFFPE_multi_pool/outs/per_sample_outs'

for (s in c('4066', '4399', '4411')) {
    mtx <- Read10X_h5(paste0(cr_output_dir, '/', s, 'scFFPE/count/sample_filtered_feature_bc_matrix.h5'), 
        use.names = TRUE, unique.features = TRUE)
    obj <- CreateSeuratObject(mtx)
    df <- obj@meta.data[, c('nCount_RNA', 'nFeature_RNA')]
    colnames(df) <- c('UMI', 'Gene')
    df$Annotation <- 'unannotated'
    df$Workflow <- 'scFFPE'
    df$sample_id <- s
    df$processing_site <- 'Australia'
    summary_list[[paste0('scFFPE_', s)]] <- df
    rm(mtx)
    rm(obj)
}


# 10X data
annotated_10x_obj_dir <- '/share/ScratchGeneral/tonwan/projects/snPATHO-Seq/data/external_data/10X_data_annotated'
samples <- c('ColonCRC_1328A', 'EndocervicalAdenocarcinoma_6707', 'Glioblastoma_1773A',
    'Kidney_1305272B', 'liver_8754A', 'Lung_20011329NL', 'lungcancer_20011329LC',
    'melanoma_7167A')

for (s in samples) {
    obj <- readRDS(file = paste0(annotated_10x_obj_dir, '/', s, '_annotated.rds'))
    df <- obj@meta.data[, c('nCount_RNA', 'nFeature_RNA', 'initial_annotation', 'processing_method')]
    colnames(df) <- c('UMI', 'Gene', 'Annotation', 'Workflow')
    df$sample_id <- s
    df$processing_site <- '10X_Genomics'
    summary_list[[s]] <- df
    rm(obj)
}


# Lyon data 
lyon_obj_dir <- '/share/ScratchGeneral/tonwan/projects/snPATHO-Seq/data/external_data/CRCL-Data_annotated'
samples <- c('Colon_213641', 'Endometrium_220952', 'Ovary_230303')

for (s in samples) {
    obj <- readRDS(file = paste0(lyon_obj_dir, '/', s, '_annotated.rds'))
    df <- obj@meta.data[, c('nCount_RNA', 'nFeature_RNA', 'initial_annotation', 'processing_method')]
    colnames(df) <- c('UMI', 'Gene', 'Annotation', 'Workflow')
    df$sample_id <- s
    df$processing_site <- 'Lyon'
    summary_list[[s]] <- df
    rm(obj)
}


# FHIL data 
obj <- readRDS(file = '/share/ScratchGeneral/tonwan/projects/snPATHO-Seq/data/external_data/FHIL_Data/annotated/Kidney_FHIL_annotated.rds')
df <- obj@meta.data[, c('nCount_RNA', 'nFeature_RNA', 'initial_annotation', 'processing_method')]
colnames(df) <- c('UMI', 'Gene', 'Annotation', 'Workflow')
df$sample_id <- 'Kidney_K48'
df$processing_site <- 'Fred_Hutch'
summary_list[['Kidney_K48']] <- df
rm(obj)


# merge all data together into a single dataframe
summary_df <- do.call(rbind, summary_list)

# change name of mislabelled sample
    # this is a Crohn's disease sample not CRC
summary_df$sample_id <- ifelse(summary_df$sample_id == 'ColonCRC_1328A', 'Colon_1328A', summary_df$sample_id)


################################################################################################################
################################################################################################################
################################################################################################################
# plots

# load predefined colors 
source('/share/ScratchGeneral/tonwan/projects/snPATHO-Seq/scripts/colors.R')
workflow_colors <- workflow_colors[names(workflow_colors) %in% unique(summary_df$Workflow)]

# control plot order
summary_df$Workflow <- factor(summary_df$Workflow , levels = c('3p', 'FLEX', 'snPATHO', 'scFFPE'))

summary_df$workflows_tested <- ifelse(summary_df$processing_site == 'Australia', 'all',
    ifelse(summary_df$processing_site == 'Lyon', 'snPATHO_only', 'sn_scFFPE'))


# plot UMI per cell/nucleus
    # plot in sections and then patch them together
plot_list <- list()
for (s in c('all', 'sn_scFFPE', 'snPATHO_only')) {
    p <- ggplot(summary_df[summary_df$workflows_tested == s, ], 
        aes(x = sample_id, y = UMI, group = interaction(sample_id, Workflow), fill = Workflow)) +
        geom_boxplot(outlier.size = 0) +
        theme_classic() +
        scale_fill_manual(values = workflow_colors) +
        scale_y_continuous(
            limits = c(0, 15000), 
            breaks = c(0, 2500, 5000, 7500, 10000, 12500, 15000),
            expand = c(0,0)) +
        theme(
            axis.text.x = element_text(angle = 45, hjust = 1, vjust = 1),
            text = element_text(size = 16),
            axis.title.x = element_blank()
        ) +
        labs(fill = 'Workflows')
    plot_list[[s]] <- p
}


# make fine tuning
    # for the first plot on the left
        # allow y axis labels
plot_list[[1]] <- plot_list[[1]] +
            labs(y = 'Numbers of UMI detected per cell / nucleus')
    # for the 2nd plot
        # disable y axis related features
plot_list[[2]] <- plot_list[[2]] +
            theme(
                legend.position = 'none',
                axis.line.y = element_blank(),
                axis.text.y = element_blank(),
                axis.ticks.y = element_blank(),
                axis.title.y = element_blank())
    # for the 3rd plot
        # disable y axis related features
plot_list[[3]] <- plot_list[[3]] +
            theme(
                legend.position = 'none',
                axis.line.y = element_blank(),
                axis.text.y = element_blank(),
                axis.ticks.y = element_blank(),
                axis.title.y = element_blank())

# merge plots together
plot <- plot_list[[1]] +
        plot_list[[2]] + 
        plot_list[[3]] +  
        plot_layout(ncol = 3, widths = c(12, 18, 3), guides = "collect")

ggsave(file = paste0(out_dir, '/', 'QC_UMI_per_unit_paper_figure.pdf'), 
    width = 30, height = 10, plot, device = 'pdf')


# plot Gene per cell/nucleus
    # plot in sections and then patch them together
plot_list <- list()
for (s in c('all', 'sn_scFFPE', 'snPATHO_only')) {
    p <- ggplot(summary_df[summary_df$workflows_tested == s, ], 
        aes(x = sample_id, y = Gene, group = interaction(sample_id, Workflow), fill = Workflow)) +
        geom_boxplot(outlier.size = 0) +
        theme_classic() +
        scale_fill_manual(values = workflow_colors) +
        scale_y_continuous(
            limits = c(0, 9000), 
            breaks = c(0, 2000, 4000, 6000, 8000),
            expand = c(0,0)) +
        theme(
            axis.text.x = element_text(angle = 45, hjust = 1, vjust = 1),
            text = element_text(size = 16),
            axis.title.x = element_blank()
        ) +
        labs(fill = 'Workflows')
    plot_list[[s]] <- p
}


# make fine tuning
    # for the first plot on the left
        # allow y axis labels
plot_list[[1]] <- plot_list[[1]] +
            labs(y = 'Numbers of genes detected per cell / nucleus')
    # for the 2nd plot
        # disable y axis related features
plot_list[[2]] <- plot_list[[2]] +
            theme(
                legend.position = 'none',
                axis.line.y = element_blank(),
                axis.text.y = element_blank(),
                axis.ticks.y = element_blank(),
                axis.title.y = element_blank())
    # for the 3rd plot
        # disable y axis related features
plot_list[[3]] <- plot_list[[3]] +
            theme(
                legend.position = 'none',
                axis.line.y = element_blank(),
                axis.text.y = element_blank(),
                axis.ticks.y = element_blank(),
                axis.title.y = element_blank())

# merge plots together
plot <- plot_list[[1]] +
        plot_list[[2]] + 
        plot_list[[3]] +  
        plot_layout(ncol = 3, widths = c(12, 18, 3), guides = "collect")

ggsave(file = paste0(out_dir, '/', 'QC_Gene_per_unit_paper_figure.pdf'), 
    width = 30, height = 10, plot, device = 'pdf')


# plot cell type proportions
    # summarise to only plot 
        # epithelial / malignant
        # stromal
        # immune
        # Other

summary_df$modified_cell_types <- ifelse(
    summary_df$Annotation %in% c(
        'Epithelial_cancer', 'Myoepithelial', 'Epithelial_basal', 'Epithelial_luminal', 'Cholangiocyte',
        'Epithelial', 'Distal_convoluted_tubule', 'Proximal_tubule', 'Loop_of_Henle_epithelial',
        'Collecting_tubule_and_connecting_tubule', 'Intercalated_type_A', 'Parietal_epithelial',
        'Intercalated_type_B', 'Podocytes', 'Cholangiocytes', 'Hepatocyte', 'Hepatocytes', 'AT2',
        'Ciliated_cells', 'AT1', 'Neuroendocrine_cells', 'Epithelial_cycling', 'Keratinocytes',
        'Melanocytes', 'Melanocytes_cycling', 'Melanocytes_heatshock_high', 'Tuft_cells', 'Intercalated_cells'), 'Epithelial / Malignant',
        ifelse(summary_df$Annotation %in% c(
            'CAF', 'Endothelial', 'PVL', 'Lymphatic_endothelial', 'LSEC', 'Fibroblasts', 'Smooth_muscle_cells',
            'Stellate_cells', 'Pericytes', 'SMCs', 'Portal_fibroblasts', 'vSMC', 'Other_endothelial',
            'CAF_PVL', 'SMC', 'Fibroblast_pericyte'), 'Stromal',
            ifelse(summary_df$Annotation %in% c(
                'MAST', 'Macrophage', 'DC', 'T_cells', 'B_cells', 'Mixed_lymphocytes', 'Plasmablasts', 'T_cycling',
                'Monocyte', 'B', 'T_CD4', 'T_CD8', 'Macrophages', 'Monocytes', 'pDC', 'T', 'Mixed_immune_cells',
                'NK', 'Other_macrophages', 'Kupffer_cells', 'NKT', 'Mast_cells', 'Myeloid', 'Lymphocytes',
                'Immune_cells', 'Monocyte_macrophage', 'Myeloid_cycling', 'Plasmablast', 'Dendritic_cell', 'T_reg'), 'Immune', 
                ifelse(summary_df$Annotation %in% c(
                    'Neuron', 'Oligodendrocyte_progenitor_cells', 'NPC_like', 'OPC_like', 'AC_like',
                    'MES_like', 'Schwann_cells'), 'Neuron', 'RBC / Unlabelled'
                )
            )
        )
    )


# define colors for the cell types
annotation_colors <- c("#D55E00", "#E69F00", "#0072B2", "#0072B2", "#999999")
names(annotation_colors) <- c('Epithelial / Malignant', 'Stromal', 'Immune', 'Neuron', "RBC / Unlabelled")

plot_df <- summary_df %>% 
    ungroup() %>% 
    group_by(modified_cell_types, Workflow, sample_id, workflows_tested) %>% 
    summarise(summarised_cell_numbers = n()) %>% 
    as.data.frame()

plot_df$modified_cell_types <- factor(plot_df$modified_cell_types, 
    levels =c("Epithelial / Malignant", "Stromal", "Immune", "Neuron" , "RBC / Unlabelled"))


# plot cell type proportion per sample
    # plot in sections and then patch them together
plot_list <- list()
for (s in c('all', 'sn_scFFPE', 'snPATHO_only')) {
    p <- ggplot(plot_df[plot_df$workflows_tested == s, ], 
        aes(x = Workflow, y = summarised_cell_numbers, fill = modified_cell_types)) +
        geom_col() +
        theme_classic() +
        scale_fill_manual(values = annotation_colors) +
        facet_grid(cols = vars(sample_id)) +
        scale_y_continuous(
            limits = c(0, 18000), 
            breaks =c(0, 3000, 6000, 9000, 12000, 15000, 18000),
            expand = c(0,0)) +
        theme(
            axis.text.x = element_text(angle = 45, hjust = 1, vjust = 1),
            text = element_text(size = 16),
            axis.title.x = element_blank(),
            strip.background = element_blank(),
            strip.text.x = element_text(size = 7, face = "bold")
        ) +
        labs(fill = 'Major cell lineages')
    plot_list[[s]] <- p
}

# make fine tuning
    # for the first plot on the left
        # allow y axis labels
plot_list[[1]] <- plot_list[[1]] +
            labs(y = 'Numbers of cells / nuclei')
    # for the 2nd plot
        # disable y axis related features
plot_list[[2]] <- plot_list[[2]] +
            theme(
                legend.position = 'none',
                axis.line.y = element_blank(),
                axis.text.y = element_blank(),
                axis.ticks.y = element_blank(),
                axis.title.y = element_blank())
    # for the 3rd plot
        # disable y axis related features
plot_list[[3]] <- plot_list[[3]] +
            theme(
                legend.position = 'none',
                axis.line.y = element_blank(),
                axis.text.y = element_blank(),
                axis.ticks.y = element_blank(),
                axis.title.y = element_blank())

# merge plots together
plot <- plot_list[[1]] +
        plot_list[[2]] + 
        plot_list[[3]] +  
        plot_layout(ncol = 3, widths = c(11, 18, 3), guides = "collect")

ggsave(file = paste0(out_dir, '/', 'QC_cells_per_sample_paper_figure.pdf'), 
    width = 30, height = 10, plot, device = 'pdf')
```
